# Higher fungal diversity is correlated with lower CO_2_ emissions from dead wood in a natural forest: BioRxiv preprint

**DOI:** 10.1101/051235

**Authors:** Chunyan Yang, Douglas A. Schaefer, Weijie Liu, Viorel D. Popescu, Chenxue Yang, Xiaoyang Wang, Chunying Wu, Douglas W. Yu

**Affiliations:** State Key Laboratory of Genetic Resources and Evolution, Kunming Institute of Zoology, Chinese Academy of Sciences, 32 Jiaochang East Rd., Kunming, Yunnan 650223 China; Key Laboratory of Tropical Forest Ecology, Chinese Academy of Sciences, Xishuangbanna Tropical Botanical Garden, Menglun, Mengla, Yunnan 666303, China; Earth to Ocean Research Group, Department of Biological Sciences, Simon Fraser University, Burnaby, British Columbia V5A1S6, Canada; School of Biological Sciences, University of East Anglia, Norwich Research Park, Norwich, Norfolk NR47TJ UK

## Abstract

Wood decomposition releases almost as much CO_2_ to the atmosphere as does fossil-fuel combustion, so the factors regulating wood decomposition can affect global carbon cycling. We used metabarcoding to estimate the fungal species diversities of naturally colonized decomposing wood in subtropical China and, for the first time, compared them to concurrent measures of CO_2_ emissions. Wood hosting more diverse fungal communities emitted less CO_2_, with Shannon diversity explaining 26 to 44% of emissions variation. Community analysis supports a ‘pure diversity’ effect of fungi on decomposition rates and thus suggests that interference competition is an underlying mechanism. Our findings extend the results of published experiments using low-diversity, laboratory-inoculated wood to a high-diversity, natural system. We hypothesize that high levels of saprotrophic fungal biodiversity could be providing globally important ecosystem services by maintaining dead-wood habitats and by slowing the atmospheric contribution of CO_2_ from the world’s stock of decomposing wood. However, large-scale surveys and controlled experimental tests in natural settings will be needed to test this hypothesis.

## Introduction

Global decomposition of wood releases CO_2_ (6 to 9.5 Pg C/year^1, 2, 3^) at similar rates to fossil-fuel combustion (9.5 Pg C/year in 2011^4^). Decomposing wood also serves as essential habitat^5, 6^. The factors controlling wood decomposition rates are therefore of broad importance to conservation and to carbon cycle-climate feedbacks.

However, temperature and moisture variables only explain minority portions of total variance in decomposition rates^7, 8^. For instance, Bradford *et al.*^9^ reported that regional temperatures explain only 28% of local variance in mass loss.

The diversity of wood-decomposing fungi might explain much of the remaining unexplained variance. In laboratory-inoculation experiments using small numbers of culturable fungal species, wood pieces with higher final fungal diversity exhibited reduced decay rates^10, 11, 12^. Inoculated wood placed in the field also showed a negative effect of final fungal species diversity on decay (R^2^ = 0.15^13^).

However, in contrast to laboratory experiments, natural wood decomposition involves much higher species diversity, more complex assembly histories, and selective faunal feeding on decomposers^14, 15, 16^. Thus, it is important to examine the relationship between fungal diversity and decomposition rates in wood that is colonized and decomposing under natural conditions.

Natural fungal communities can be characterized using metabarcoding^17^, in which nuclear ribosomal internal transcribed spacer (ITS) regions are PCR-amplified and read using high-throughput sequencing^18, 19, 20, 21^. ITS1 and ITS2 are each sufficiently variable to differentiate fungal species^18, 19^ and return similar estimates of OTU (Operational Taxonomic Units) richness and community structure^19, 22^.

Here we metabarcoded ITS2 to examine fungal communities in naturally colonized wood pieces sampled across a wide range of decay classes in the Ailao Mountain forest of Yunnan, China. These wood pieces were sampled from a larger experiment involving three tree species (LC: *Lithocarpus chintungensis* [Fagaceae], LX: *L. xylocarpus*, and SN: *Schima noronhae* [Theaceae]) from which naturally occurring dead-wood pieces were regularly measured for CO_2_ emission rates over three years^8^. We measured the extent to which variation in the species diversity and composition of fungal communities can explain variation in emission rates.

## Results

*Taxonomy results* — Numbers of fungal OTUs ranged from 17 to 199 across wood pieces, tree species, and sampling dates, with means of 73.8 (LC, Sep 2012), 76.7 (LC, June 2013), 87.0 (LX, Sep 2012), 90.5 (LX, June 2013), 83.7 (SN, Sep 2012), and 88.4 (SN, June 2013).

41.1% of the 1,807 OTUs produced by *uclust* and 76.3% of the 1,565 OTUs produced by CROP were assigned to Fungi, and the proportions assigned to each fungal class were similar across assignment methods (Table 1). Because we removed non-Fungi reads from the dataset before taxonomic assignment, we attribute the taxonomically unassigned OTUs to the still highly incomplete UNITE and Genbank databases used for taxonomic assignment.

*Fungal diversity and CO_2_ emissions* — In June 2013, the month with the highest CO_2_ emissions, emissions declined with fungal diversity in all wood species (R^2^ = 26% to 44%, Fig. 1C, F).

**Figure 1.**
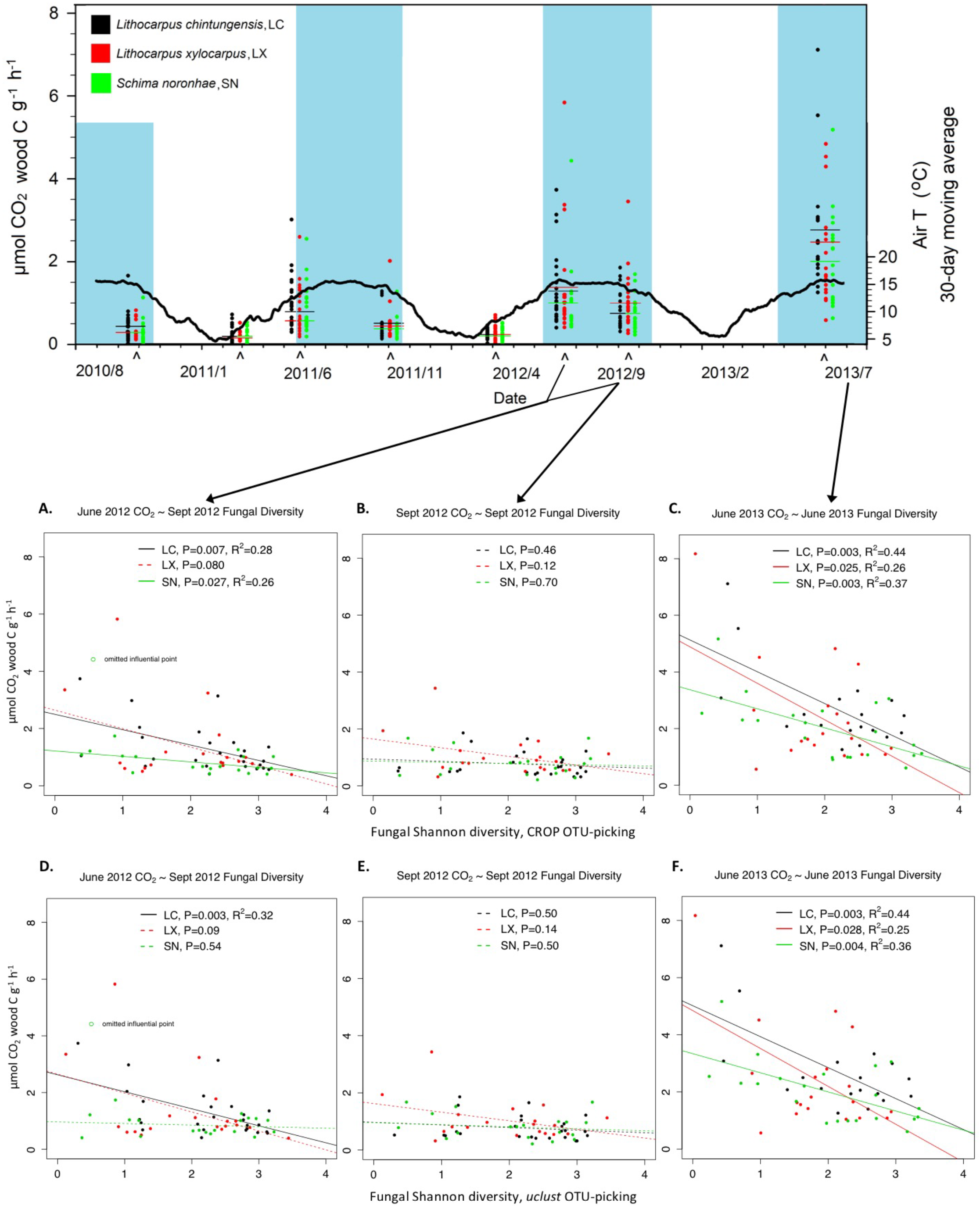
Linear regressions of CO_2_ emission rates on Shannon fungal diversities measured from individually metabarcoded wood pieces. **Top**. The solid black curve indicates the air temperature. Carets indicate times of CO_2_ measurements. Blue shading indicates the warm months when wood decomposition is >50% of maximum. **A-C**. CO_2_ emissions decline with increased fungal species diversity in two of the species in June 2012 (LC and SN) and in all three species in June 2013. In September 2012, CO_2_ emissions are lower, and there is no relationship. The OTU-picking method is *de novo* clustering with CROP. **D-F**. Same as A-C but the OTU-picking method is QIIME’s reference-based matching against the UNITE database, with *de novo* clustering of non-matched reads with *uclust*. Non-significant regressions are indicated by dashed lines. Shown here are the non-rarefied datasets. Rarefaction does not change the results (Supporting Information S1). LC = *Lithocarpus chintungensis*, LX = *L. xylocarpus*, SN = *Schima noronhae*.

**Table 1.** Taxonomic assignments to Class level for the ITS2 Operational Taxonomic Units (OTUs). *uclust* and CROP refer to the two OTU-clustering methods used, and GenBank and UNITE refer to the fungal reference databases used (see Methods: *Bioinformatic Analyses* for details).

| CROP + Genbank (1,565 OTUs) |  |  |  |  |  |  |  |
| --- | --- | --- | --- | --- | --- | --- | --- |
| Kingdom Fungi | Ascomycota | Basidiomycota | Chytridiomycota | Zygomycota | Glomeromycota | Mortierellales | Phylum unidentified |
| 1194 OTUs | 632 | 319 | 1 | 3 | 1 | 3 | 235 |
| 76.3% of total | 52.9% of Fungi | 26.70% | 0.10% | 0.30% | 0.10% | 0.30% | 19.70% |
| *uclust* + UNITE (1,806 OTUs) |  |  |  |  |  |  |  |
| Kingdom Fungi | Ascomycota | Basidiomycota | Chytridiomycota | Zygomycota | Phylum unidentified |  |  |
| 742 OTUs | 378 | 169 | 2 | 5 | 188 |  |  |
| 41.1% of total | 50.9% of Fungi | 22.80% | 0.30% | 0.70% | 25.30% |  |  |

In June 2012, CO_2_ emissions from LC and SN also declined with fungal diversity, even though we used a fungal diversity estimate taken three months later (September 2012) and even after conservatively omitting an influential datum from SN (high CO_2_, low diversity) (Fig. 1A, D). The third species (LX) did not return a significant regression, but its CO2-diversity relationship was visually nearly indistinguishable from its congener LC, suggesting that wood species partly governs the emissions-diversity relationship. Variances explained (26% to 28%, Fig. 1A, D) were lower than in June 2013.

Finally, in September 2012, CO_2_ emissions did not decline with higher fungal diversity (Fig. 1B, E), which is consistent with the generally lower CO_2_ emissions in September (Fig. 1).

The above results were robust to two OTU-picking methods (CROP and *uclust*, Fig. 1), rarefaction (non-rarefied shown in Fig. 1; rarefied in Supporting Information S1), and two diversity estimates (Shannon in Fig. 1, Simpson in S1). Regressions using Simpson diversities were generally statistically *more* significant (S1). We also analyzed after omitting single-read OTUs (which are more likely to be pipeline artefacts^23^) and achieved the same results, except that the previously non-significant SN regressions in Sept 2012 (Fig. 1A, D) became statistically *significant* (authors’ unpublished results). In short, the analyses presented in Fig. 1 are conservative.

*Chemistry of decomposing wood* — We analyzed the chemistry of a separate subset of 27 wood pieces from the larger experiment. Mean CO_2_ emissions from 2010-2012 showed no correlation with the densities of carbon, nitrogen, phosphorus, or lignin within any of the decay classes, with one exception, nitrogen density in decay class 1 (statistical details in Supporting Information S2).

*Pure-diversity versus species-selection effects* — Two general mechanisms could explain the observed diversity-function relationships. The first is a ‘pure diversity’ effect where species identity does not matter, only that increased species richness and evenness *per se* is somehow responsible for slower wood decomposition. The second is a ‘species-selection’ effect where more diverse fungal communities might be more likely to contain particular species that cause slow decomposition and somehow also govern the overall decomposition rate of the wood piece. To differentiate these two, we used a method devised by Sandau *et al.* ^24^ to generate a parameter λ for each regression in Fig. 1 (statistical details in Supporting Information S3). λ ranges between 0 and 1, with 0 indicating that variation in species composition does not account for variation in CO_2_ emissions (i.e. a ‘pure-diversity’ effect). For two tree species, LC and LX, λ always took values near zero (Table 2). For the third tree species SN, λ was also nearly zero in June 2012 but took intermediate values in June 2013, suggesting that fungal composition in this tree species at this time had some explanatory power. The general failure to detect composition effects can be observed in the community ordinations (Fig. 2) by noting that the SN/June 2013 samples were the only ones to line up along the CO_2_ emissions gradient (except the two lowest diversity samples). Not surprisingly, conventional community-analysis tests returned the same conclusion: variation in community composition is not explained by CO_2_ emissions (statistical details in Supporting Information S4).

**Figure 2.**
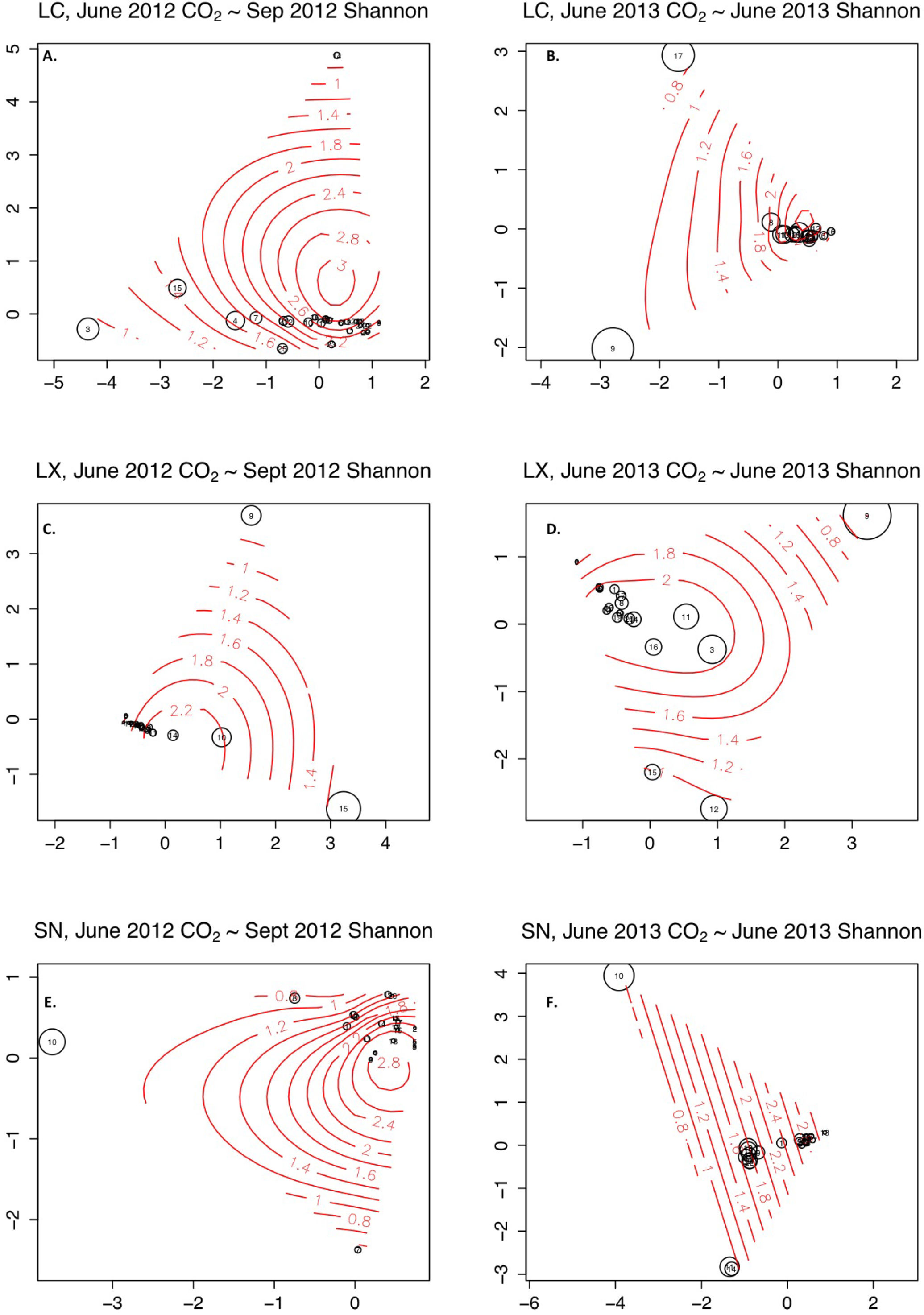
Correspondence analysis ordinations of fungal communities, by tree species and sampling date. Point size is scaled to CO_2_ emissions, and the gradient represents fungal Shannon diversity. In all ordinations (**A-F**), CO_2_ emissions decrease with higher fungal diversity (point size decreases up the gradient, echoing Fig. 1). Also evident is that the lower diversity wood pieces are compositionally very dissimilar to each other and to the higher diversity wood pieces. **Left-hand column (A, C, E)**. June 2012 CO_2_ vs. September 2012 fungal diversity. **Right-hand column (B, D, F)**. June 2013 CO_2_ vs. June 2013 fungal diversity. **A-B**. *Lithocarpus chintungensis*. Note that the label for point 14 at the top of A is obscured by the small point size. **C-D**. *L. xylocarpus*. **E-F**. *Schima noronhae*. Shown here are the non-rarefied datasets clustered using CROP (see Methods). Rarefaction or using *uclust*-clustering does not change the results (Supporting Information S4).

**Table 2.** Estimates of the contribution of fungal community composition to CO_2_ emissions, using the method of Sandau *et al.*^24^. The generated parameter λ varies between 0 and 1, with 0 indicating that variation in composition does not explain variation in emissions. Composition only contributes to explaining variation in one tree species in one sampling date (SN, June 2013). Conventional community analysis (Supporting Information S4) also detected a contribution of composition in SN in June 2013. LC = *Lithocarpus chintungensis*, LX = *L. xylocarpus*, SN = *Schima noronhae*.

| CO<sub>2</sub><br>sample | Species | OTU<br>clustering | $\lambda$ |
| --- | --- | --- | --- |
| Jun-12 | LC | CROP | 0.000 |
|  |  | *uclust* | 0.000 |
|  | LX | CROP | 0.002 |
|  |  | *uclust* | 0.000 |
|  | SN | CROP | 0.001 |
|  |  | *uclust* | 0.004 |
| Jun-13 | LC | CROP | 0.000 |
|  |  | *uclust* | 0.000 |
|  | LX | CROP | 0.000 |
|  |  | *uclust* | 0.001 |
|  | SN | CROP | 0.366 |
|  |  | *uclust* | 0.500 |

## Discussion

We found that naturally colonised wood with more diverse fungal communities decomposes more slowly (Fig. 1), resulting in a negative relationship between fungal biodiversity and the ecosystem function of decomposition. This result suggests positive relationships between fungal biodiversity and the ecosystem services of carbon storage and the provision of decomposing-wood habitat in forests.

Our results are consistent with five published experiments using laboratory-inoculated wood, which have all found negative relationships between fungal diversity and decomposition rates^10, 11, 12, 13, 25^. The one exception, Valentin *et al.*^26^, found a positive relationship, but in that study, field-collected microbial communities were serially diluted and re-inoculated into laboratory wood incubations. Serial dilution does not necessarily remove microbial species, but it does make all species less abundant^27^, which might have reduced decomposition rates.

In contrast, field studies to date have reported only ambiguous relationships between fungal diversity and decomposition rates. For instance, Hoppe *et al*.^28^ found non-significant correlations between fungal OTU richness and decay class (reflecting different numbers of years of decomposition), albeit negative relationships consistent with our results. Van Der Wal *et al.*^29^ measured tree-stump decomposition and reported weakly *positive* effects of fungal species richness (but not Simpson diversity) on sapwood decomposition, but only in late decay. Kurbartová *et al.^30^* found no relationship between wood loss and fungal OTU diversity after 12 years of decomposition but also reported that the least-decayed logs had the highest community diversities, again consistent with our results. All three studies found differences in community composition for logs that differed in remaining undecomposed weights.

Importantly, none of those three field studies made concurrent measurements of fungal diversities and CO_2_ emission rates, as we did here (Fig. 1). We observe that relationships between CO_2_ emissions rate and fungal diversity varied from month to month and across tree species (Fig. 1, Table 2, S1), suggesting that fungal activity and composition are dynamic and environmentally responsive. Thus, the fungal community measured after years of decomposition might not reflect the communities that were active during decomposition, obscuring any relationship between mass loss and fungal diversity.

Our study helps to reconcile the differing results found in the published laboratory and field studies, by making concurrent measurements of emissions and fungal diversity in a field setting that is the most natural on the spectrum of possibilities: colonization of locally dominant tree species by the local fungal community, with uninterrupted and full exposure to local environmental variability and the local faunal community, including fungivores, and long-term succession of fungal and other microbial communities. Our collected wood pieces span a range of at least one to fifteen years of decomposition on the forest floor^8^. Our results suggest that laboratory experiments correctly reveal negative relationships between CO_2_ emissions and fungal diversity.

*Mechanisms*. – In contrast to wood, microbial diversity is reported to accelerate the decomposition of soil organic matter^31, 32^, and this is thought to represent a general pattern^33, 34^ (but see Creed *et al.*^35^ for leaf litter). We hypothesise that because soil organic matter presents a much higher diversity of resources than does dead wood, niche complementarity amongst decomposer species drives positive relationships between diversity and soil organic matter decomposition.

By contrast, niche overlap provides a plausible biological mechanism for why wood decomposition should *slow* with fungal diversity. Interference competition has long been predicted to evolve when niche overlap is high and the disputed resource is valuable^36^. In forests, decomposing wood resources are available to many fungal species, and aggressive interactions are indeed observed among these fungi^37^. Elsewhere, it has been shown that interference competition reduces virulence (= host consumption rate) in endosymbioses^38, 39, 40^ and productivity in bacterial communities^41, 42^. Consistent with those findings in other contexts, interference competition can also explain why fungal biomass has been found to explain variance in wood mass loss^9^. When a piece of wood is colonized by many fungal species, the hypothesized higher levels of interference competition would result in less wood converted into fungal biomass (or CO_2_). High niche overlap is also consistent with the observed ‘pure diversity’ effect of fungal diversity on emissions (Table 2, S3, S4) since any species should fight all others (note, however, that we could not test for community composition effects at higher taxonomic levels, see Maherali & Klironomos^43^). Thus, theory suggests that the arrow of causation can be drawn in the direction of fungal diversity driving decomposition rate. Finally, interference competition results in competitive exclusion, which will cause community composition to change over time. This means that measures of fungal diversity made after years of decomposition are unlikely to explain final variance in decomposition.

*Sources of error and proposed future experiments*. – One source of error is that metabarcoding provides only approximate estimates of species frequencies, due to the many errors known to be introduced by metabarcoding, especially PCR-primer mismatches that lead to biased amplification^22, 23^. Also, DNA is environmentally persistent, so fungal species that are no longer represented by living colonies might still be detected by PCR. Nonetheless, we found that Shannon and Simpson indices, which both incorporate species frequency information by discounting rare species (here, rare=low-read OTUs), were able to explain variation in CO_2_ emissions. There are two likely and non-exclusive explanations. (1) Low-read OTUs were more likely to have been the remains of dead species and/or sequence artifacts from the metabarcoding pipeline and thus indeed should be discounted, and (2) in a Norway spruce forest, Ovaskainen *et al.*^21^ found that abundances of fungal fruiting bodies and OTU read numbers were positively correlated, suggesting that low-read OTUs indeed represent low-biomass species, which should have weaker influence on decomposition rates.

Another possible source of error is that we did not experimentally control for the age of the wood pieces, and thus an alternative explanation is that the observed correlations between fungal diversity and CO_2_ emissions rates (Fig. 1) might be caused by sampling along a successional gradient in which older wood pieces have less remaining wood to decompose (and thus lower emissions) and have also accumulated more fungal species. However, we found no relationship between decay class and emissions rates (*Methods: Experimental setup* and *Statistical analyses*) in our dataset, nor did we in the 320-piece superset from which our samples were drawn^8^, whereas this alternative explanation predicts that the least-decayed wood pieces should show the highest emissions. Also, we found mostly ‘pure-diversity’ effects of fungal communities on emissions (Table 2, S3, S4), whereas this alternative explanation invokes a successional sere and thus predicts compositional effects. We suggest a long-term experiment in which even-aged and sterilized wood pieces are allowed to be colonized and sampled for CO_2_ emissions rates and fungal diversity over many years in the field. We suggest that bacterial communities also be measured for correlations with CO_2_ emissions, although we caution that to estimate alpha diversity, sequencing effort will need to be much higher than for fungal communities. In addition to DNA-based metabarcoding, it might also be informative to sequence reverse-transcribed RNA from wood samples, in order to isolate the effect of living fungal species.

*Conclusions*. – The slopes of our diversity-emissions relationships (Fig. 1) are steep enough to suggest that even modest declines in fungal diversity in dead wood could cause several-fold increases in CO_2_ emissions rates. For example, in June 2013, CO_2_ emissions varied by 5.6-and 14.4-fold among LC and LX wood pieces. These negative relationships between diversity and wood decomposition provide a strong justification to conduct large-scale surveys of the status of fungal biodiversity and its trajectories in the world’s forests. Global forest fragmentation, reduction of tree-species diversity by fires, logging and replanting, the removal of dead trees, and even increased rainfall could all reduce fungal biodiversity in forests^6, 44^. All these changes could lead to faster wood decomposition. On the other hand, fragmented forests are drier, fungal distributions are being globalized^45^, and their local diversity is increased by rising CO_2_ ^46^. The net effect of all these changes on carbon emissions from the world’s stock of decomposing wood is difficult to predict.

## Methods

*Site description*. — This study was conducted in the Ailao Mountains National Nature Reserve, Yunnan, China, which preserves the largest area of undisturbed, subtropical moist forest in China and has a substantial pool of woody debris (branches and logs, 74.9 × 10^3^ kg ha^-1^, Ref. 47). The study site was at an elevation of 2476 m, about 2 km north of the Ailao Field Station for Forest Ecosystem Studies (24.533 °N, 101.017 °E), and receives 1840 mm annual average precipitation. The climate is monsoonal with distinct cool/dry (November to April) and warm/wet (May to October) seasons^48^. Annual mean air temperature is 11.3 °C with monthly means ranging from 5 to 16 °C. Surface soils (0 to 10 cm) of the area are Alfisols with pH of 4.2 (in water). The surficial organic layer is 3 to 7 cm deep^49^. The study site is a broad-leaved evergreen subtropical forest, with the canopy dominated by *Lithocarpus chintungensis, Rhododendron leptothrium, Vaccinium ducluoxii, Lithocarpus xylocarpus, Castanopsis wattii, Schima noronhae, Hartia sinensis*, and *Manglietia insignsis*^50^.

*Experimental setup*. — At our site, most woody debris comes from *Lithocarpus chintungensis*, (LC), *Lithocarpus xylocarpus* (LX), and *Schima noronhae* (SN), so we only examined those three species. In early 2010, branches from these three species, already decomposing on the forest floor, were identified to species by a botanist from the Xishuangbanna Tropical Botanical Garden, collected, and cut into a total of 320 wood pieces, sized to fit a field-respiration chamber (*ca*. 10 cm diameter and 20 to 30 cm length), tagged, weighed, and measured for size and decay class (further details in Liu *et al.*^8^). The three decay classes were DKC1 = a knife could not penetrate, DKC2 = a knife could slightly penetrate with appreciable resistance, DKC3 = a knife could deeply penetrate with little resistance^51^. We used similar-sized pieces to control for potential effects of wood size on fungal communities^52^. The pieces were placed on the forest floor within a 60 × 3 m belt transect following an elevation contour. We collected source wood from within 500 m of this transect, utilizing about 5% of downed woody debris from these species in these decay classes, potentially arising from about 6000 source trees (D.A. Schaefer, unpublished data).

Each wood piece was initially weighed with a GLL portable electronic balance (accuracy 0.5 g) and for moisture content with an Extech MO210 moisture meter (calibrated as in Liu *et al.*^8^). Their volumes were calculated as cylinders, based on length and the average of 5 circumferential measurements along their lengths. From those, initial weight, volume, and density were all calculated. Oven drying of these wood pieces was not done, because it would have altered microbial communities and wood chemistry.

*CO_2_ emissions-rate measurements*. — Individual wood-piece CO_2_ release rates were measured in the field in a closed, ventilated chamber (10 L) connected to an infrared gas analyzer (Licor 820, Lincoln, NE, USA). After chamber closure and initial stabilization, linear CO_2_ concentration increase rates were logged for at least 5 min. Pieces remained in the field for CO_2_ measurements (within 5 m) and were handled carefully to limit fragmentation. Temperature and moisture were measured for each sample at each sampling time. The wood-piece CO_2_ release rate (R_WD_, μmol C g^-1^ h^-1^) was calculated as follows:

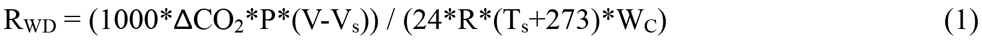

where ΔCO_2_ represents the measured CO_2_ concentration increase (ppm day^-1^), P is the internal pressure (kPa; measured by the Licor 820), V is the volume of the system (10.08 L, including the chamber volume and tubing volume and Licor optical path), V_s_ is the volume of the wood piece (L), R is the gas constant (8.314 L^-1^ kPa^-1^ K^-1^ mol^-1^), T_s_ is the wood temperature (°C), and WC is the carbon weight of each piece (g; 47% of its dry weight).

These measurements were made eight times from September 2010 to June 2013, approximately every four months (Fig. 1).

*Sampling for genetic analysis*. — The larger ongoing study includes three species, each having three decay classes, thus producing nine strata. 65 wood pieces were selected from all nine strata. In addition, for each stratum, we calculated a mean CO_2_ emission rate during 2010-2012, and the pieces chosen for this experiment included ones that were consistently below, at, or above the mean rate, and also some that showed variation around the mean over time. Pieces for each of these subgroups were selected at random from the larger study. This stratified random sampling ensured that all nine groups and the full range of CO_2_ emission variation were represented in the metabarcoded samples. Those wood pieces were collected in September 2012, and 59 of those samples were recollected in June 2013 (6 samples were not relocated in June 2013, having lost their tags or been buried under new litter). We chose June and September because CO_2_ emissions from wood are typically higher in June and lower in September, despite similarly warm temperatures and high moisture availability^8^ (Fig. 1 top). Superficial litter and bark were removed from each wood piece with a flame-sterilized knife before drilling. An electric drill with a flame-sterilized, 11-mm drill bit was used to extract wood powder at three holes located near the ends and the middle of each wood piece. Each sample consisted of ~5 cm^3^ pooled material from those three holes, collected onto aluminum-foil sheets, and then immediately stored in 50 ml tubes and frozen at −20 °C for 2-3 days until transport on ice packs to our laboratory 10 hr away, where they were stored at −40 °C until extractions.

*Comparing cumulative CO_2_ emissions and gravimetric weight losses*. — Sixty-four additional wood pieces (*i.e*. not used for metabarcoding) were retrieved from the field in April 2013 to test the extent to which CO_2_-emissions-estimated mass loss (averaged over the measurements taken from the year 2010 experiment start) accurately estimated directly measured mass loss (gravimetric weight loss). In the laboratory, these wood pieces were re-measured for volume (as above), and twenty-two wood pieces exhibiting >15% volumetric weight loss since the start of the experiment, indicating substantial fragmentation in the field, were excluded from the analysis. The remaining wood pieces were dried at 70 °C to constant weights and then individually weighed to the nearest 0.1 g on an electronic balance.

CO_2_-estimated mass loss was positively correlated with gravimetric loss (linear regression, Gravimetric loss in grams over 3 years = 30.7 + 0.741 * CO_2_-based-decomposition Cd (R^2^ = 0. 769, n = 17, p <0.01); 28.7 + 0.56 * Cd (R^2^ = 0.762, n = 13, p <0.01); and 40.3 + 0.468 * Cd (R^2^ = 0.592, n = 13, p <0.01), for DKC1, 2, and 3, respectively. Inspection of scatterplots (Supporting Information S5) revealed that the main discrepancy was that CO_2_-emission mass-loss estimates slightly underestimated small mass losses.

*DNA extraction, PCR, and 454pyrosequencing of ITS2 amplicons*. — Total DNA was extracted from each sample of wood powder by adding 10 mL CTAB buffer (2% cetyl trimethyl ammonium bromide, 50 mM NaCl, 5 mM EDTA, 10 mM Tris, pH 8) and 20 μL β-mercaptoethanol per 5 cm^3^ of sample, homogenizing with a TH-02 homogenizer (Omni International, Kennesaw, GA USA) for 5 min at room temperature, incubating at 65 °C for 1 hour, centrifuging at 4,000 rpm for 1 min. After centrifugation, the supernatant was transferred to new Axygen^®^ 2.0 mL microcentrifuge tubes and extracted using one volume of chloroform by vortexing for 20 min and centrifuged at 4 °C, 12,000 rpm for 10 min. The supernatant was then transferred to new microcentrifuge tubes and precipitated with 1.5 volumes of precooled isopropanol at −20 °C overnight. After centrifugation at 4 °C, 12,000 rpm for 20 min, the precipitate was washed with 70% ethanol and dissolved in 100 μL TE buffer (10 mM Tris, 1 mM EDTA, pH 8.0). DNA was purified by using QIAquick PCR Purification Kit. The quantity and quality of purified DNA was assessed with a Nanodrop 2000 spectrophotometer (Thermo Fisher Scientific, Wilmington, DE, USA). Samples were PCR amplified using the forward primer ITS3_KYO2 5’-GATGAAGAACGYAGYRAA-3’ ^18^ and reverse primer ITS4 5’-TCCTCCGCTTATTGATATGC-3’ ^53^. The standard Roche A-adaptor and a unique 10 bp MID (Multiplex Identifier) tag for each sample were attached to the forward primer. PCRs were performed using approximately 10 ng DNA in a 20 μL reaction mixture containing 2 μL of 10X buffer (Mg^2+^ Plus), 0.02 mM dNTPs, 20 μg Bovine Serum Albumin, 1 μL DMSO, 0.4 μM of each primer, and 0.5 U HotStart Taq DNA polymerase (TaKaRa Biotechnology Co., Dalian, China) under a temperature profile of 95 °C for 10 min, followed by 35 cycles of 94 °C for 20 sec, 47 °C for 30 sec, and 72 °C for 2 min, and final extension at 72 °C for 7 min. For pyrosequencing, PCR products were gel-purified using Qiagen QIAquick PCR purification kit, quantified using the Quant-iT PicoGreen dsDNA Assay kit (Invitrogen, Grand Island, NY, USA), pooled and A-amplicon-sequenced on a Roche GS FLX (Branford, Connecticut, USA) at the Kunming Institute of Zoology.

*Bioinformatic analyses*. — The sequences obtained were run through a pipeline for quality control, denoising and chimera removal, OTU-picking and taxonomic assignment. Quality Control: Header sequences and low-quality reads were removed from the raw output in the QIIME 1.8.0 environment (*split libraries.py:* −l 100 −L 500 −H 30)^54^. The 65 samples collected in 2012 had been sequenced on four 1/8 regions, producing 679,361 raw reads and 525,679 post-quality-control reads (mean read length 286 bp). The 59 samples collected in 2013 had been sequenced on three 1/8 regions, producing 327,226 raw reads and 256,996 post-quality-control reads (mean read length 308 bp). These amplicon lengths were consistent with the expected mean length for ITS2 (327.2 bp, SD=40) reported by Toju *et al.*^18^.

*Denoising and chimera removal: Denoiser* in QIIME^55^ was used to remove characteristic 454 sequencing errors. Next, ITSx 1.0.3^56^ was used to extract the variable ITS2 region from the whole reads (i.e. conserved 5.8S and LSU flanking sequences were stripped) and to remove non-fungal-ITS reads. The extracted sequences were clustered at 99% similarity with USEARCH v7.0.1090^57^ to remove replicate sequences and chimeras. *OTU-picking and taxonomic assignment*: We used two methods to cluster the reads into OTUs. First, we used a reference-based method in QIIME (*pick_open_reference_otus.py*: max_accepts 20 max_rejects 500 stepwords 20 length 12-suppress_align_and_tree) in which reads were first clustered by matching at 97% similarity to the UNITE 12_11 fungal database^58^, which itself had previously been clustered at 97% similarity for use within QIIME. Unassigned reads (the vast majority) were then clustered *de novo* using the *uclust* option at 97% similarity, producing 1,807 OTUs in total. For these latter OTUs, we attempted to assign taxonomies using QIIME’s *assigntaxonomy.py* against the UNITE database. Second, we performed *de novo* 97%-similarity clustering with CROP 1.33^59^, producing 1,565 OTUs. We assigned taxonomies against Genbank using the NNCauto and QCauto methods in Claident^60^.

Sequence data are deposited at datadryad.org (doi: to be assigned) and in GENBANK’s Short Read Archive (Accession number: PRJNA252416). An example bioinformatic script is in Supplementary Information and also deposited at datadryad.org (doi: to be assigned).

*Statistical analyses*. — Analyses were performed using *vegan* 2.0-10^61^ and *mvabund* 3.8.4^62^ in *R* 3.1.0^63^. The HTML outputs of the R scripts are in Supplementary Information and also deposited at datadryad.org (doi: to be assigned). From both the OTU tables generated (CROP and *uclust*, see *Bioinformatic analysis*), we deleted one SN wood piece that had only 35 reads and then split the tables by wood species (LC, LX, and SN) and sample time (September 2012 and June 2013). We then generated a second pair of OTU tables by rarefying to the lowest read number per wood piece in the dataset (*rrarefy()* in *R*). Thus, for each wood species and sampling time, we have four OTU tables: CROP/non-rarefied, CROP/rarefied, *uclust*/non-rarefied, and *uclust*/rarefied. Finally, for each table, we used *vegan’s diversity()* function to estimate Shannon and Simpson diversity in each wood piece.

For each wood species, we used *lm()* in *R* to linearly regress emissions against fungal diversity. Thus, the June 2012 and September 2012 CO_2_ estimates were tested against the September 2012 fungal diversity estimate, and the June 2013 CO_2_ estimate was tested against the June 2013 fungal diversity estimate. Residuals were all adjudged visually to be near normally distributed, but with small indications of nonlinearity. We ignored the nonlinearities because the residuals suggested accelerating CO_2_ emissions at the lowest fungal diversity, making our results conservative. In trial models, we also tested for significant effects of wood surface temperature and decay class (see *Experimental setup*), but they did not interact significantly with the fungal diversity term and mostly did not enter significantly as additive terms, and so have been omitted here for simplicity. Liu *et al.*^8^ also did not find a correlation between decay class and CO_2_ emissions rates for these wood pieces.

To test whether a ‘pure diversity’ effect is sufficient to explain the observed diversity-function relationships, we use a method by Sandau *et al.*^24^ where community similarities (1-Jaccard binary) between all wood pieces are used to create a variance-covariance matrix that is then included in the linear regressions, thus taking into account potential non-independence of wood pieces due to the fact that some communities are similar to each other (Supporting Information S3). In Supporting Information S4, we also use conventional community analyses to test for an effect of community composition on CO_2_ emissions. We limit our tests to June 2012 and 2013, as only these exhibited significant declines in emissions with fungal-species diversity (Fig. 1A, C, D, F).

## Acknowledgments

We thank our two reviewers for their very helpful suggestions. CYY, DWY, CXY, XYW and CYW were supported by Yunnan Province (20080A001), the Chinese Academy of Sciences (0902281081, KSCX2-YW-Z-1027), the National Natural Science Foundation of China (31400470, 31170498), the Ministry of Science and Technology of China (2012FY110800), the University of East Anglia, and the State Key Laboratory of Genetic Resources and Evolution at the Kunming Institute of Zoology. DAS and WJL were supported by the Asia-Pacific Network for Global Change Research (ARCP 2009-18MY), the National Natural Science Foundation of China (30970535 and 41271278), the Chinese Academy of Sciences 135 program (XTBG-T01), and the Xishuangbanna Tropical Botanical Garden. Zhang Yiping provided Ailao Mountain meteorological data; Yang Gunping identified the wood pieces; Qiao Lu and Liu Xianbin assisted in the field work.

## Author contributions

DAS, CYY, and DWY designed the study. CYY, WJL and DAS carried out the field work. CYY, CXY, XYW and CYW performed the DNA analyses. CYY and DWY carried out the bioinformatics. DWY and VP carried out the statistical analyses. DWY, DAS and CYY wrote the first draft. WJL and CXY commented on the manuscript.

## Additional information

DWY is co-founder of a UK company, NatureMetrics, which provides DNA metabarcoding services to the private and public sector. All other authors declare no conflicts of interest.

## Supplementary Information

### S1 Summary of Regression Analyses

**S1.** Summary of Regression analyses

| OTU-pickng | rarefaction | CO2 measurement | species | diversity | p-value | notes |
| --- | --- | --- | --- | --- | --- | --- |
| CROP | nonrarefied | Jun-12 | LC | Shannon | **0.007** |  |
| CROP | nonrarefied | Jun-12 | LC | Simpson | 0.015 |  |
| CROP | nonrarefied | Jun-12 | LX | Shannon | **0.080** |  |
| CROP | nonrarefied | Jun-12 | LX | Simpson | 0.040 |  |
| CROP | nonrarefied | Jun-12 | SN | Shannon | **0.027** | removed influential woodpiece=110 |
| CROP | nonrarefied | Jun-12 | SN | Simpson | 0.018 | removed influential woodpiece=110 |
| CROP | nonrarefied | Sep-12 | LC | Shannon | **0.460** |  |
| CROP | nonrarefied | Sep-12 | LC | Simpson | 0.642 |  |
| CROP | nonrarefied | Sep-12 | LX | Shannon | **0.116** |  |
| CROP | nonrarefied | Sep-12 | LX | Simpson | 0.057 |  |
| CROP | nonrarefied | Sep-12 | SN | Shannon | **0.700** |  |
| CROP | nonrarefied | Sep-12 | SN | Simpson | 0.474 |  |
| CROP | nonrarefied | Jun-13 | LC | Shannon | **0.003** |  |
| CROP | nonrarefied | Jun-13 | LC | Simpson | 0.000 |  |
| CROP | nonrarefied | Jun-13 | LX | Shannon | **0.025** |  |
| CROP | nonrarefied | Jun-13 | LX | Simpson | 0.016 |  |
| CROP | nonrarefied | Jun-13 | SN | Shannon | **0.003** |  |
| CROP | nonrarefied | Jun-13 | SN | Simpson | 0.001 |  |
| CROP | rarefied | Jun-12 | LC | Shannon | 0.005 |  |
| CROP | rarefied | Jun-12 | LC | Simpson | 0.014 |  |
| CROP | rarefied | Jun-12 | LX | Shannon | 0.084 |  |
| CROP | rarefied | Jun-12 | LX | Simpson | 0.044 |  |
| CROP | rarefied | Jun-12 | SN | Shannon | 0.020 | removed influential woodpiece=110 |
| CROP | rarefied | Jun-12 | SN | Simpson | 0.037 | removed influential woodpiece=110 |
| CROP | rarefied | Sep-12 | LC | Shannon | 0.563 |  |
| CROP | rarefied | Sep-12 | LC | Simpson | 0.572 |  |

The (bolded) Shannon results from this first section are plotted in Fig. 1A-C.

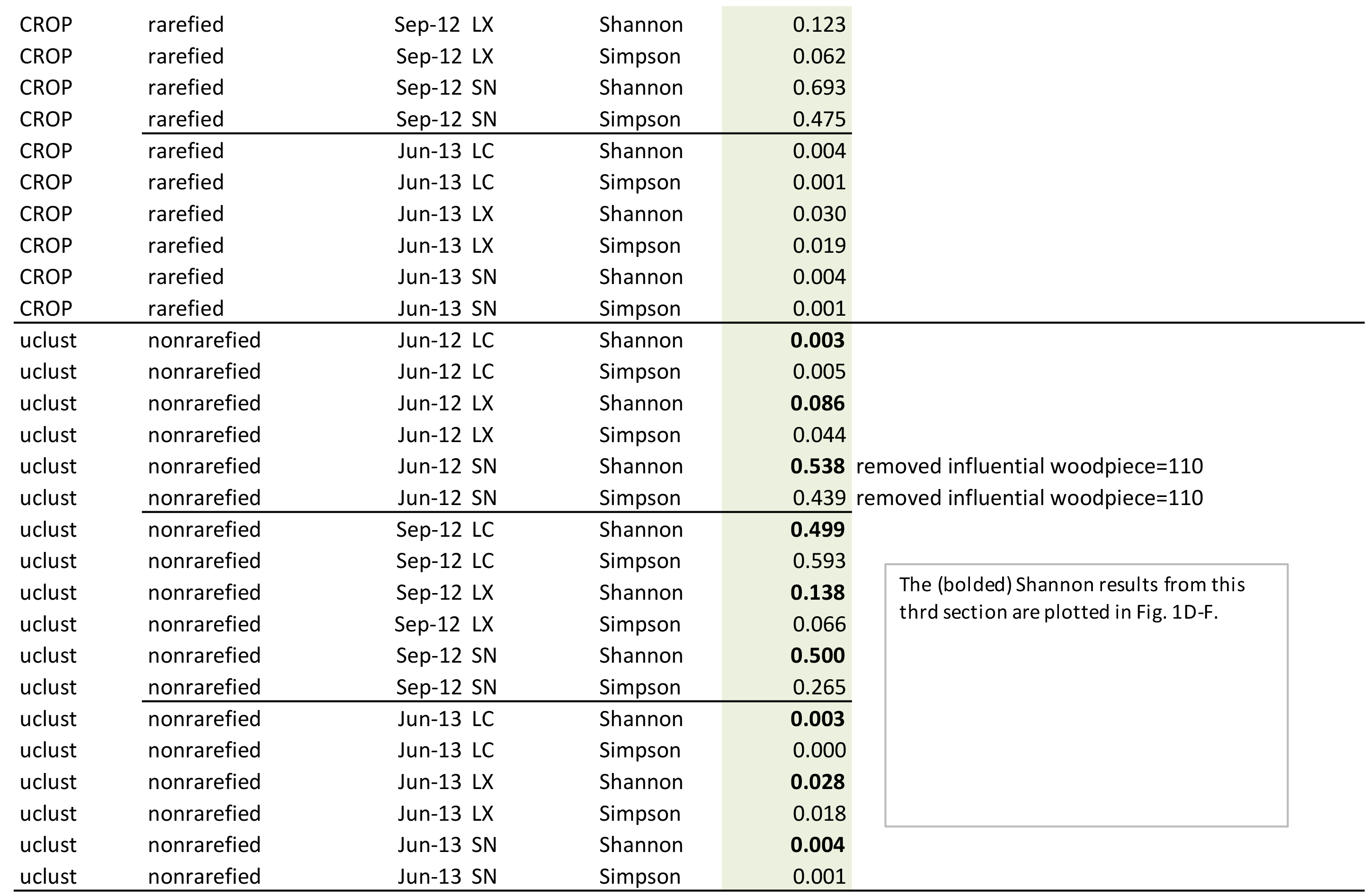

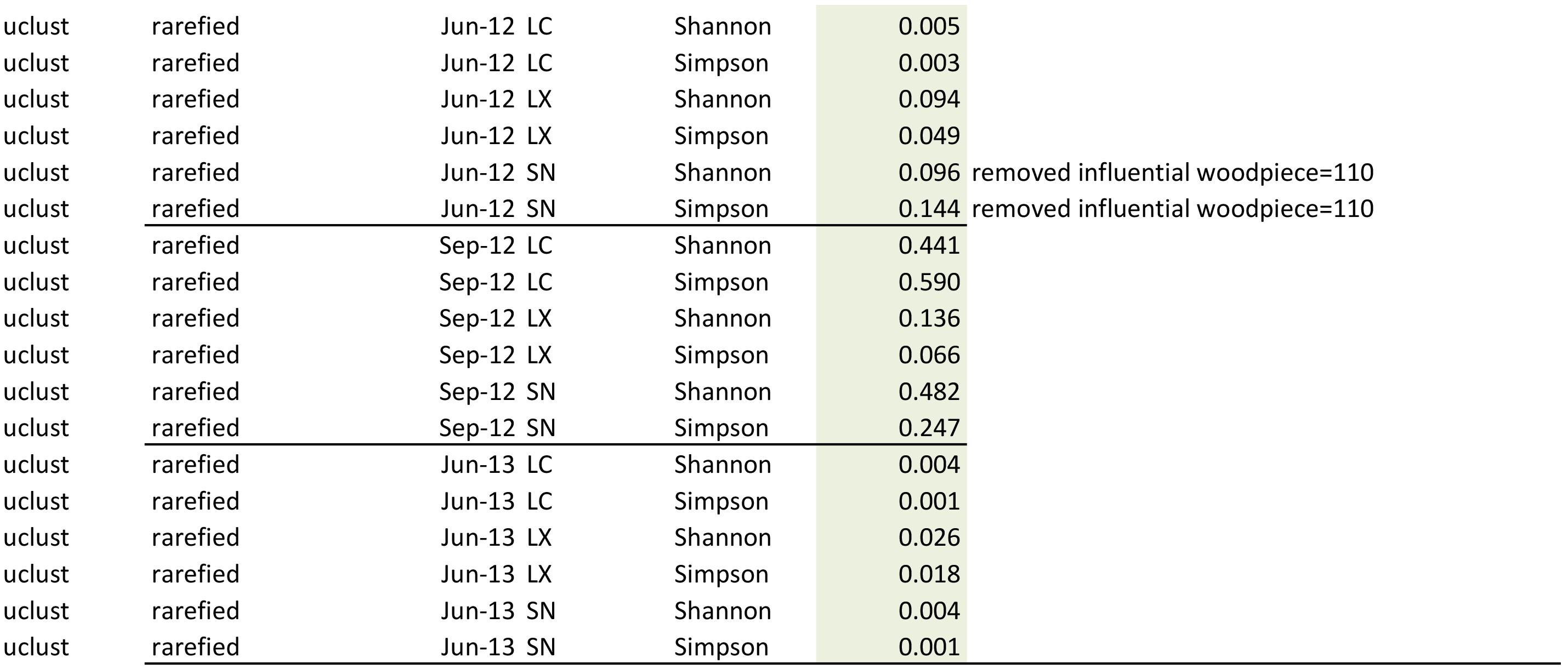

### S2 Wood chemistry analyses

#### Methods

In January 2013, 27 decaying wood samples from our larger study were removed from the field for chemical analyses. Included were 3 pieces from each combination of wood species (LC, LX, and SN) and decay class (DKC 1, 2, 3), spanning the range of previously observed decomposition rates. These pieces were dried at 60 °C to constant weights and milled to pass a 40 mesh (425 micron opening) sieve. They were analyzed for total carbon and nitrogen (Macro Elemental CN Analyzer, Vario MAX CN, Germany), for total phosphorus (continuous Flow Analyzer, Auto Analyzer 3, Germany) after acid digestion, and for lignin (Fibertec 2010 System, Fibertec, Denmark). The metabarcoded wood pieces remain in the field and were not analyzed.

#### Column headings explanation

Tag: Permanent identification number assigned to each piece of wood in this study Tree_species: LC = *Lithocarpus chintungensis*, LX = *Lithocarpus xylocarpus* (LX), and SN = *Schima noronhae*

Average_CO_2_: CO_2_ release rate for each wood piece from 2010 to 2012, average of 6 measurements, micromoles CO_2_ per dry gram of wood carbon per hour Total_carbon_g/kg: Carbon as grams per dry kilogram of wood weight Total_Nitrogen_g/kg: Nitrogen as grams per dry kilogram of wood weight Total_Phosphorous_g/kg: Phosphorus as grams per dry kilogram of wood weight lignin_%: Lignin as percent of dry wood-piece weight

| Tag | Tree_species | Decay_class | Average_CO<sub>2</sub> | Total_carbon_g/kg | Total_Nitrogen_g/kg | Total_Phosphorus_g/kg | lignin_% |
| --- | --- | --- | --- | --- | --- | --- | --- |
| 52 | LC | 1 | 0.529 | 458 | 3.96 | 0.20 | 26.8 |
| 160 | LC | 1 | 0.482 | 452 | 3.7 | 0.16 | 19.6 |
| 164 | LC | 1 | 0.335 | 455 | 3.51 | 0.12 | 19.7 |
| 2 | LC | 2 | 0.601 | 450 | 3.67 | 0.17 | 22.1 |
| 119 | LC | 2 | 1.980 | 496 | 4.30 | 0.22 | 18.6 |
| 214 | LC | 2 | 0.298 | 460 | 2.65 | 0.07 | 16.4 |
| 11 | LC | 3 | 0.450 | 458 | 3.08 | 0.13 | 23.7 |
| 67 | LC | 3 | 0.723 | 488 | 7.56 | 0.27 | 31.0 |
| 76 | LC | 3 | 0.404 | 491 | 4.69 | 0.11 | 25.2 |
| 92 | LX | 1 | 0.264 | 497 | 2.69 | 0.08 | 23.0 |
| 219 | LX | 1 | 0.358 | 454 | 3.33 | 0.20 | 19.3 |
| 223 | LX | 1 | 0.796 | 455 | 4.19 | 0.25 | 17.8 |
| 141 | LX | 2 | 1.146 | 492 | 5.78 | 0.36 | 16.0 |
| 142 | LX | 2 | 1.430 | 505 | 8.38 | 0.64 | 19.4 |
| 243 | LX | 2 | 0.392 | 454 | 3.44 | 0.11 | 24.8 |
| 191 | LX | 3 | 1.493 | 452 | 5.42 | 0.57 | 17.3 |
| 267 | LX | 3 | 1.454 | 459 | 7.40 | 0.37 | 24.0 |
| 269 | LX | 3 | 0.534 | 464 | 5.47 | 0.27 | 18.7 |
| 29 | SN | 1 | 0.269 | 455 | 2.29 | 0.06 | 16.7 |
| 40 | SN | 1 | 0.752 | 451 | 3.92 | 0.17 | 20.8 |
| 317 | SN | 1 | 0.463 | 527 | 2.42 | 0.12 | 20.6 |
| 7 | SN | 2 | 0.803 | 454 | 3.17 | 0.13 | 19.3 |
| 108 | SN | 2 | 0.980 | 453 | 1.45 | 0.07 | 16.2 |
| 313 | SN | 2 | 0.421 | 453 | 3.11 | 0.09 | 18.4 |
| 23 | SN | 3 | 0.449 | 470 | 3.19 | 0.14 | 19.2 |
| 25 | SN | 3 | 0.848 | 452 | 1.88 | 0.07 | 16.6 |
| 321 | SN | 3 | 0.335 | 503 | 3.13 | 0.10 | 21.8 |

Wood Chemistry: Supplementary Information S2

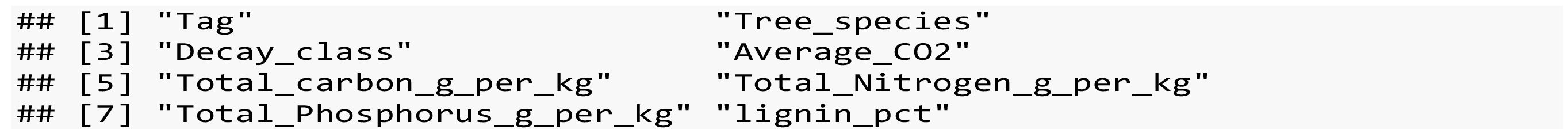

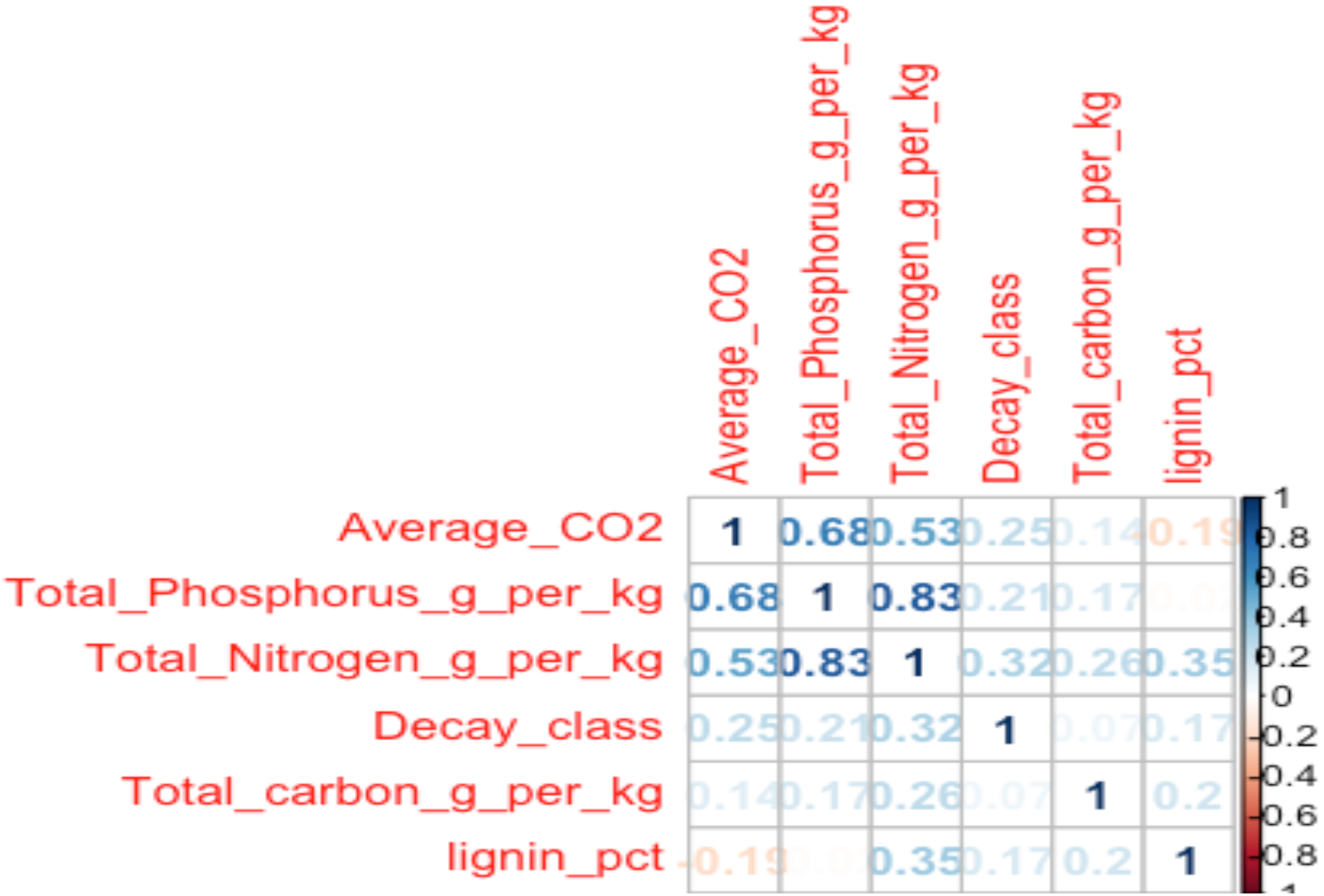

The high correlation between Total_Phosphorous_g_per_kg and Total_Nitrogen_g_per_kg means that we cannot include them in the same model (collinearity). So we run two separate models. We also run models separately for each decay class, because wood chemistry changes with age, due to selective retention of N and P while C is lost. This gives us 6 total models (3 decay classes X 2 models). As a result, we informally carry out a Table-wide correction of P-values by multiplying all P-values by 6 before assessing statistical significance.

Declay class 1

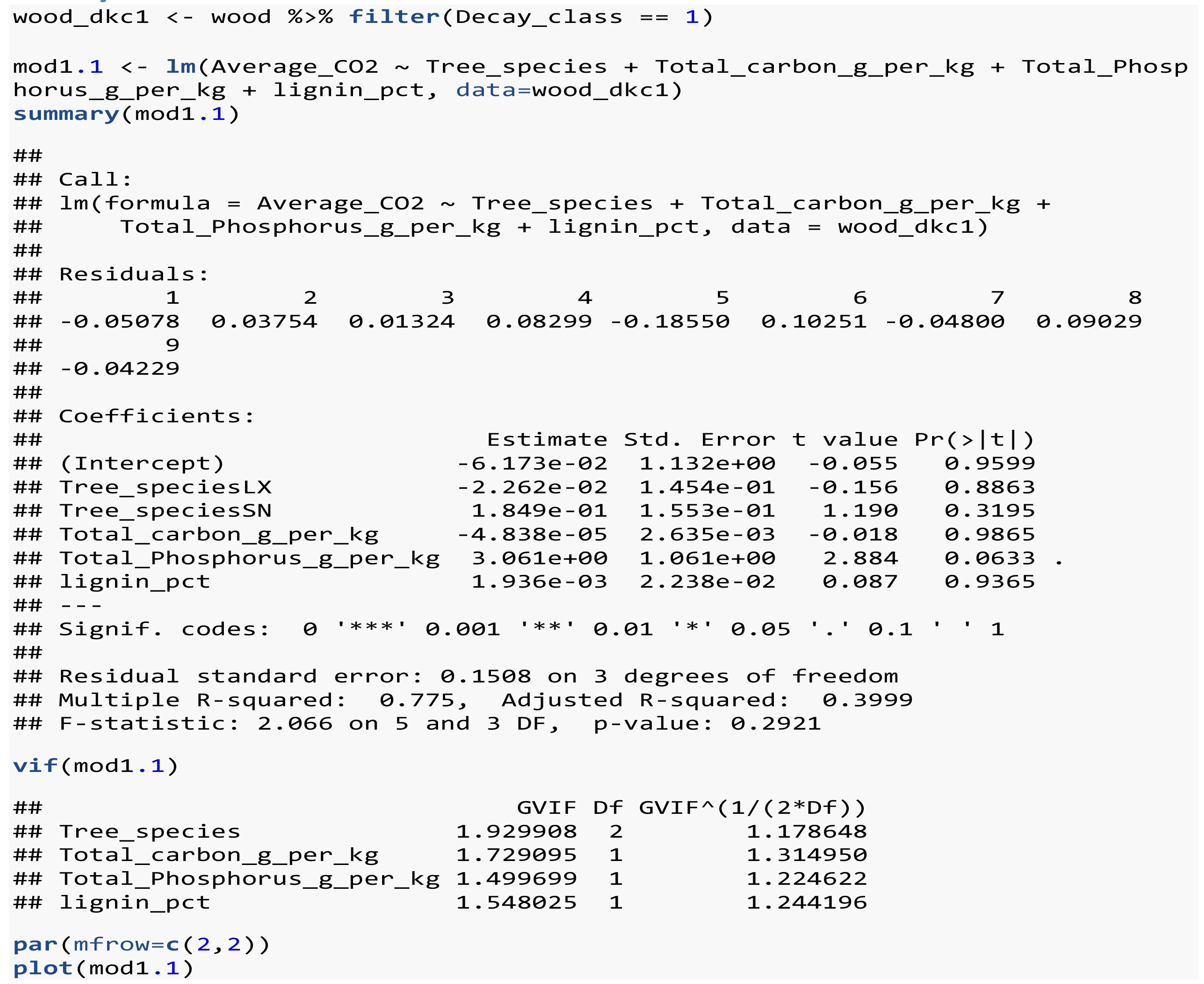

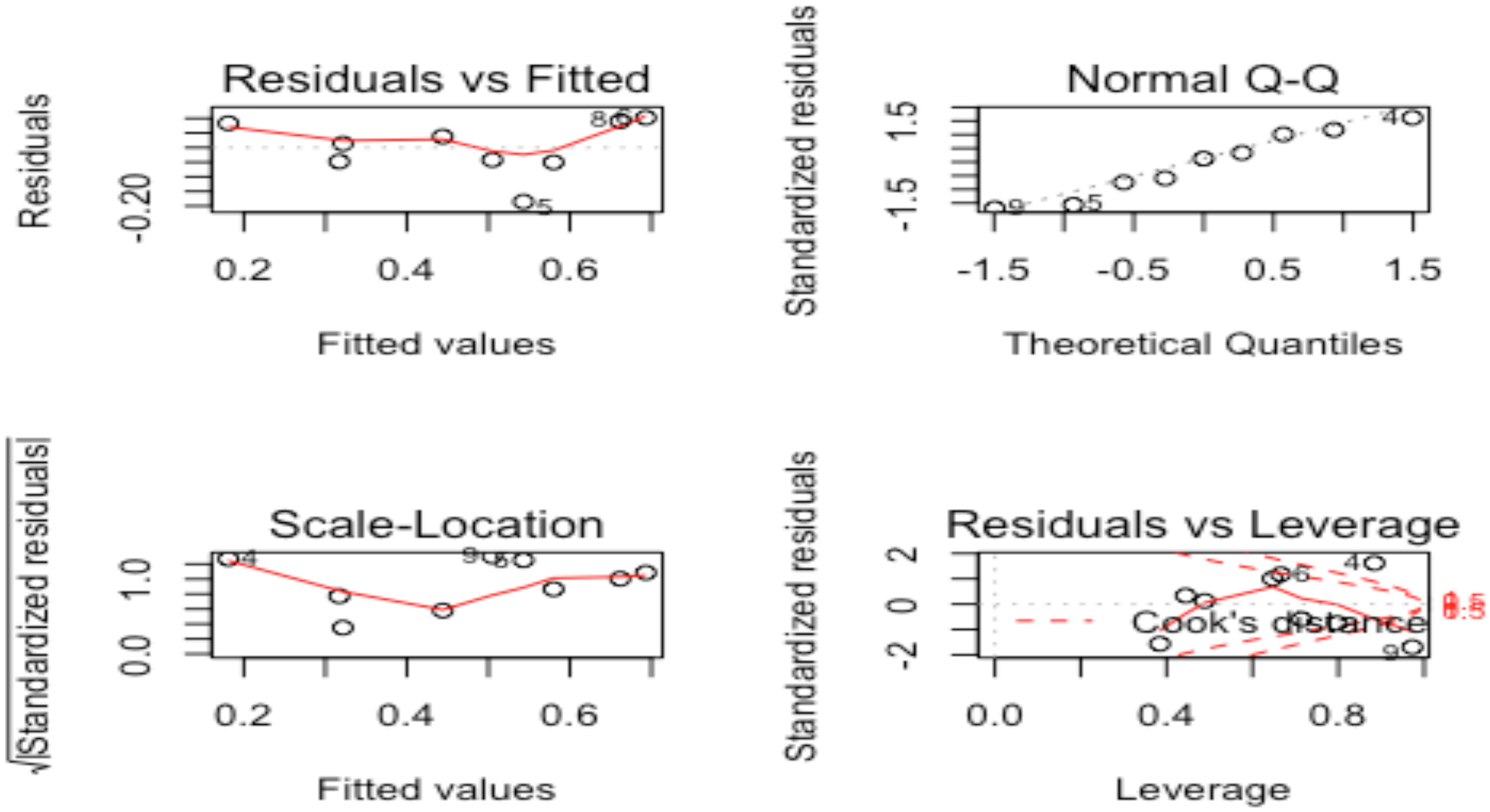

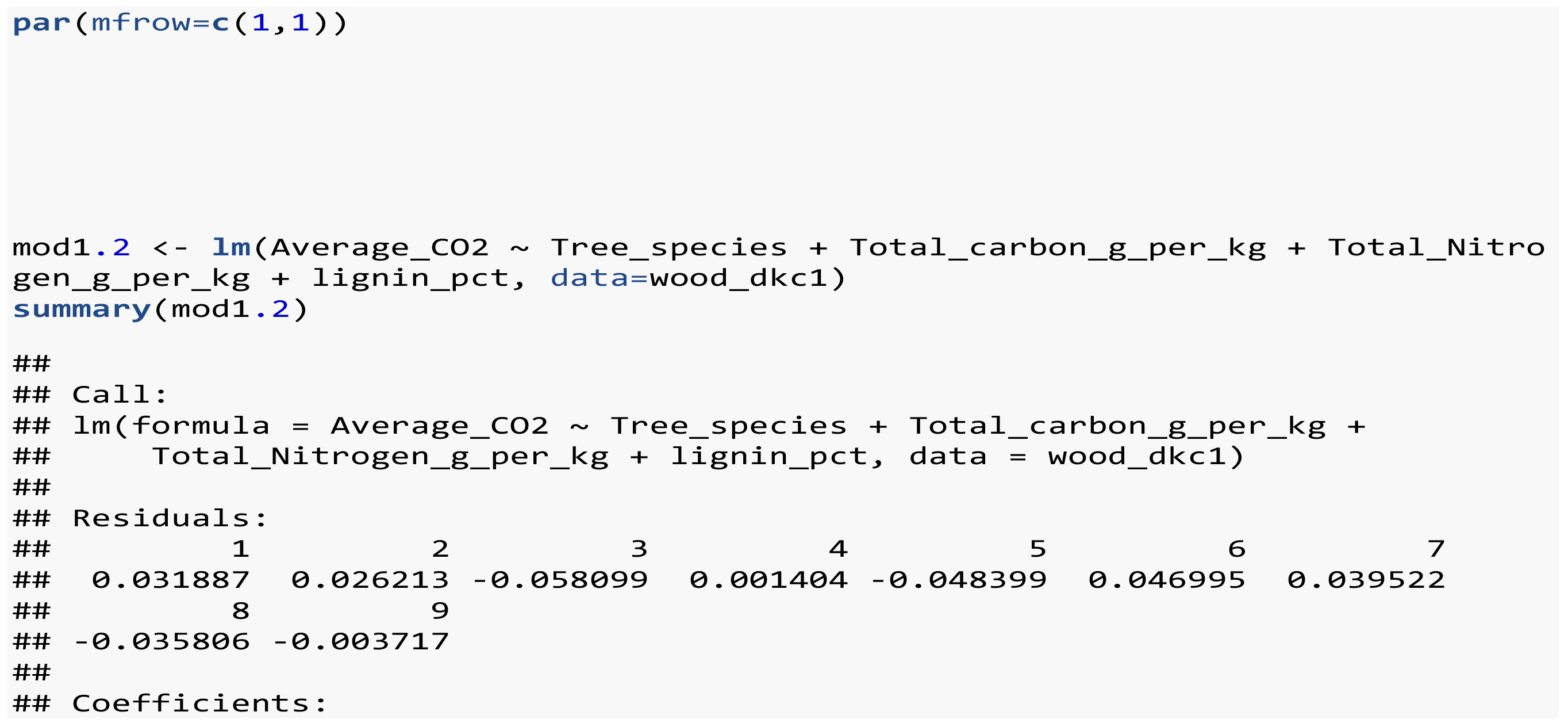

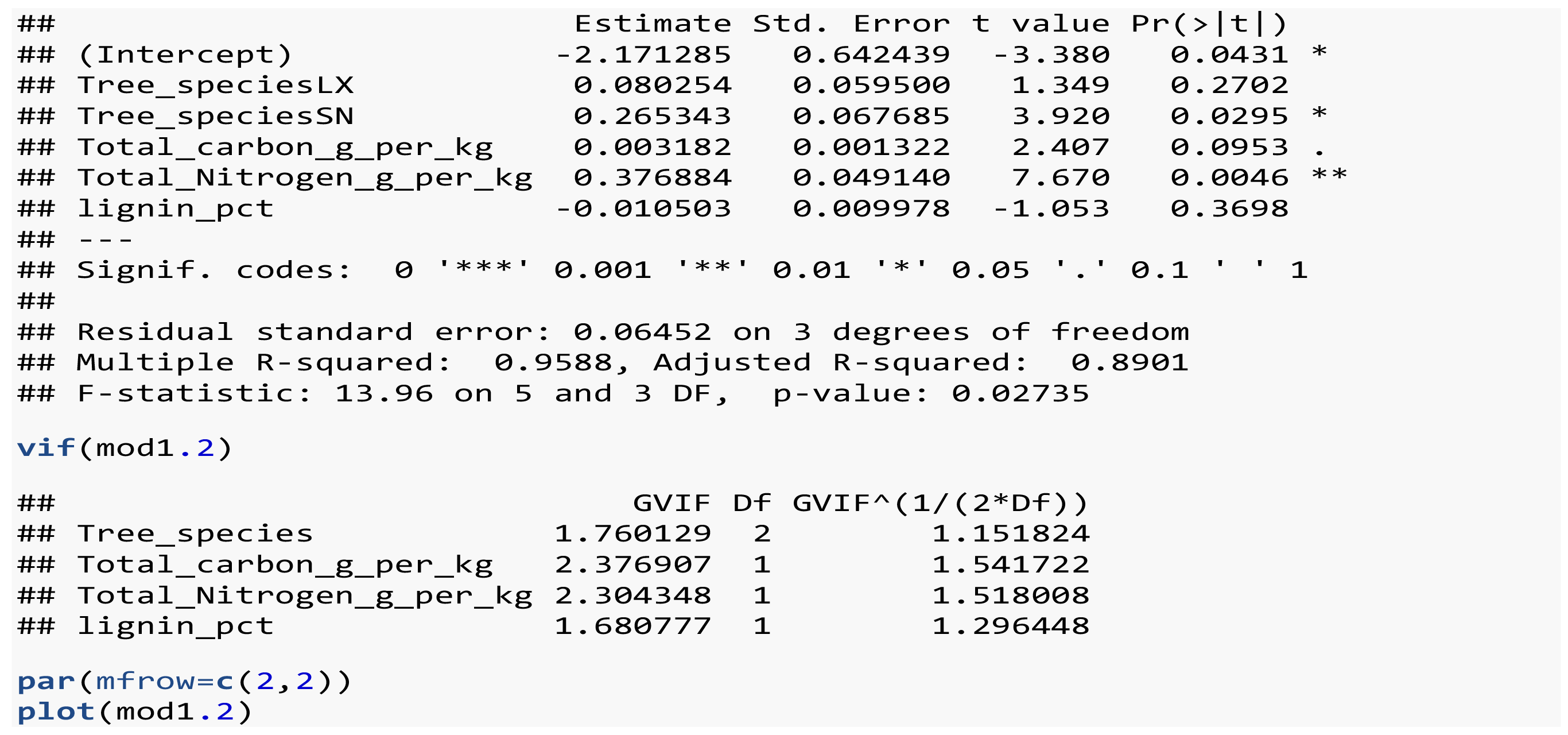

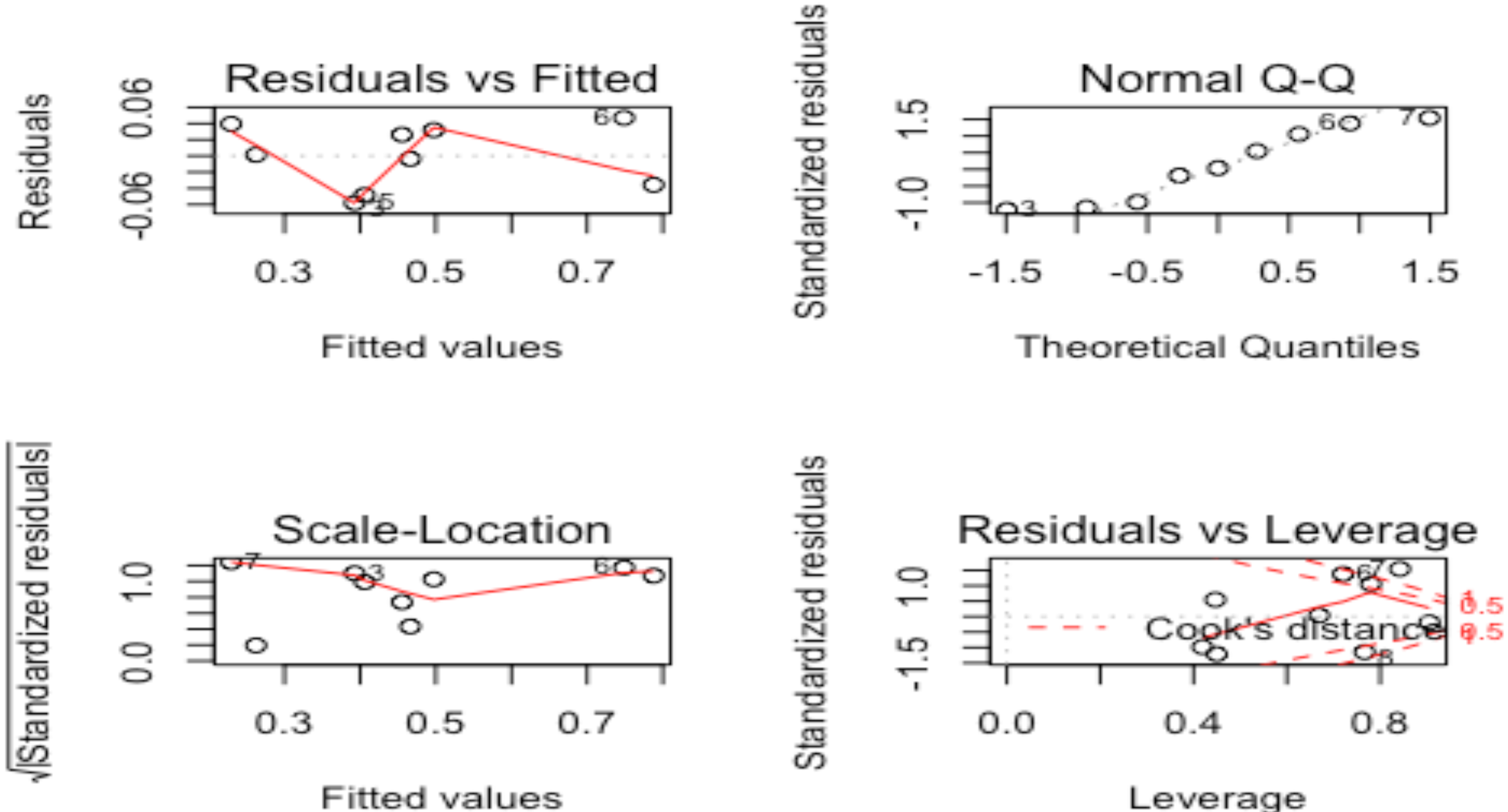

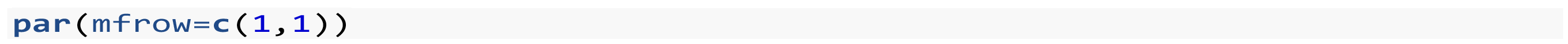

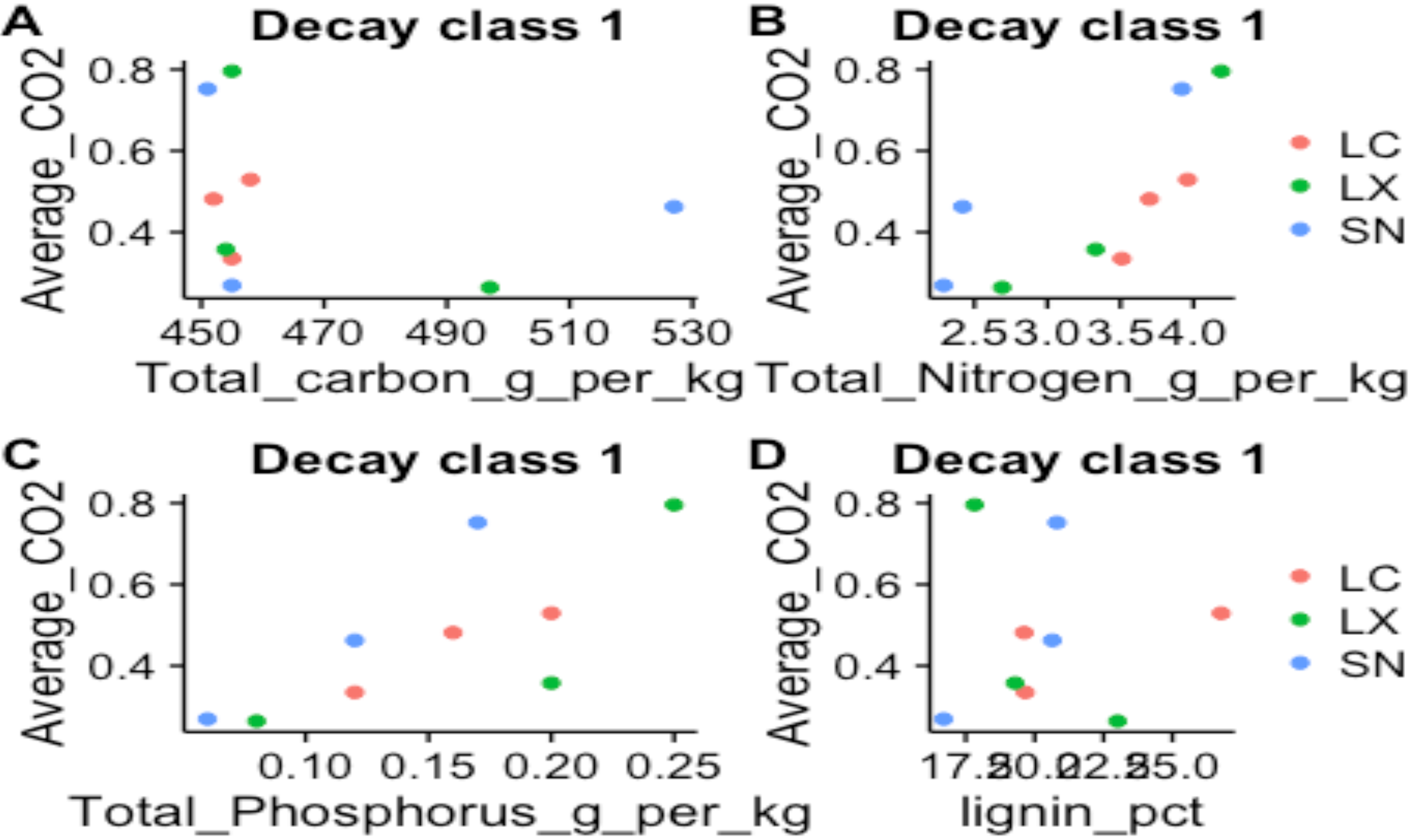

Declay class 2

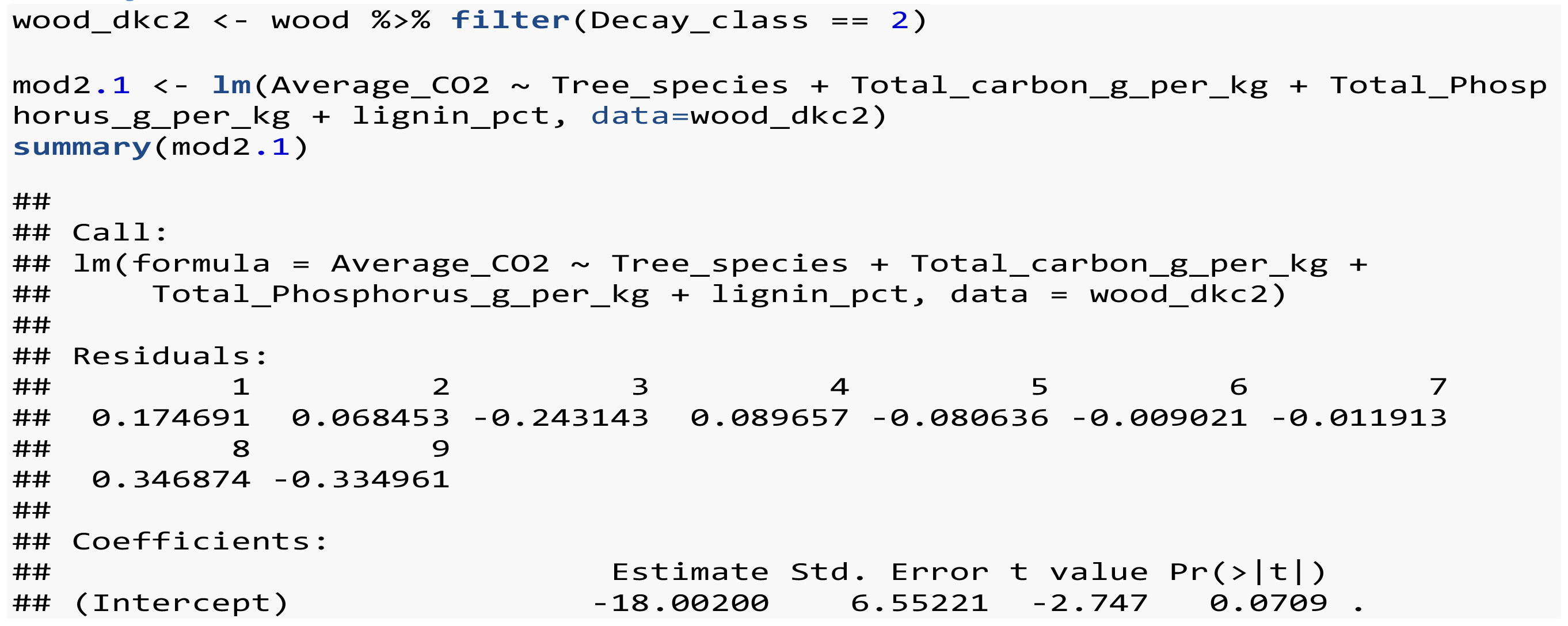

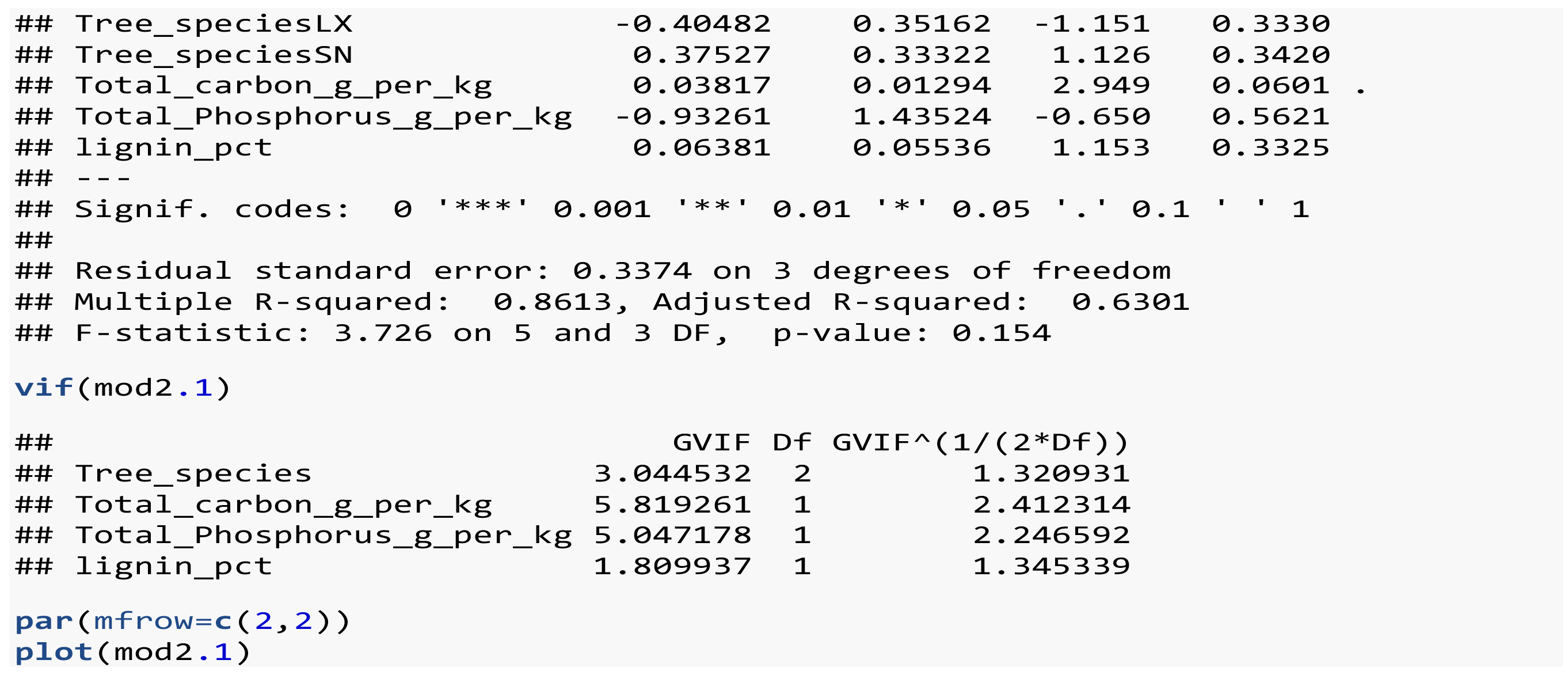

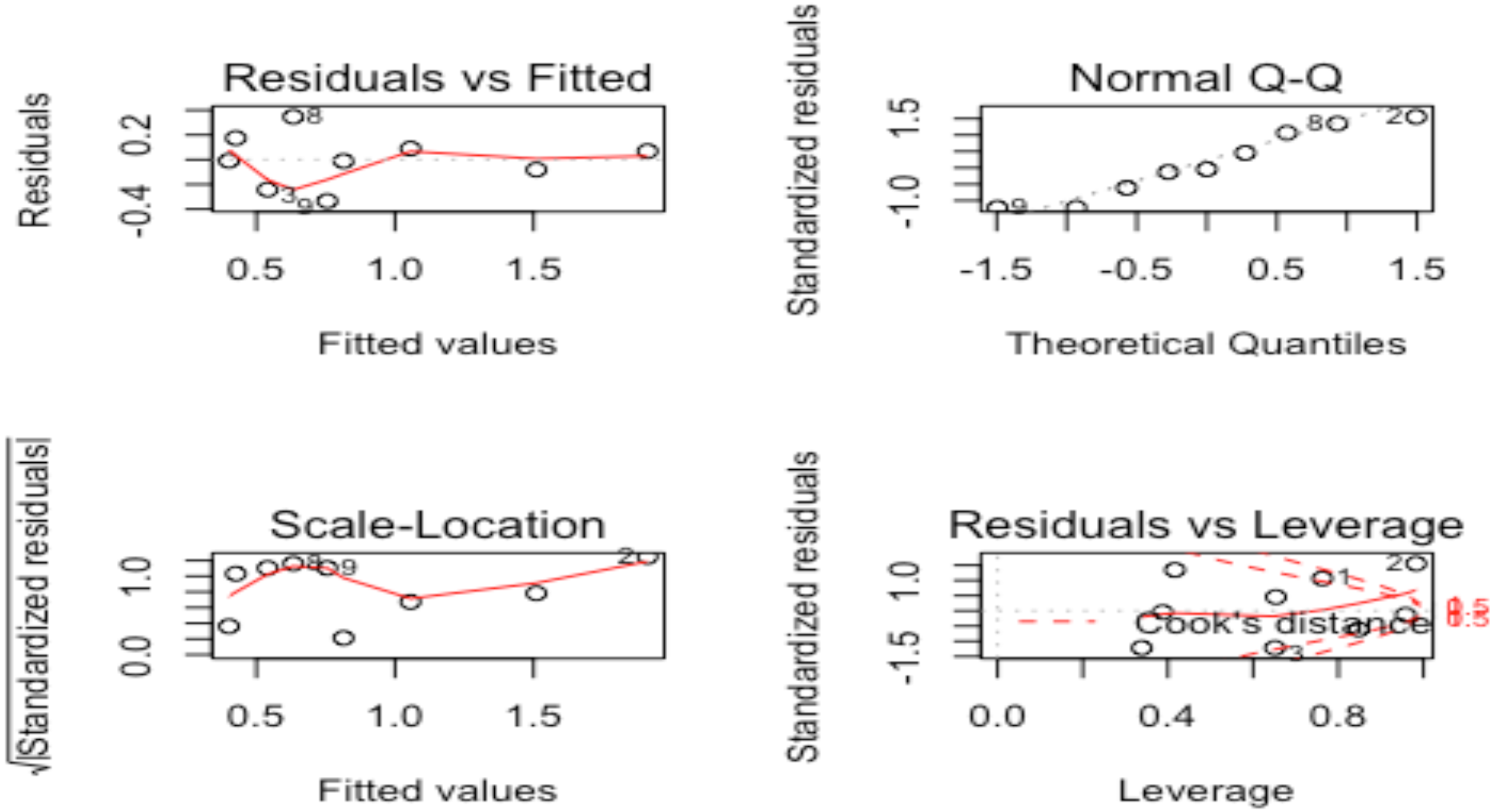

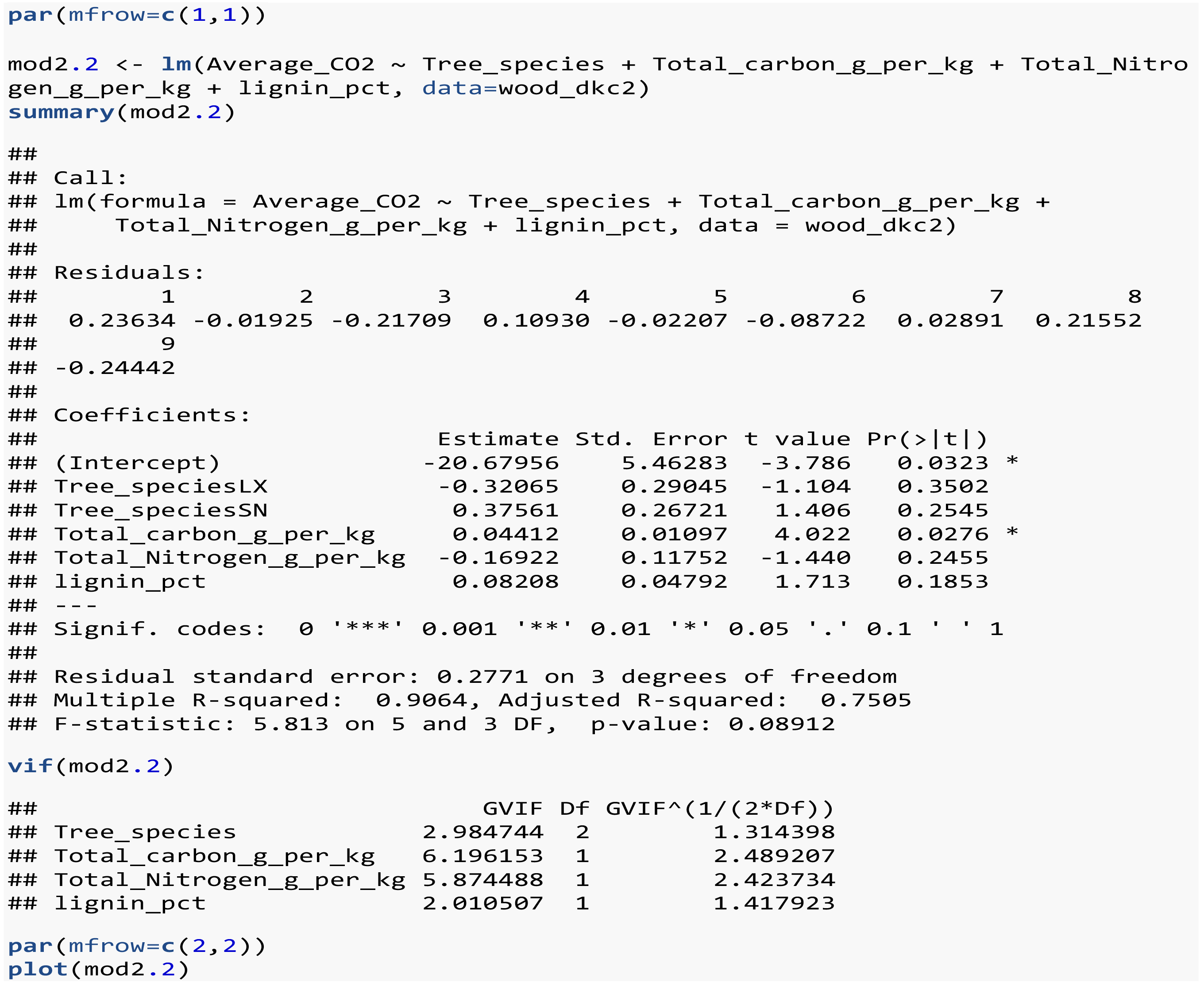

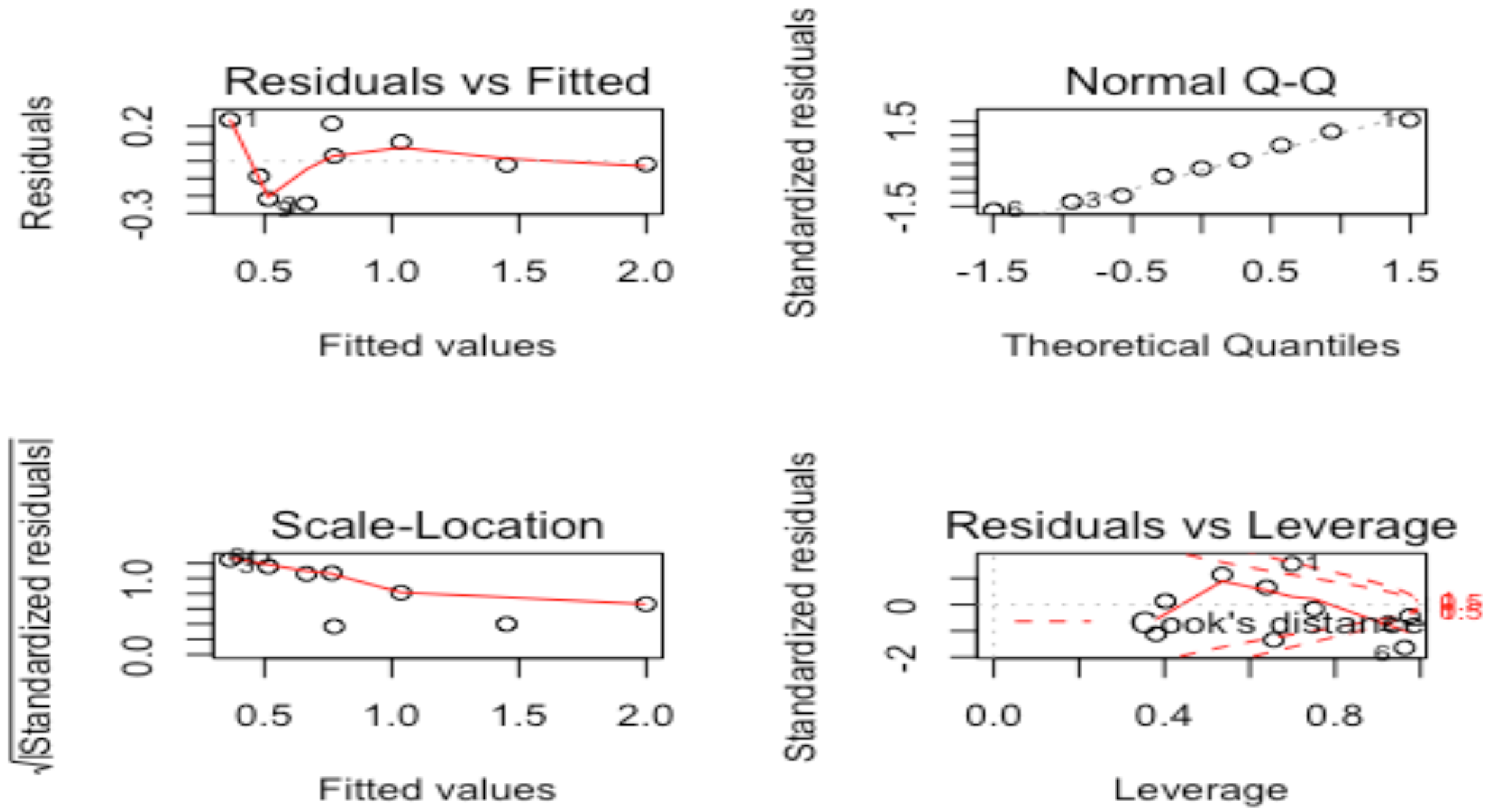

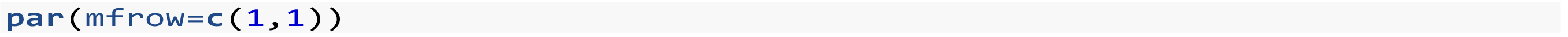

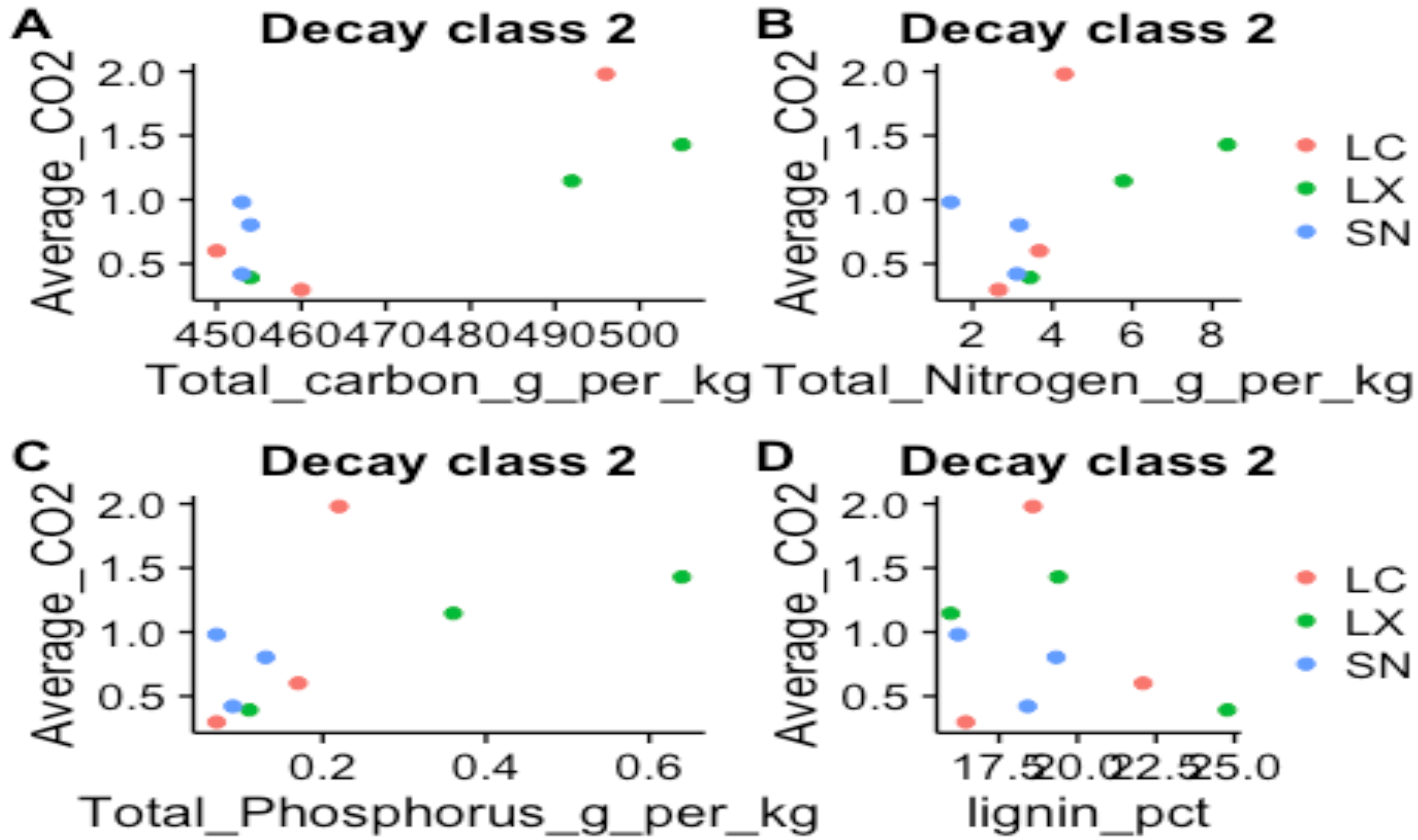

Declay class 3

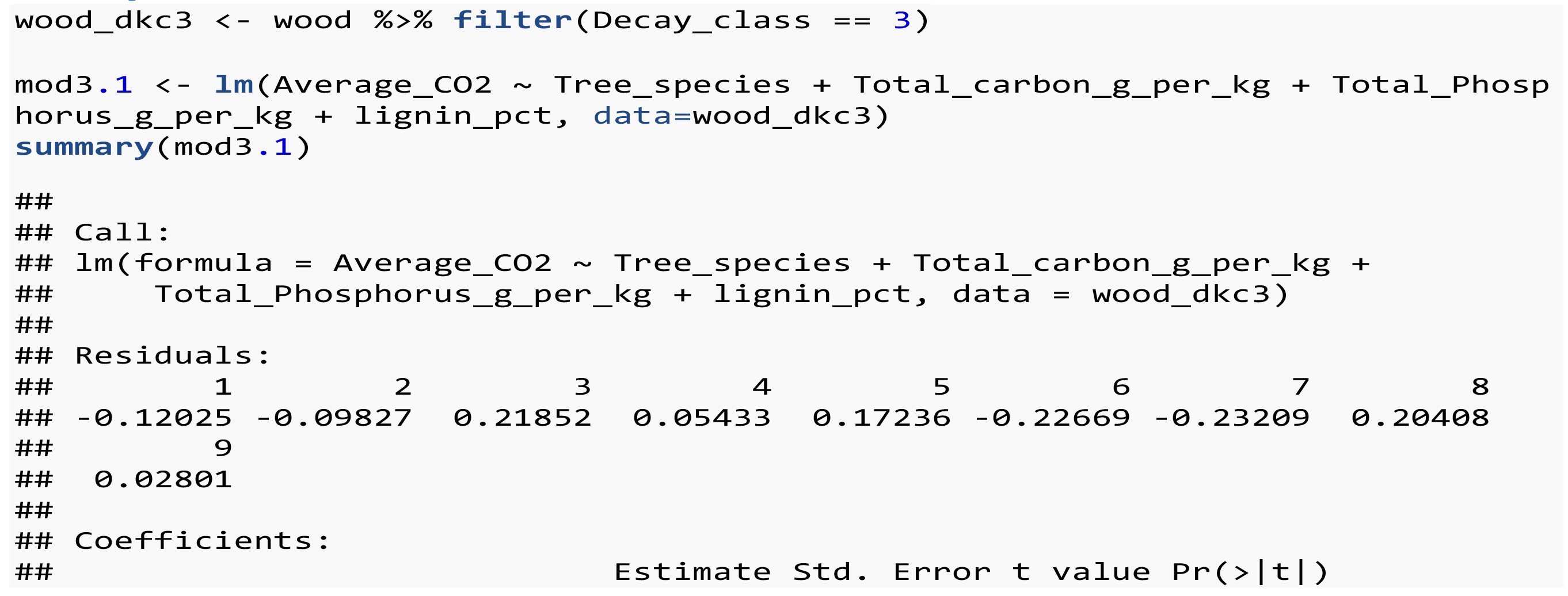

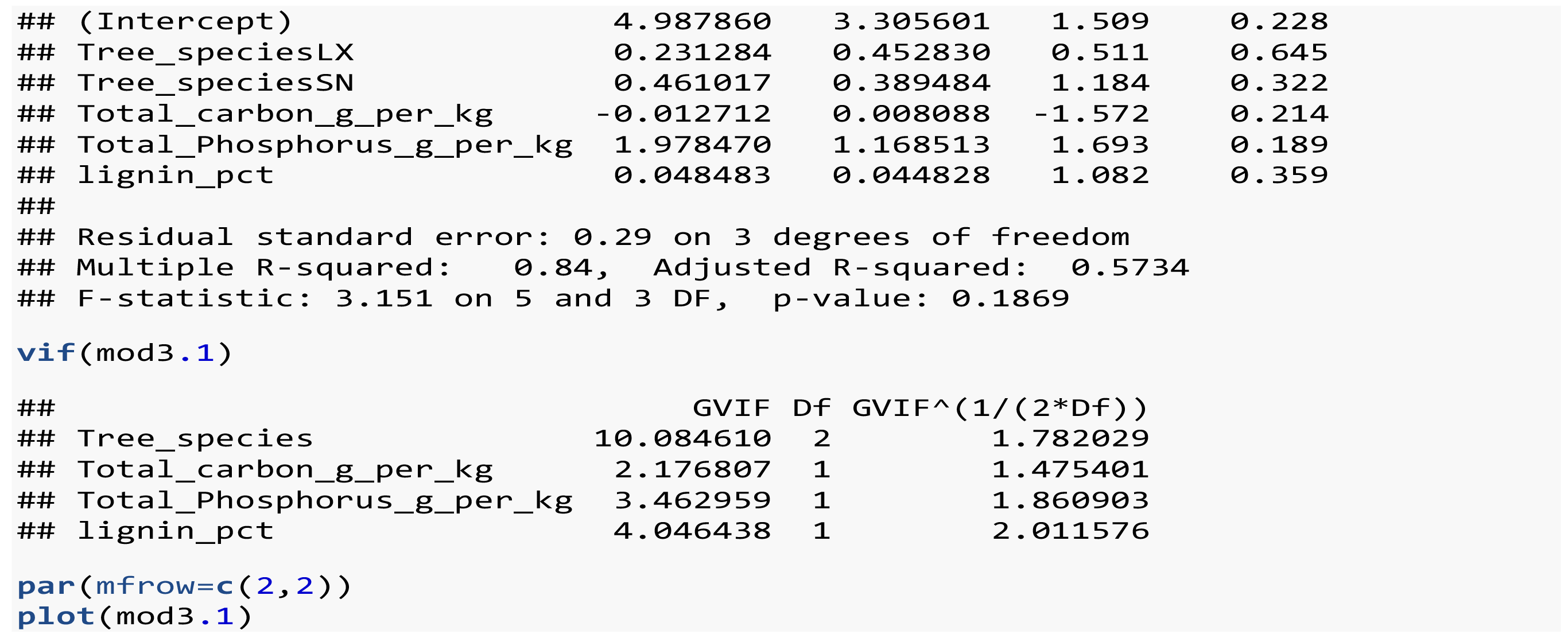

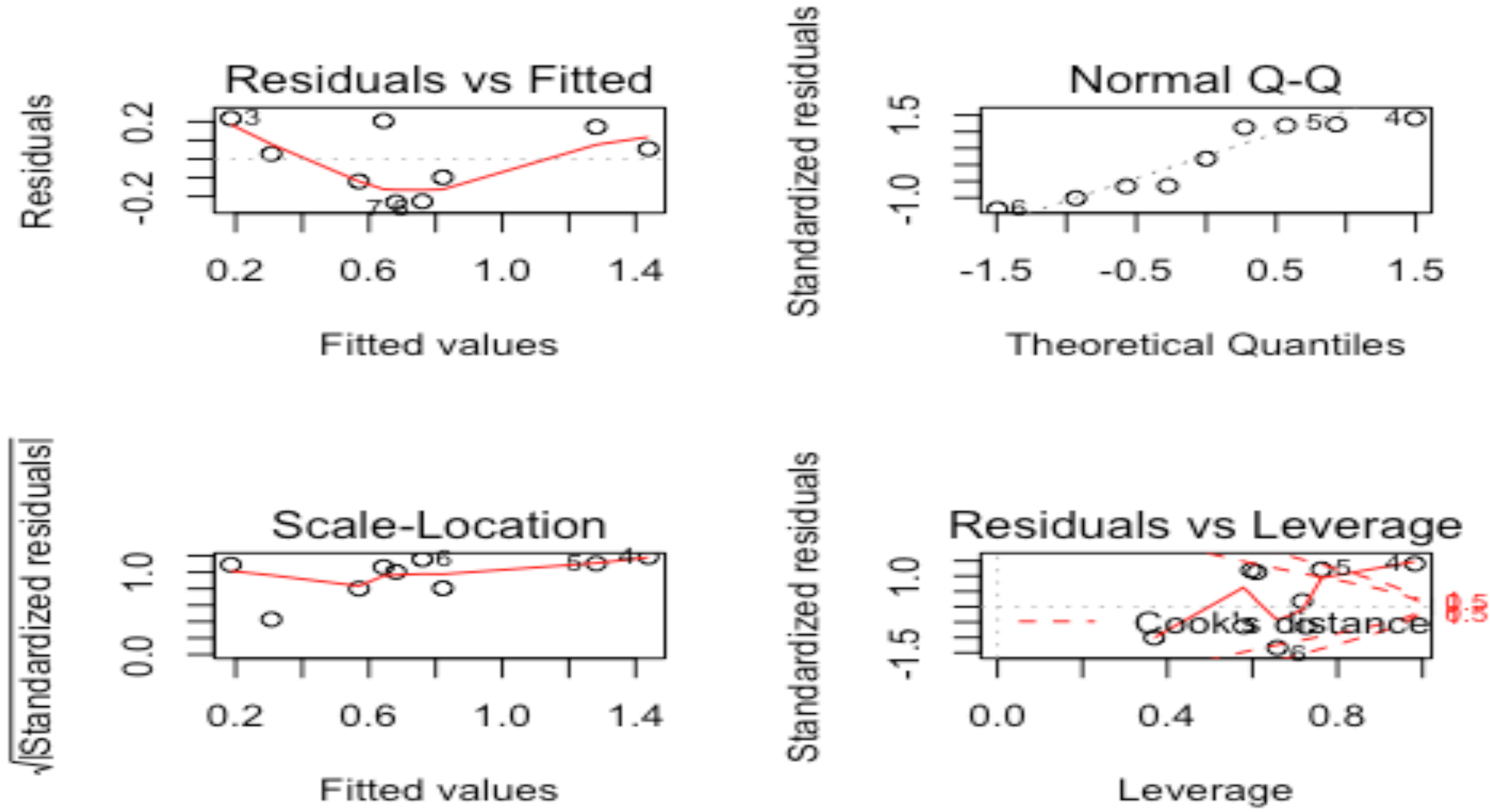

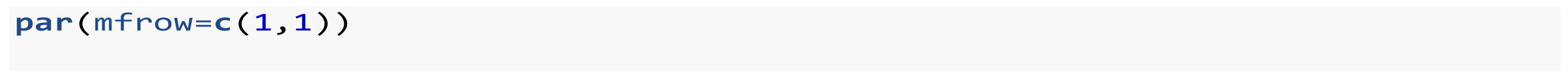

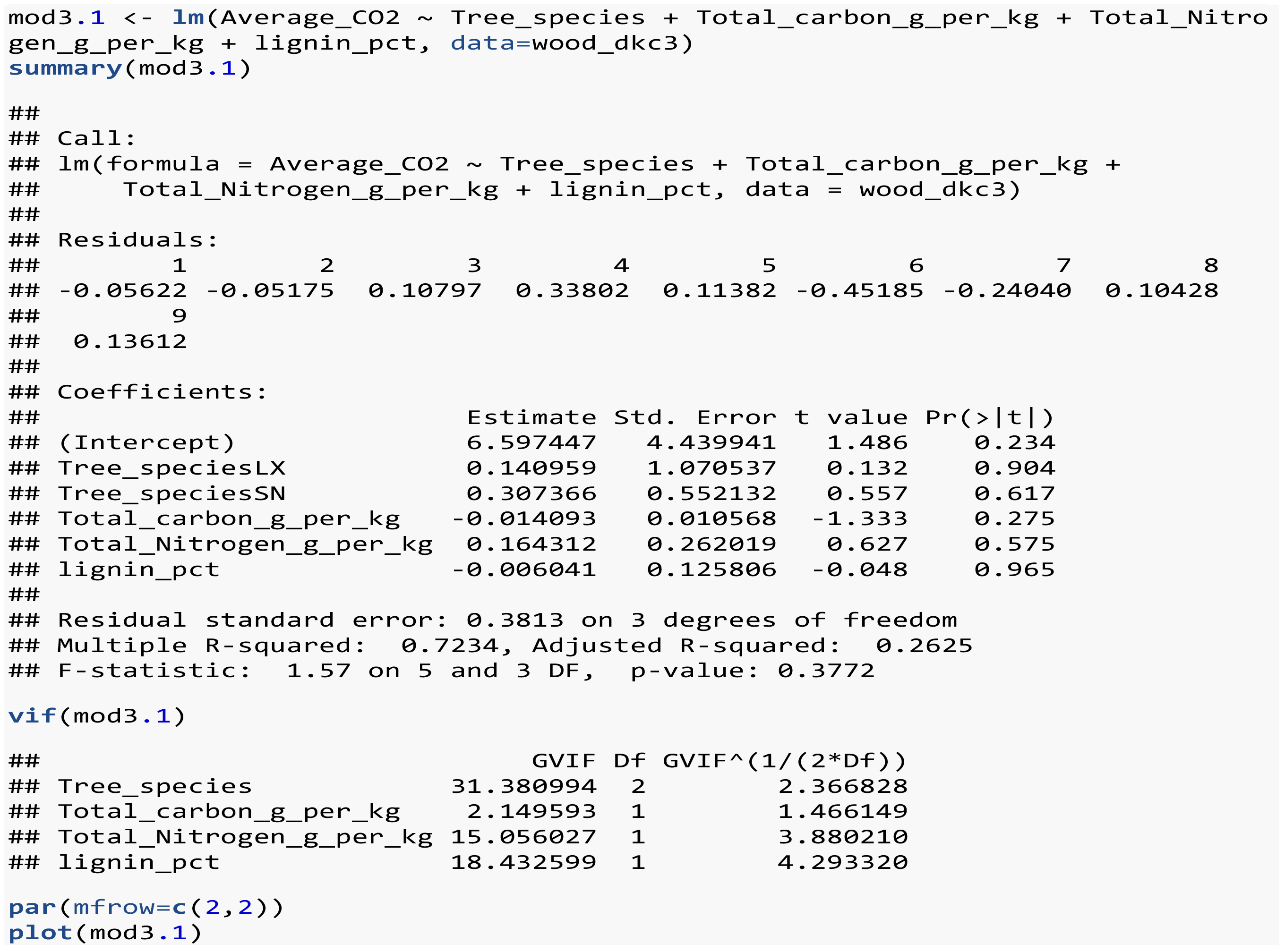

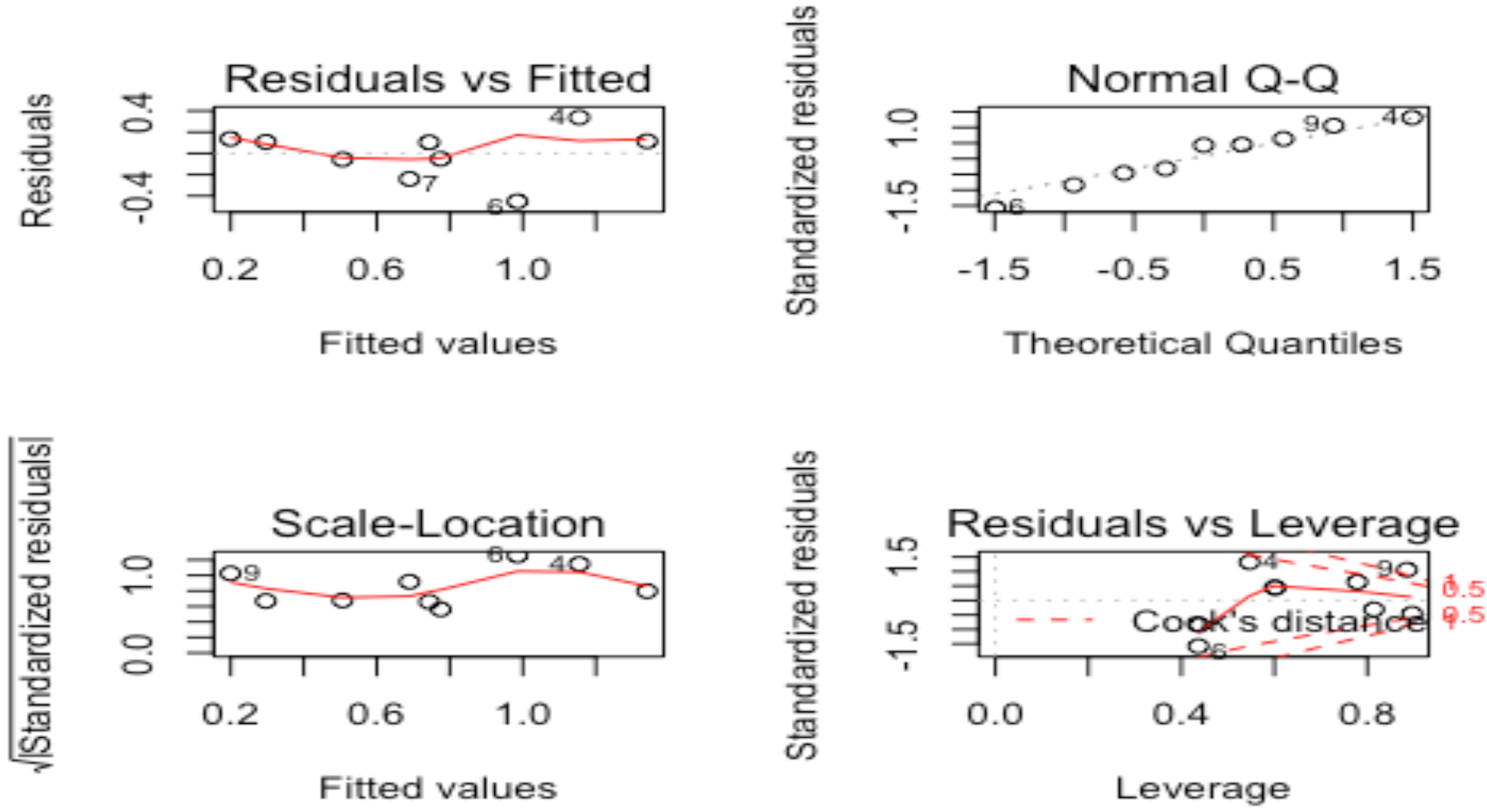

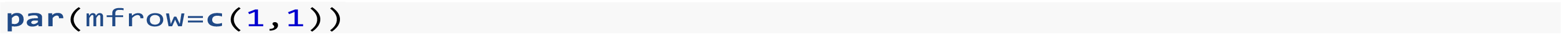

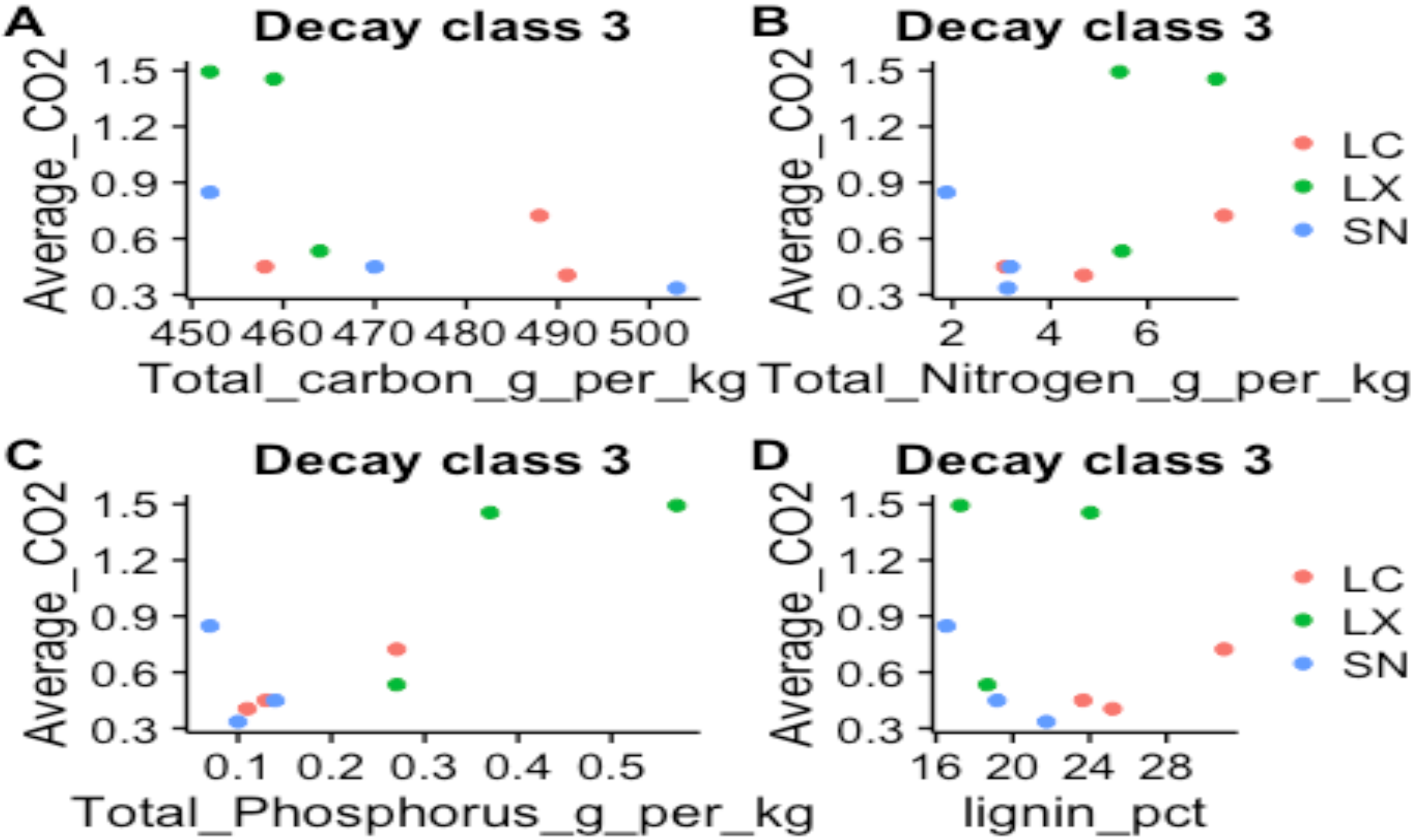

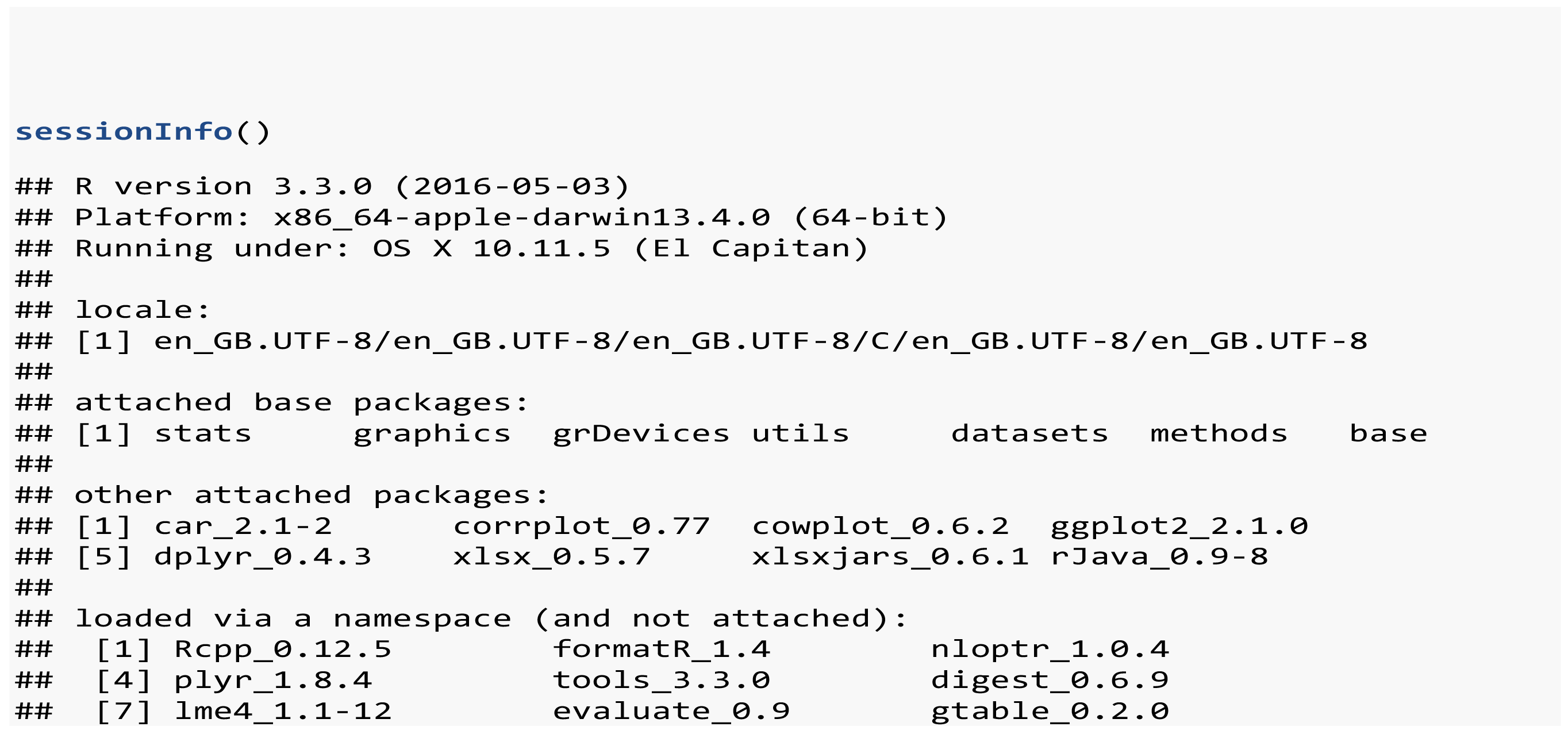

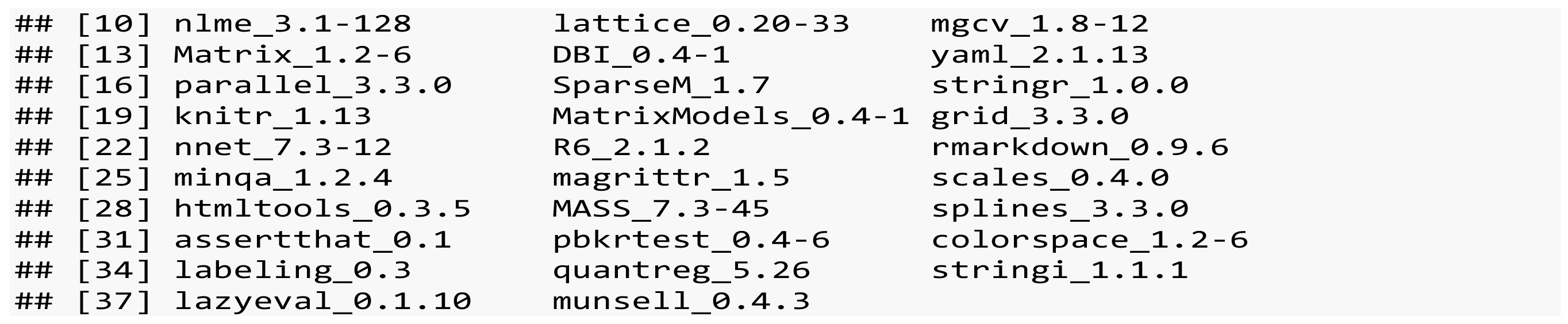

### S3 Sandau et al. analyses

**fungus_Sandau_analysis.R**

***Negorashi2011***

***Thu Oct 30 17:34:54 2014***

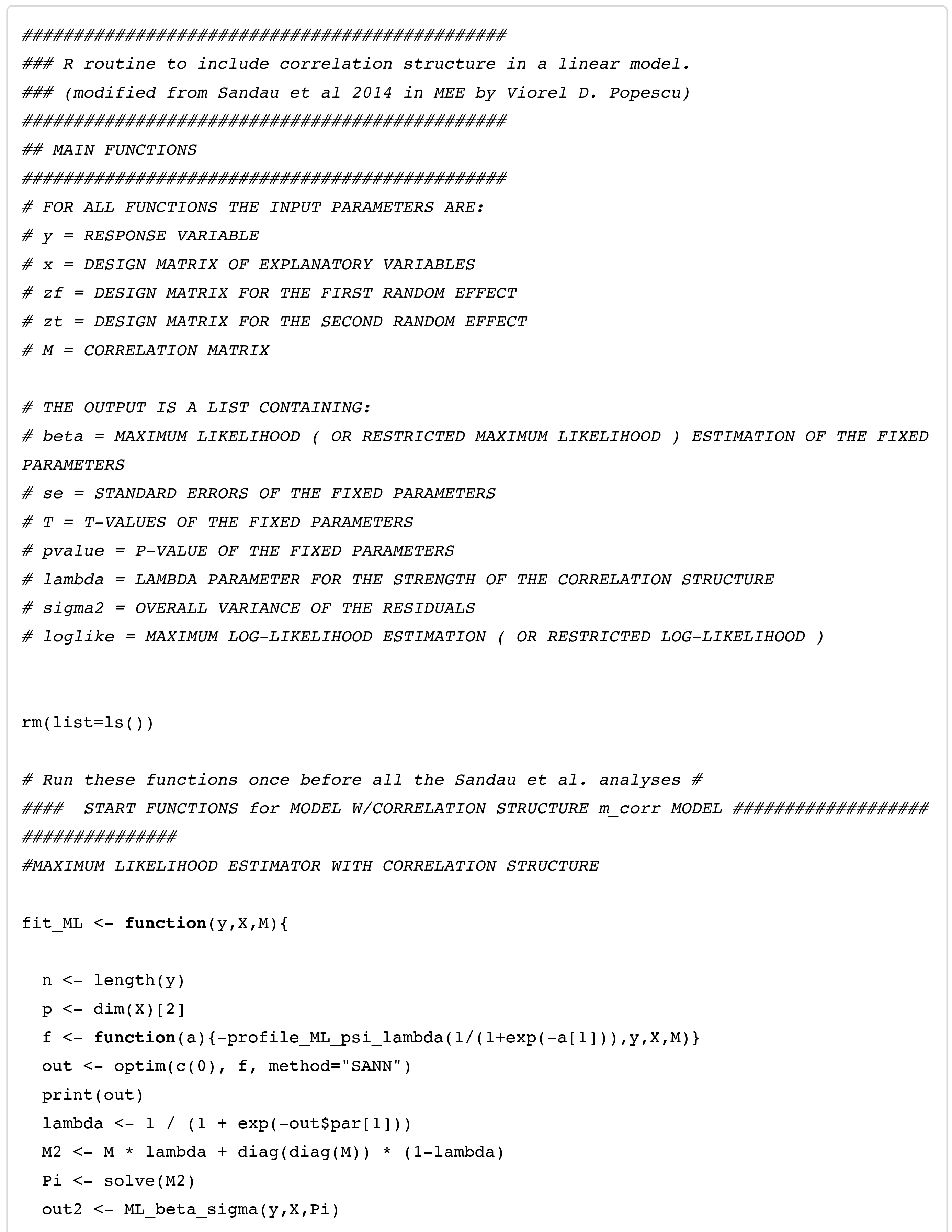

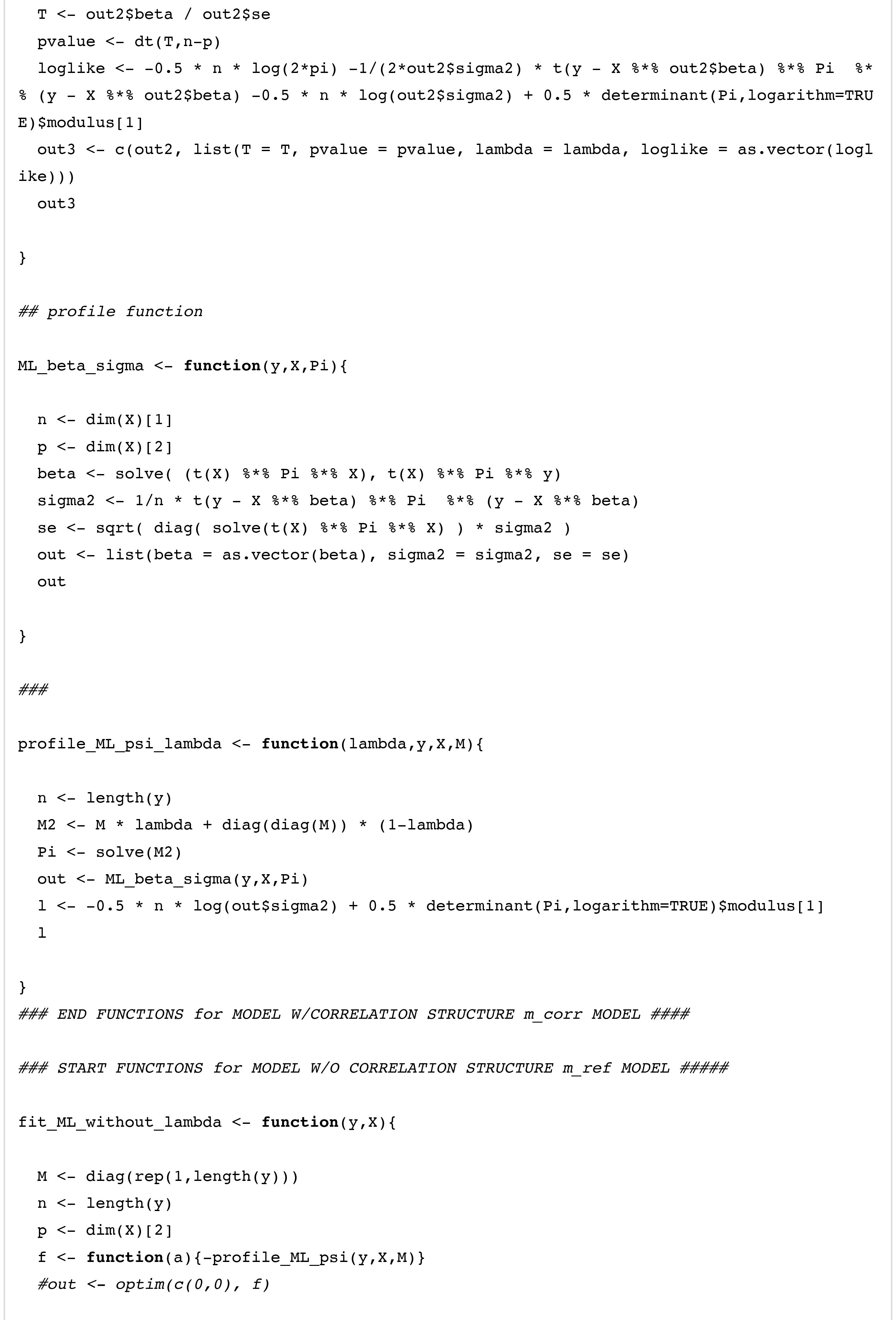

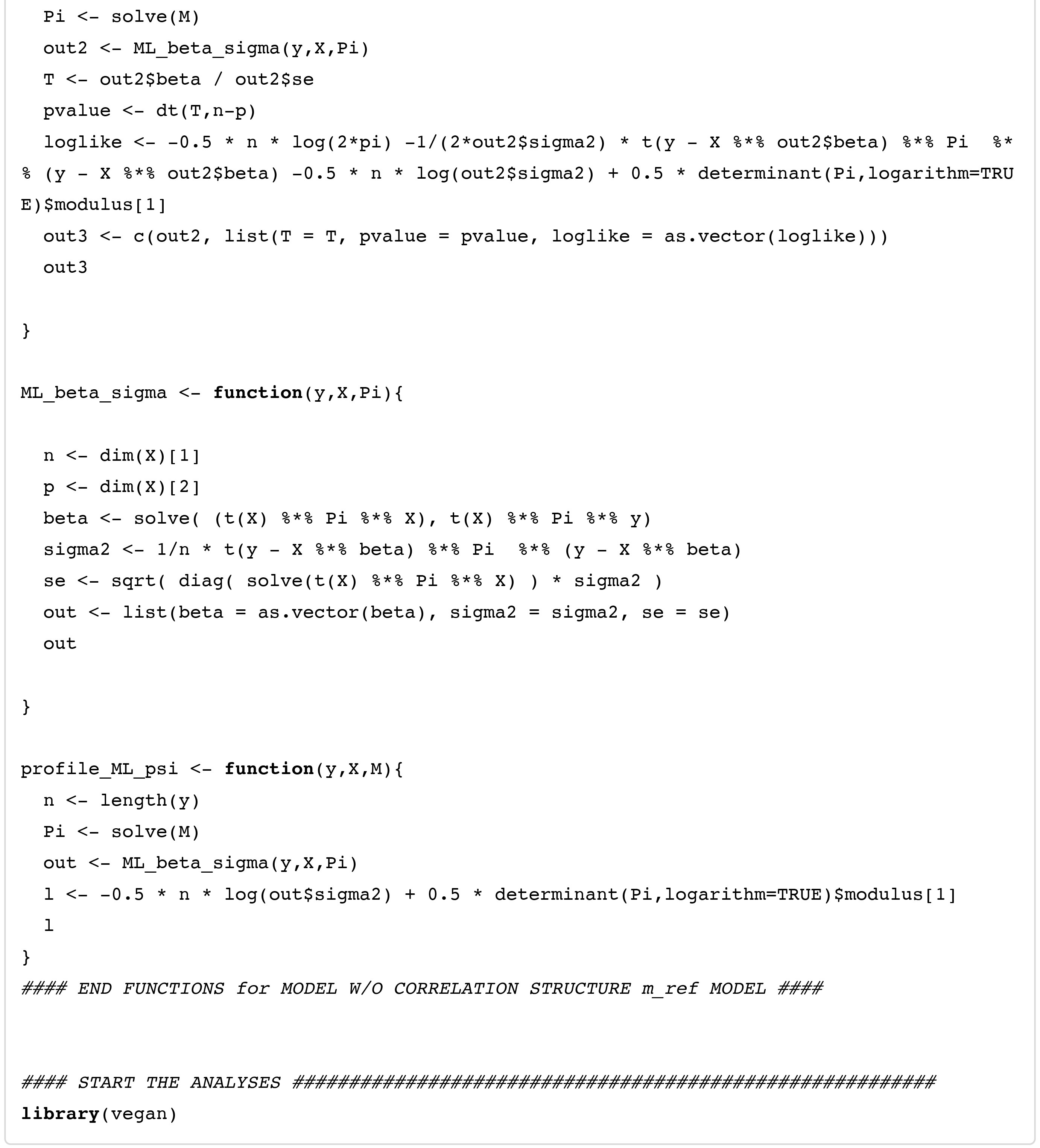

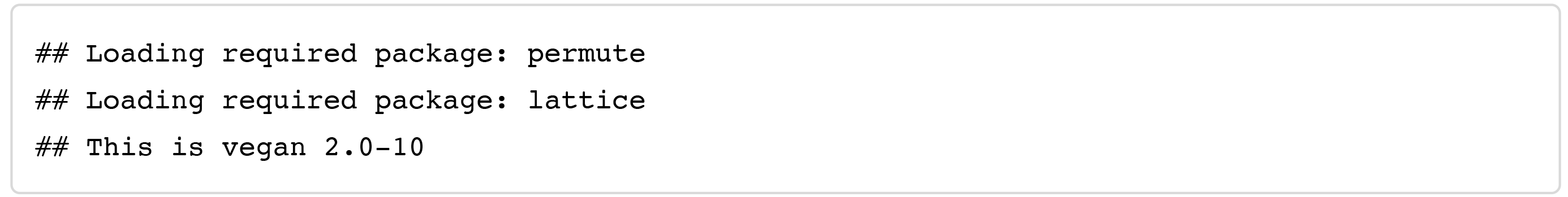

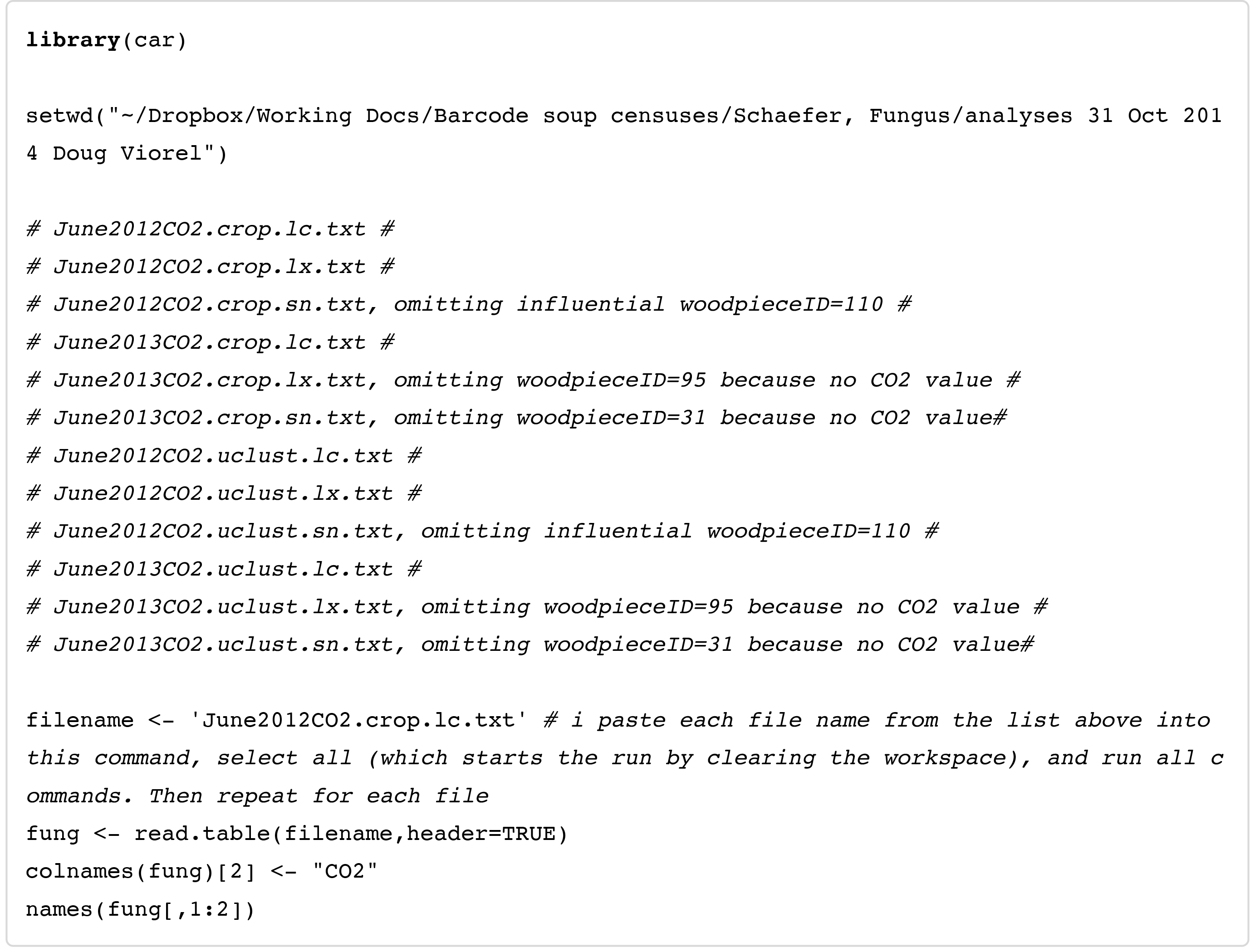

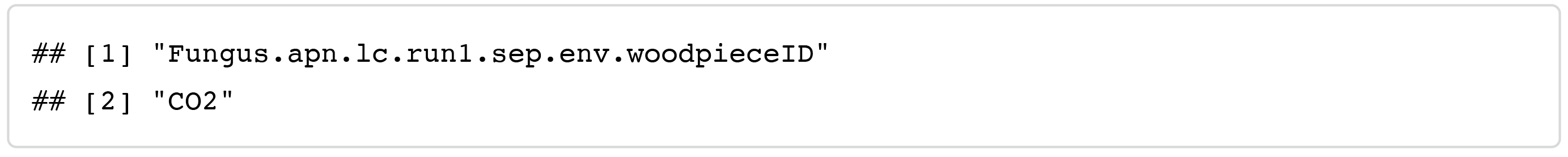

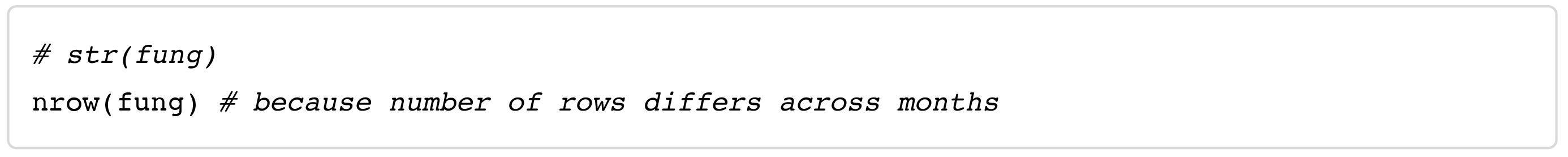

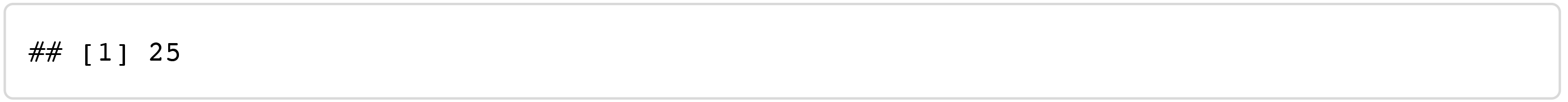

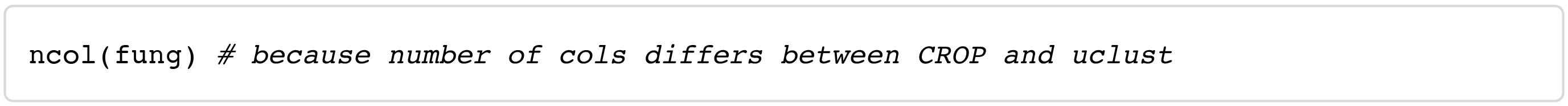

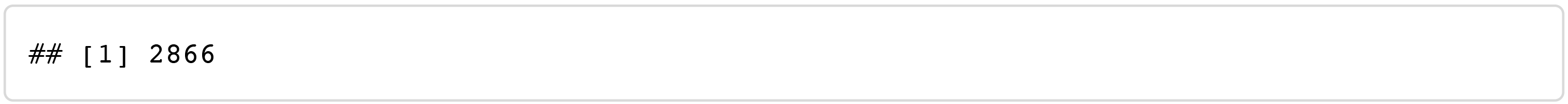

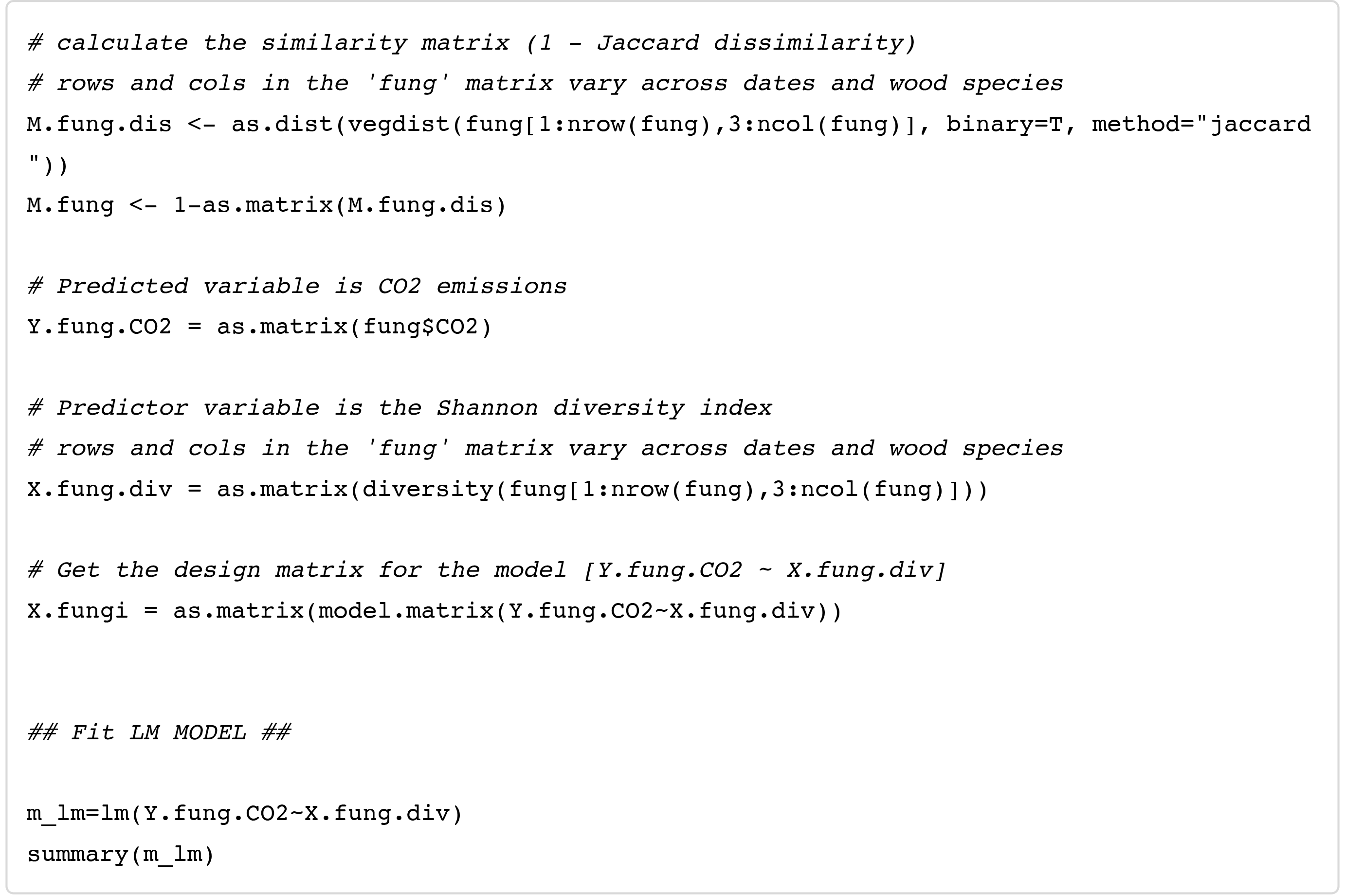

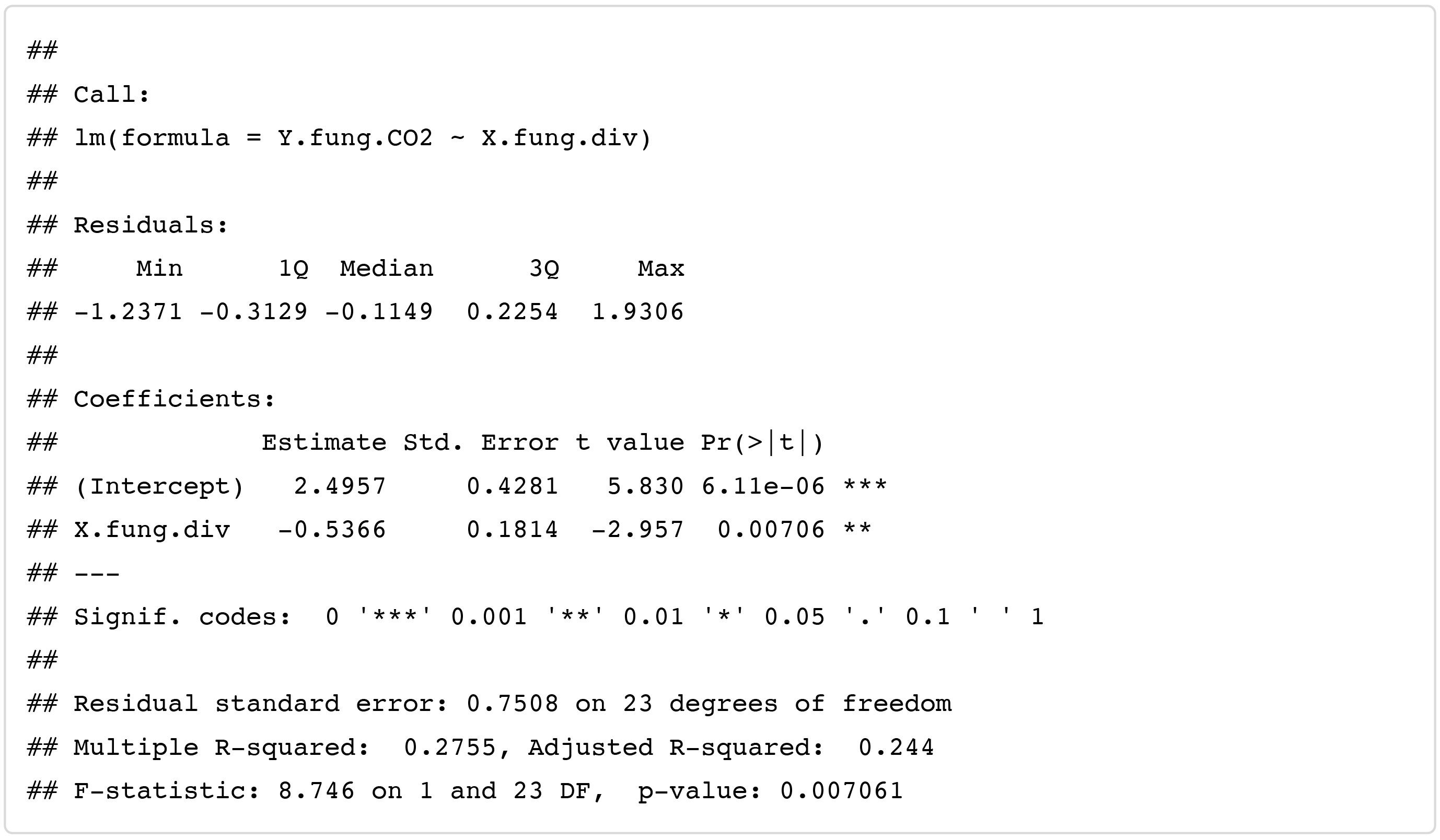

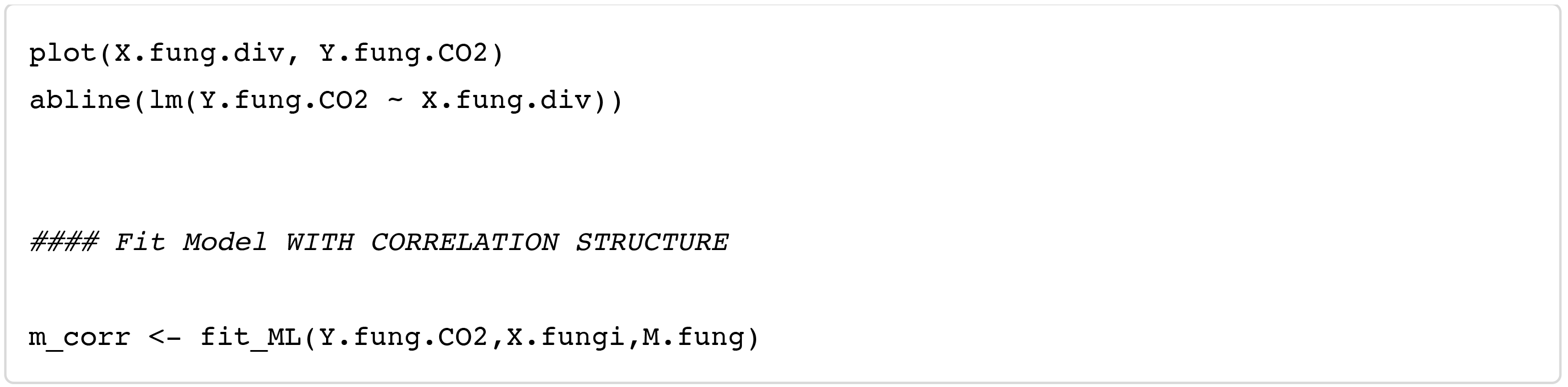

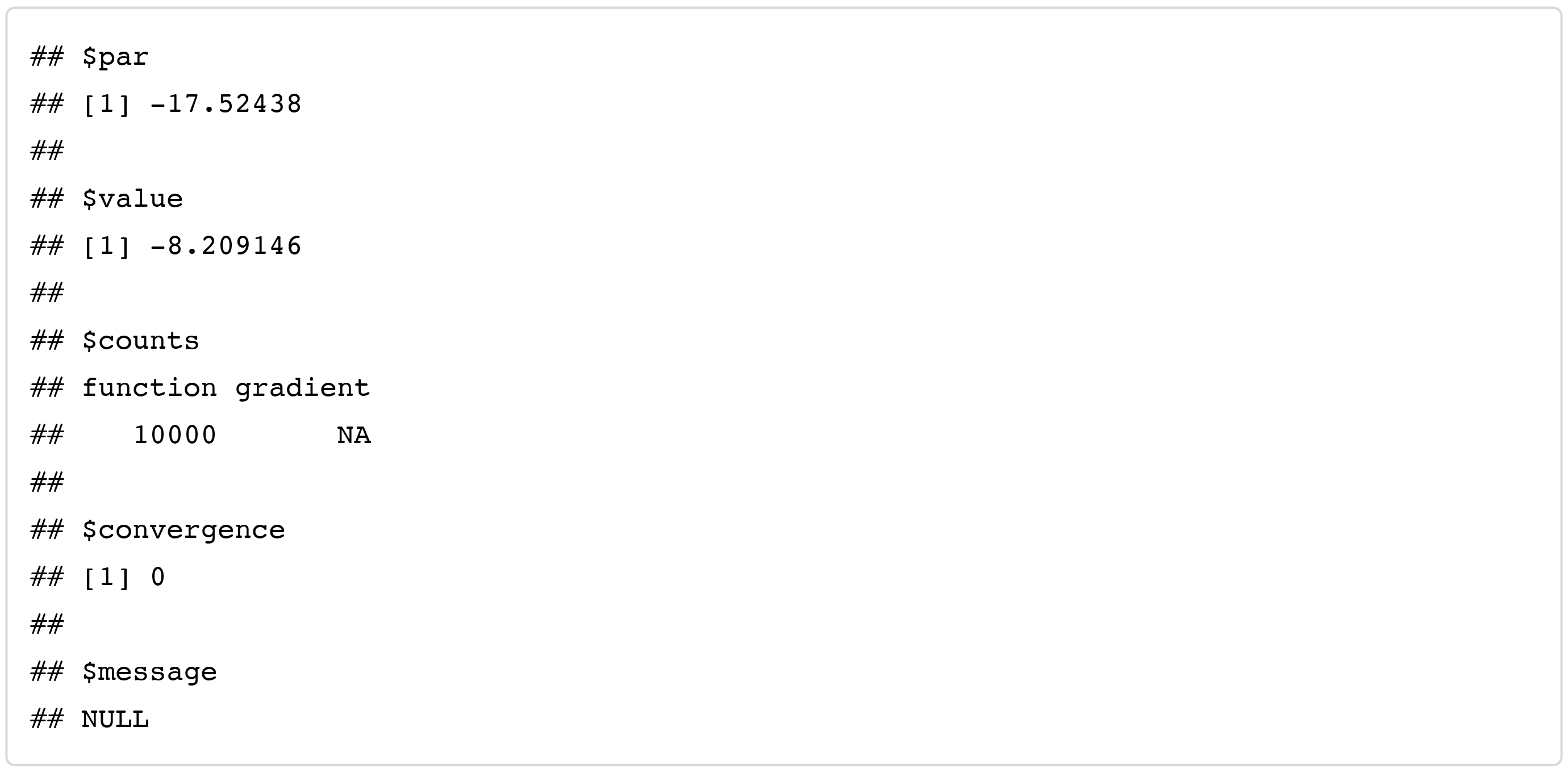

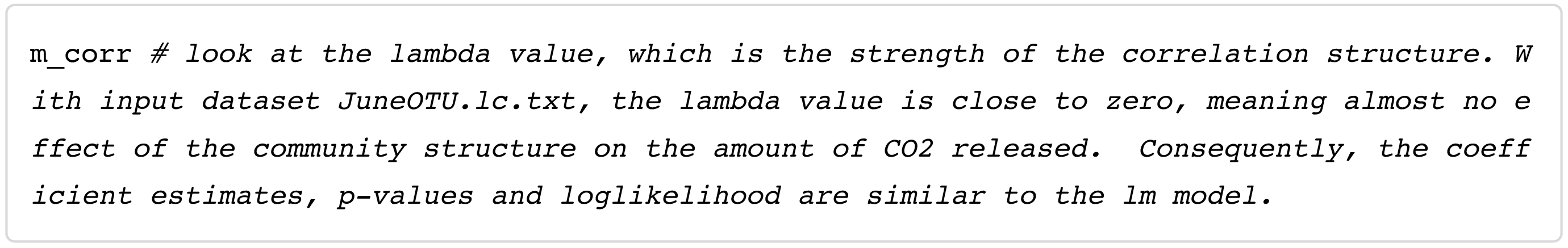

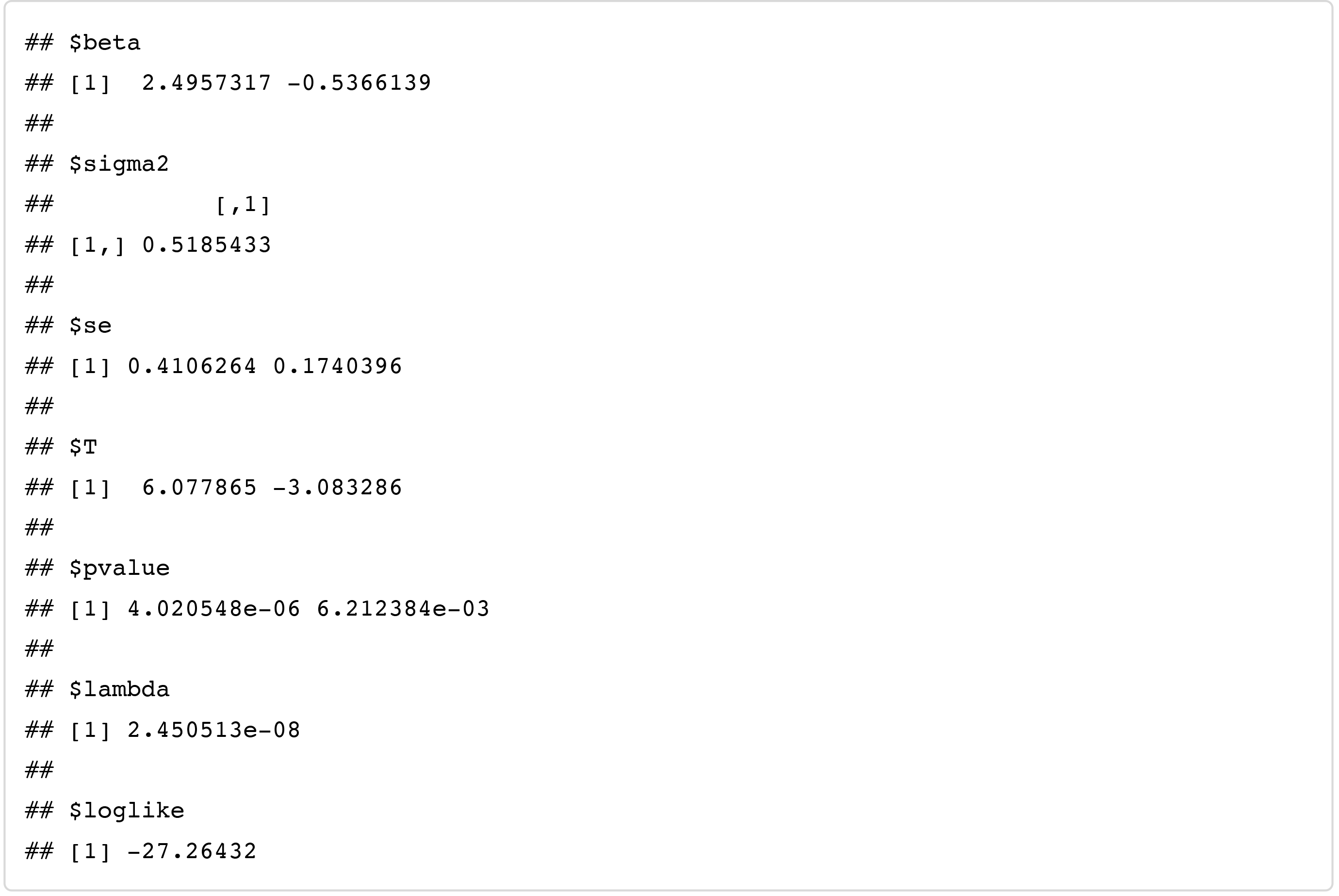

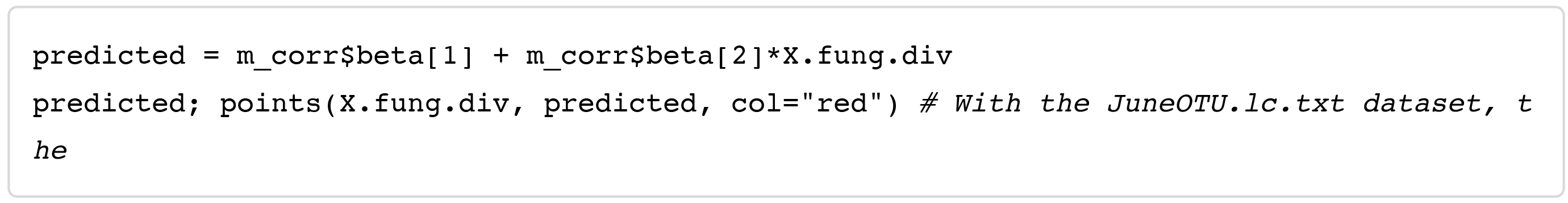

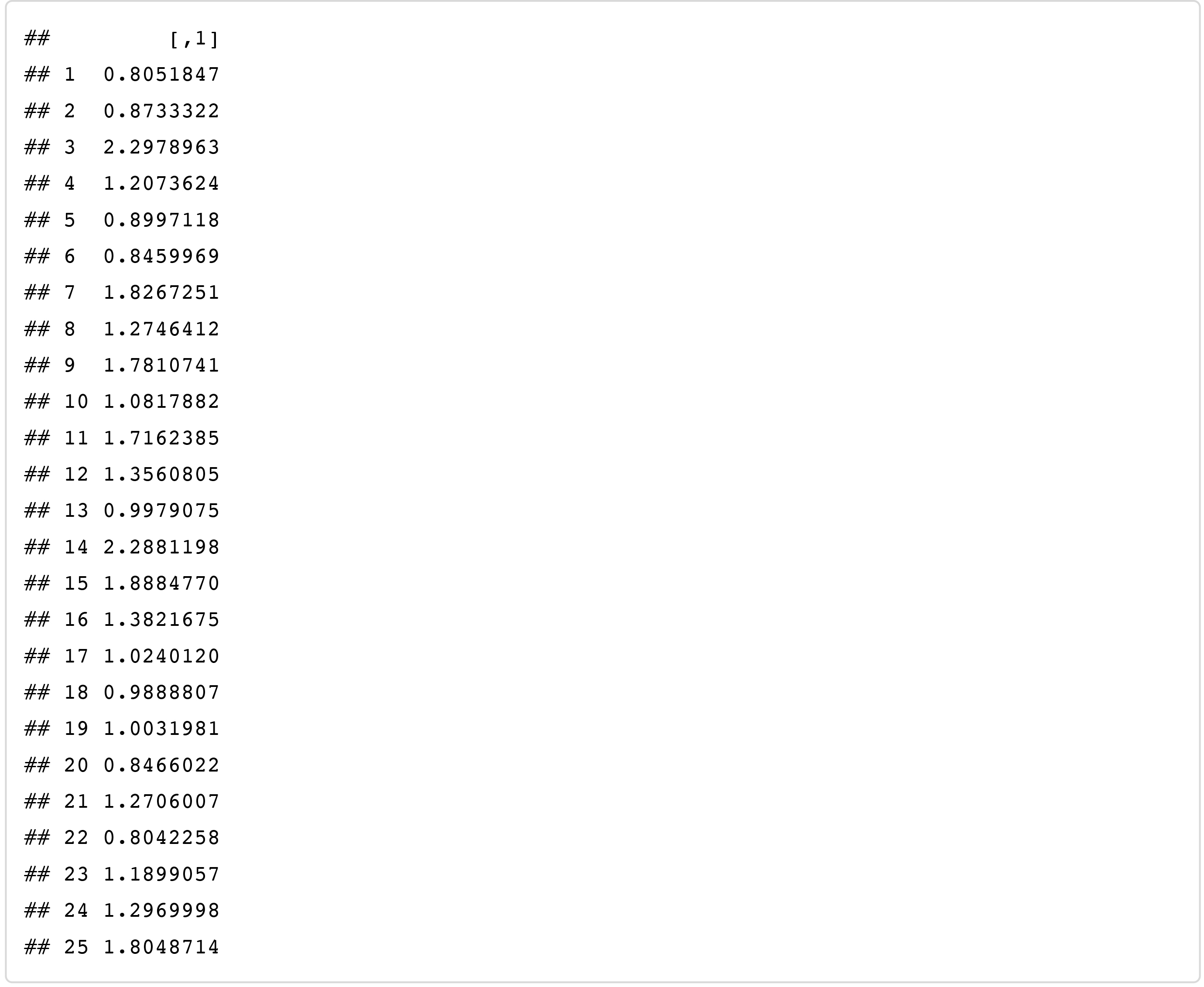

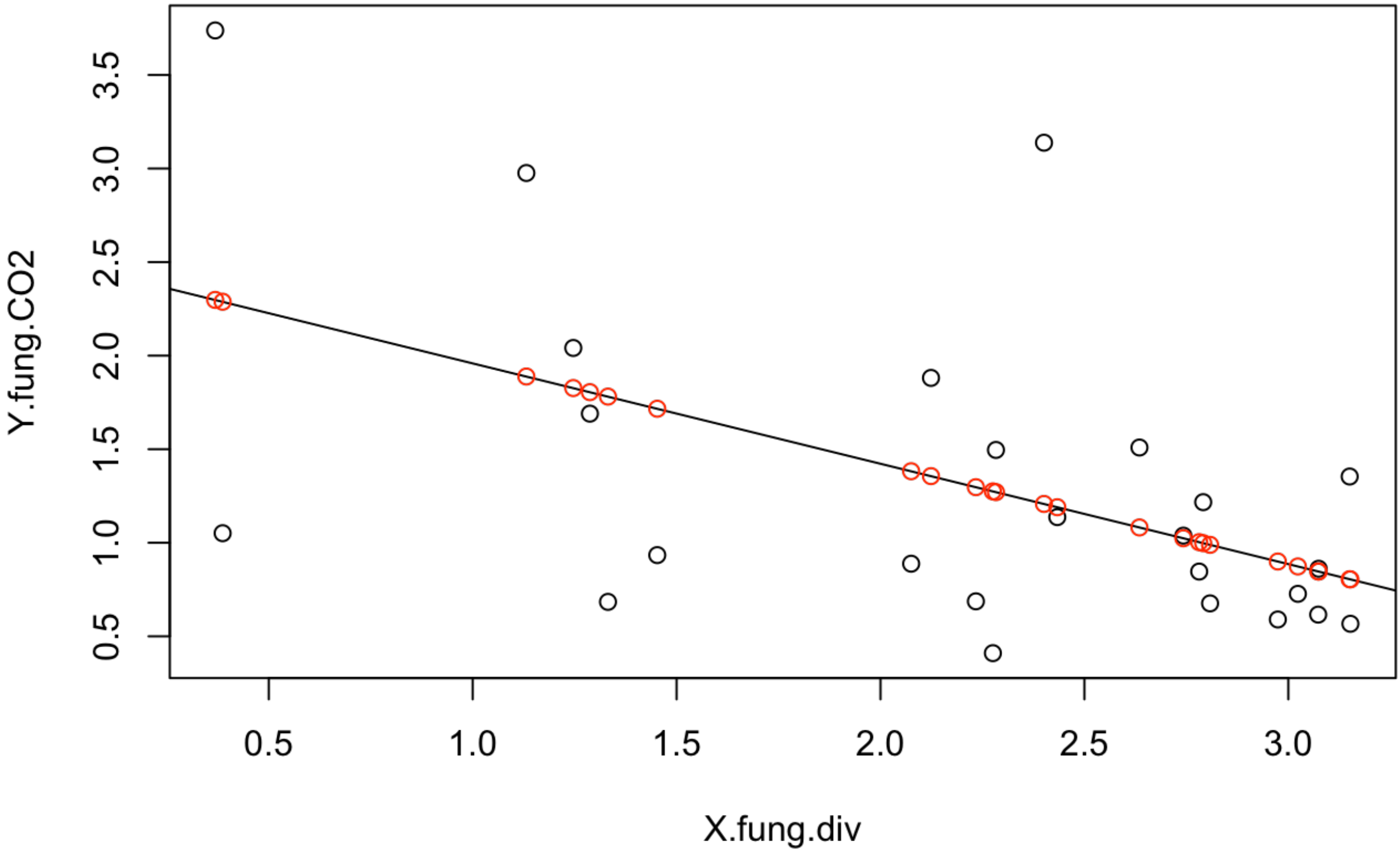

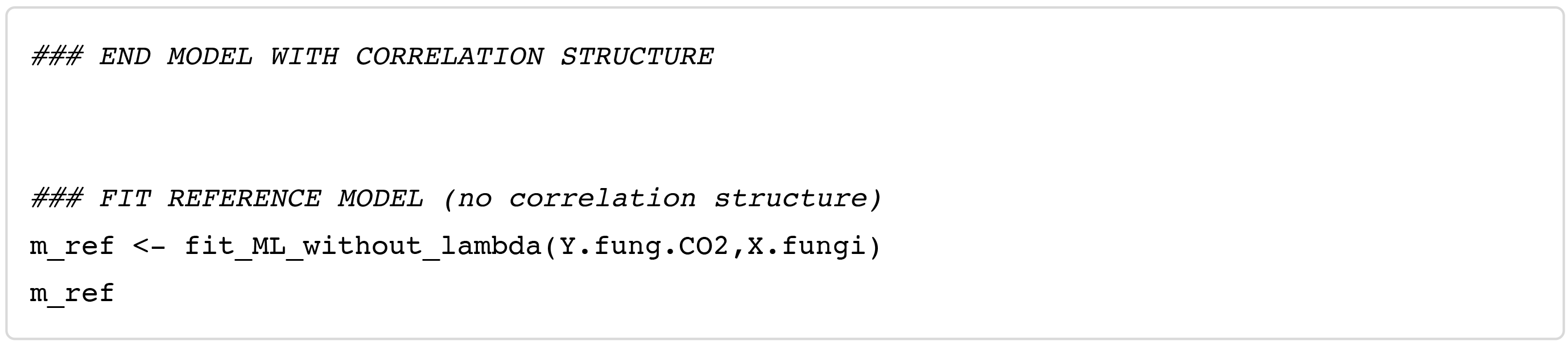

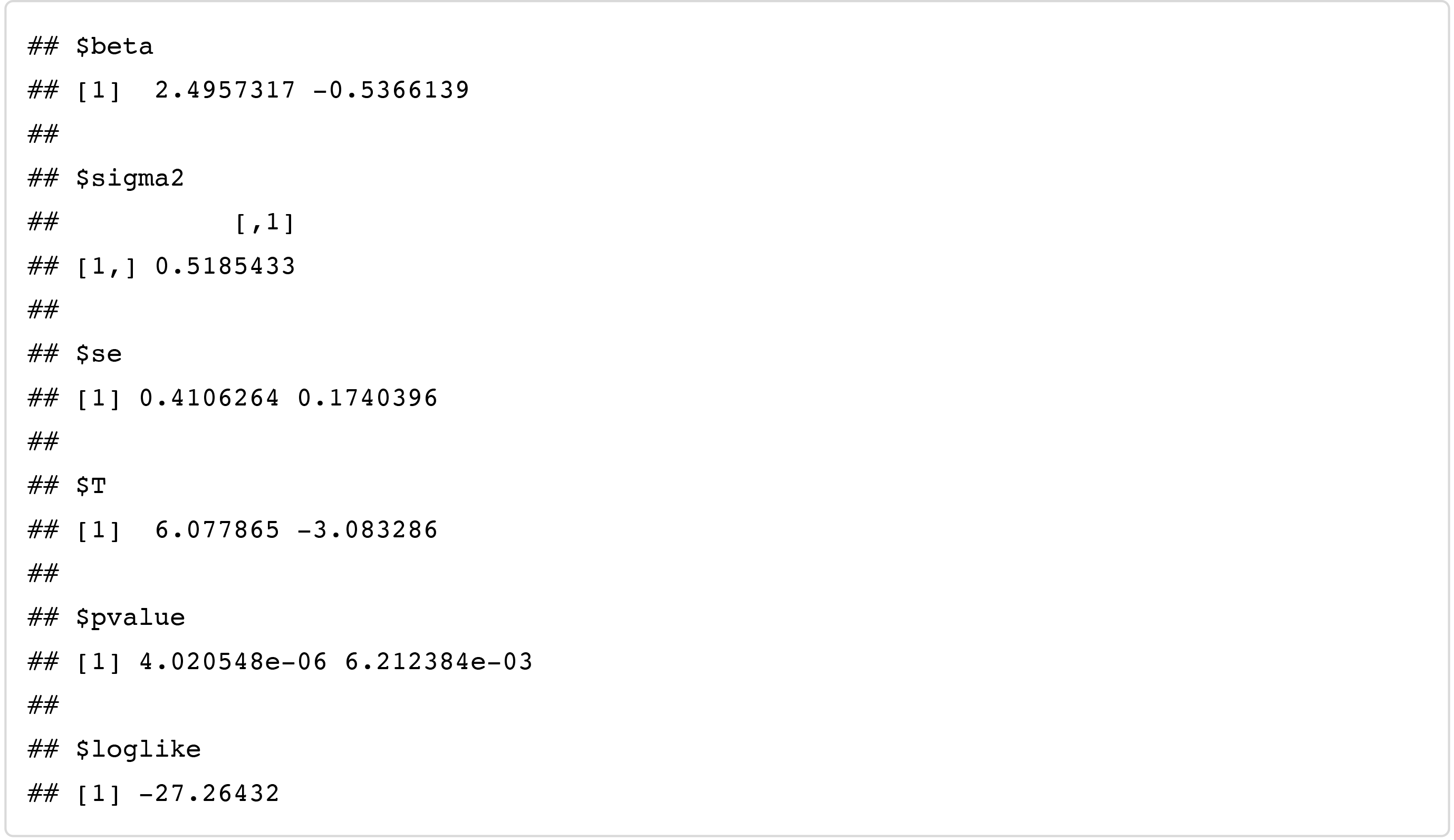

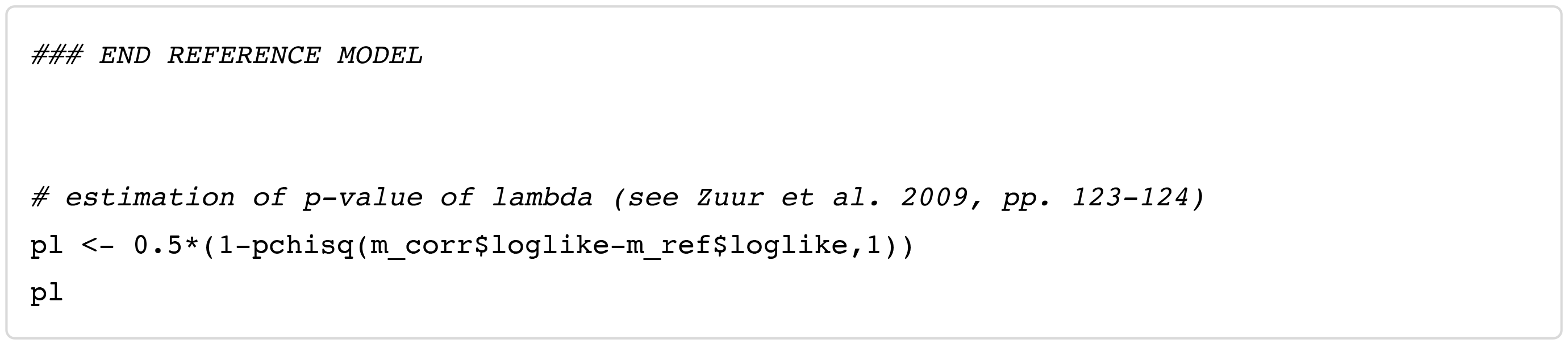

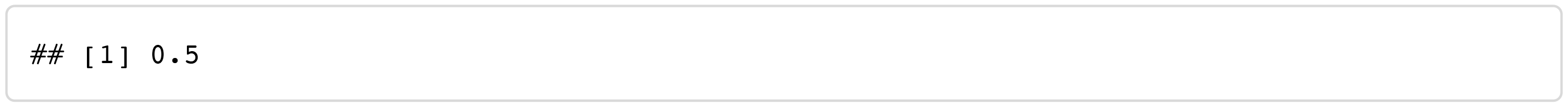

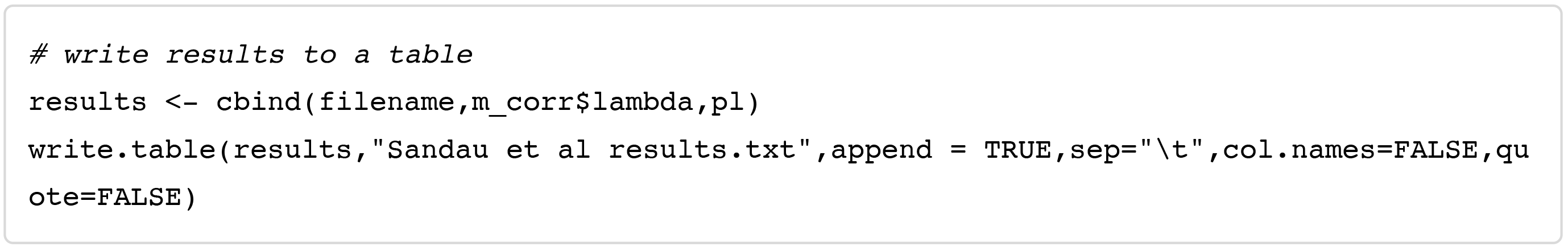

### S4 Conventional Community Analyses

**Supporting Information S4. Conventional community analyses to test for an effect of fungal community composition *per se*on CO_2_ emissions**.

#### Methods

We removed singleton OTUs (OTUs occurring in only one sample) and visualized the OTU tables using correspondence analysis (*cca()* in *R*). We tested the hypothesis that CO_2_ emissions explain variation in community composition by using: (1) *cca*(OTUtable ~ CO_2_) + *anova*(by=“term”, perm=9999) in *vegan (**Oksanen et al., 2013**)* and (2) *manyglm*(OTUtable ~ CO_2_, family=“negative binomial”) + *anova*(resamp=“pit.trap”, nBoot=999) in *mvabund (**Warton et al., 2012**)*. As four versions of each OTU table were tested, we Bonferroni-corrected for multiple tests by multiplying the resulting p-values by 4. *mvabund* is a multivariate implementation of generalized linear models, and unlike dissimilarity-based methods such as *cca, mvabund* does not confound location with dispersion effects, which can inflate both type 1 and 2 errors (*Warton et al., 2012*). Based on the correspondence-analysis plots, we deemed some low-diversity wood pieces to be possible influential outliers (*i.e*. individual samples likely to be largely responsible for any significant community effects), and we removed these data points and repeated the analyses.

#### Results

After Bonferroni correction, no June 2012 samples showed significant effects of fungal composition on CO_2_ emissions, regardless of wood species, statistic, OTU-clustering method, or rarefaction (Table S4).

For June 2013 samples, CCA detected marginally significant effects of composition for all three wood species after Bonferroni correction, but for LC and LX, statistical significance relied on one or two highly dissimilar, low-fungal-diversity wood samples (Main text, Fig. 2). For SN, significant CCA effects were robust to the removal of three potentially influential points, but the *mvabund* test for SN was non-significant after Bonferroni correction. Overall, our data do not reject the null hypothesis that higher fungal species diversity per se, and not any particular fungal species, is responsible for reducing CO_2_ emissions.

**Table S4.** Inferring a ‘pure diversity’ effect of fungal communities on CO_2_ emissions. For each of the six fungal communities visualized in Figure 2, two statistical methods (canonical correspondence analysis and *mvabund*, see Methods), were used to test for significant effects of fungal community composition on CO_2_ emissions. For all June 2012 CO_2_ samples, no significant effects of fungal community (September 2012) were found. For the June 2013 CO_2_ samples, marginally significant effects after Bonferroni correction (p < 0.05) were detected, but only by CCA, not by *mvabund*. Moreover, the significant CCA effects in the two *Lithocarpus* species were not robust to the removal of a few outlying, low-diversity wood pieces, as seen in Fig. 2. Wood species are indicated by LC, LX, SN, for *Lithocarpus chintungensis, L. xylocarpus*, and *Schima noronhae*, respectively.

| CO<sub>2</sub> sample date<sup>1</sup> | Species | Test method | Low-diversity wood pieces removed?<sup>2</sup> | p-value range<sup>3</sup> |
| --- | --- | --- | --- | --- |
| June 2012 | LC | CCA | no, full dataset | 0.10-0.37 |
|  |  |  | 14 removed | 0.13-0.39 |
|  |  | *mvabund* | no, full dataset | 0.15-0.23 |
|  | LX | CCA | no, full dataset | 0.04-0.09 |
|  |  |  | 9, 15 removed | 0.17-0.26 |
|  |  | *mvabund* | no, full dataset | 0.39-0.47 |
|  | SN | CCA | no, full dataset | 0.09-0.15 |
|  |  |  | 7, 10 removed | 0.04-0.24 |
|  |  | *mvabund* | no, full dataset | 0.06-0.54 |
| June 2013 | LC | CCA | no, full dataset | 0.01-0.02* |
|  |  |  | 9, 17 removed | 0.17-0.46 |
|  |  | *mvabund* | no, full dataset | 0.03-0.08 |
|  | LX | CCA | no, full dataset | 0.01-0.03* |
|  |  |  | 9 removed | 0.04-0.18 |
|  |  | *mvabund* | no, full dataset | 0.15-0.32 |
|  | SN | CCA | no, full dataset | 0.01-0.02* |
|  |  |  | 10, 11, 14 removed | 0.01-0.05* |
|  |  | *mvabund* | no, full dataset | 0.03-0.05 |

<sup>2</sup> Numbers refer to points in Fig. 2.

^1^ September 2012 fungal community was tested against June 2012 CO_2_ emissions, as in Fig. 1.

^2^ Numbers refer to points in Fig. 2.

^3^ Asterisks indicate formally significant effects (p<0.05) after Bonferroni correction for four tests: CROP/non-rarefied, CROP/rarefied, *uclust*/non-rarefied, and *uclust*/rarefied. Original p-values reported here.

**S4 A.**
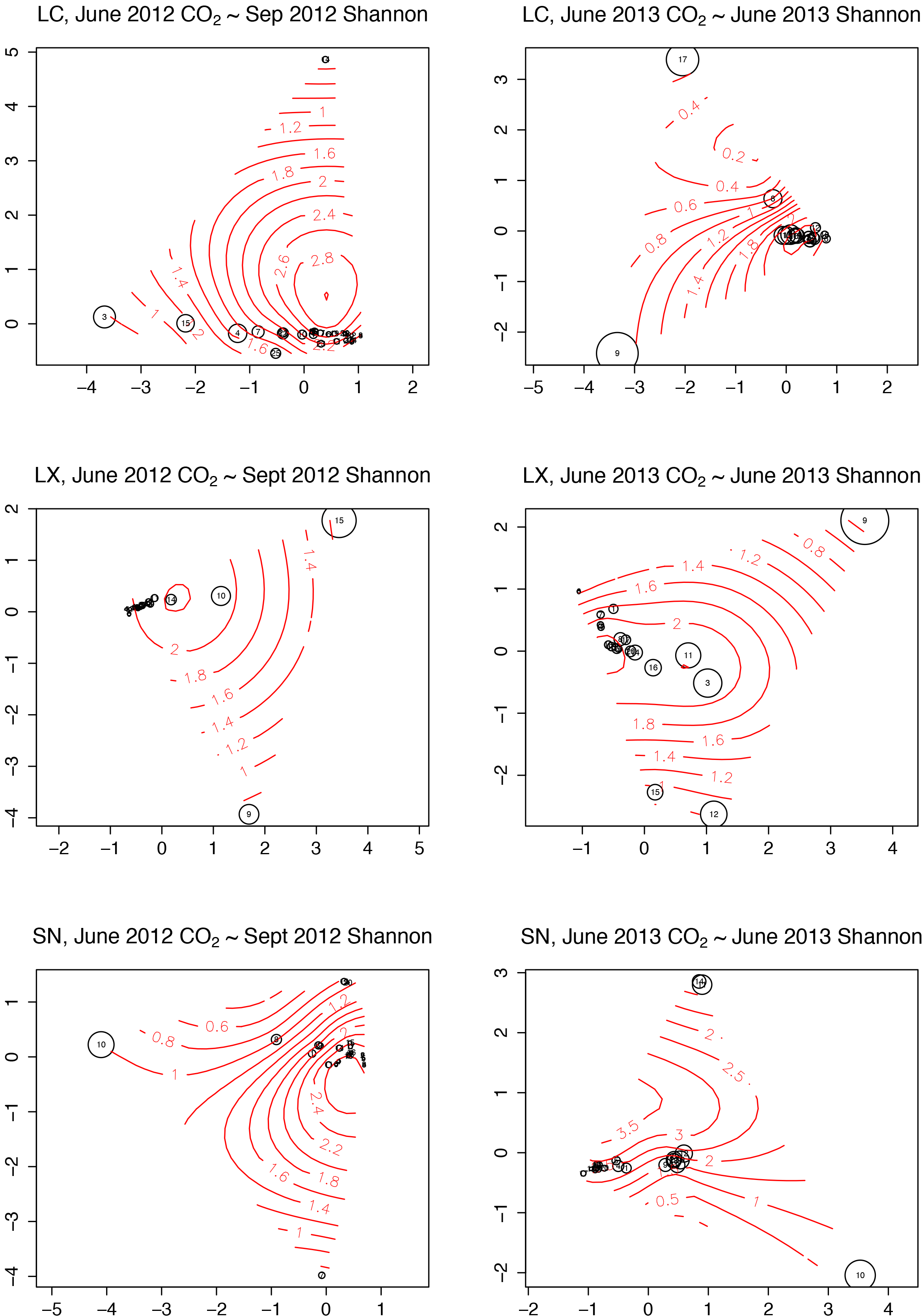
CROP OTU-picking, rarefied

**S4 B.**
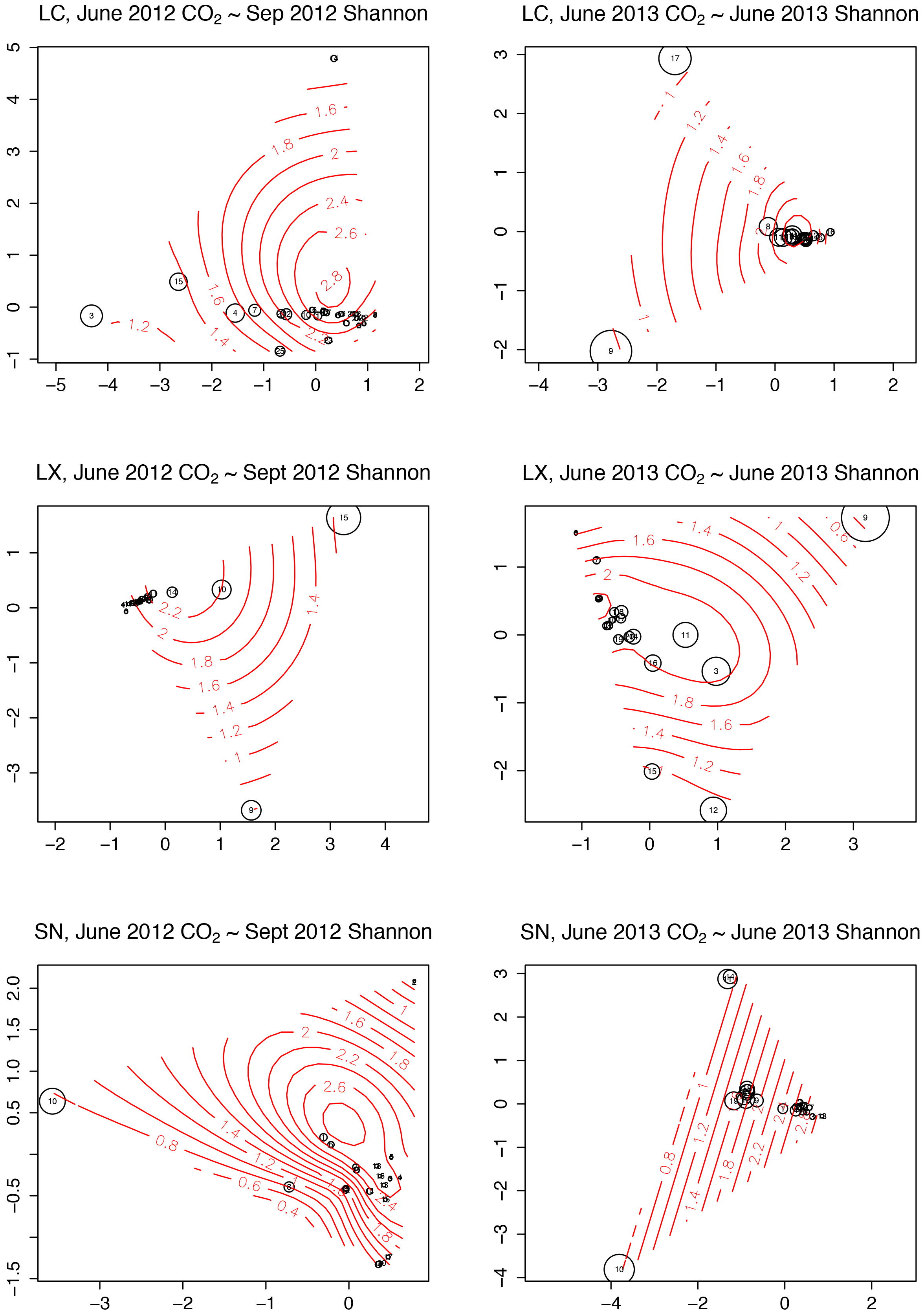
*uclust* OTU-picking, non-rarefied

**S4 C.**
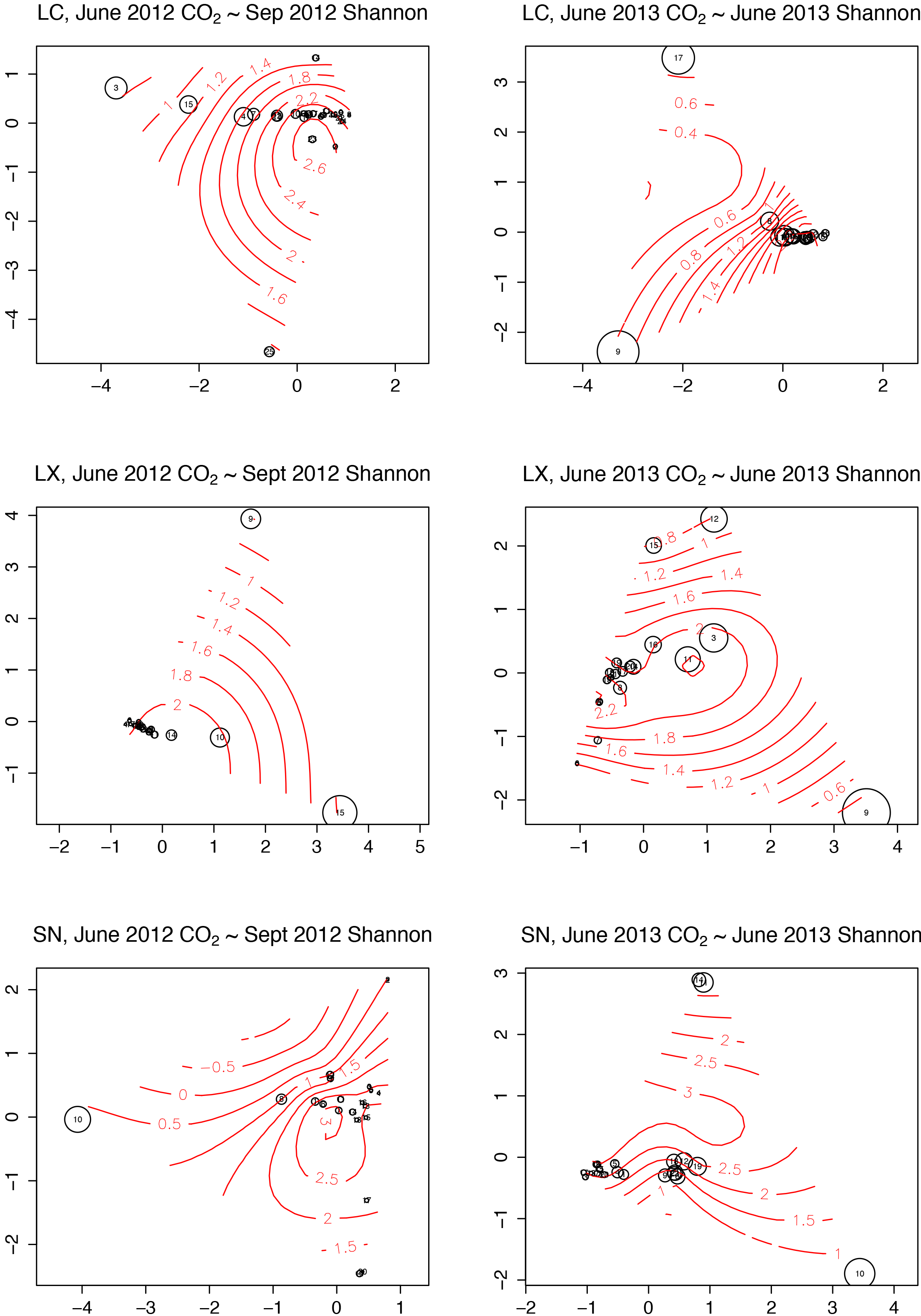
*uclust* OTU-picking, rarefied

### S5 Gravimetric vs CO_2_ estimated loss

**Supporting Information S5.**
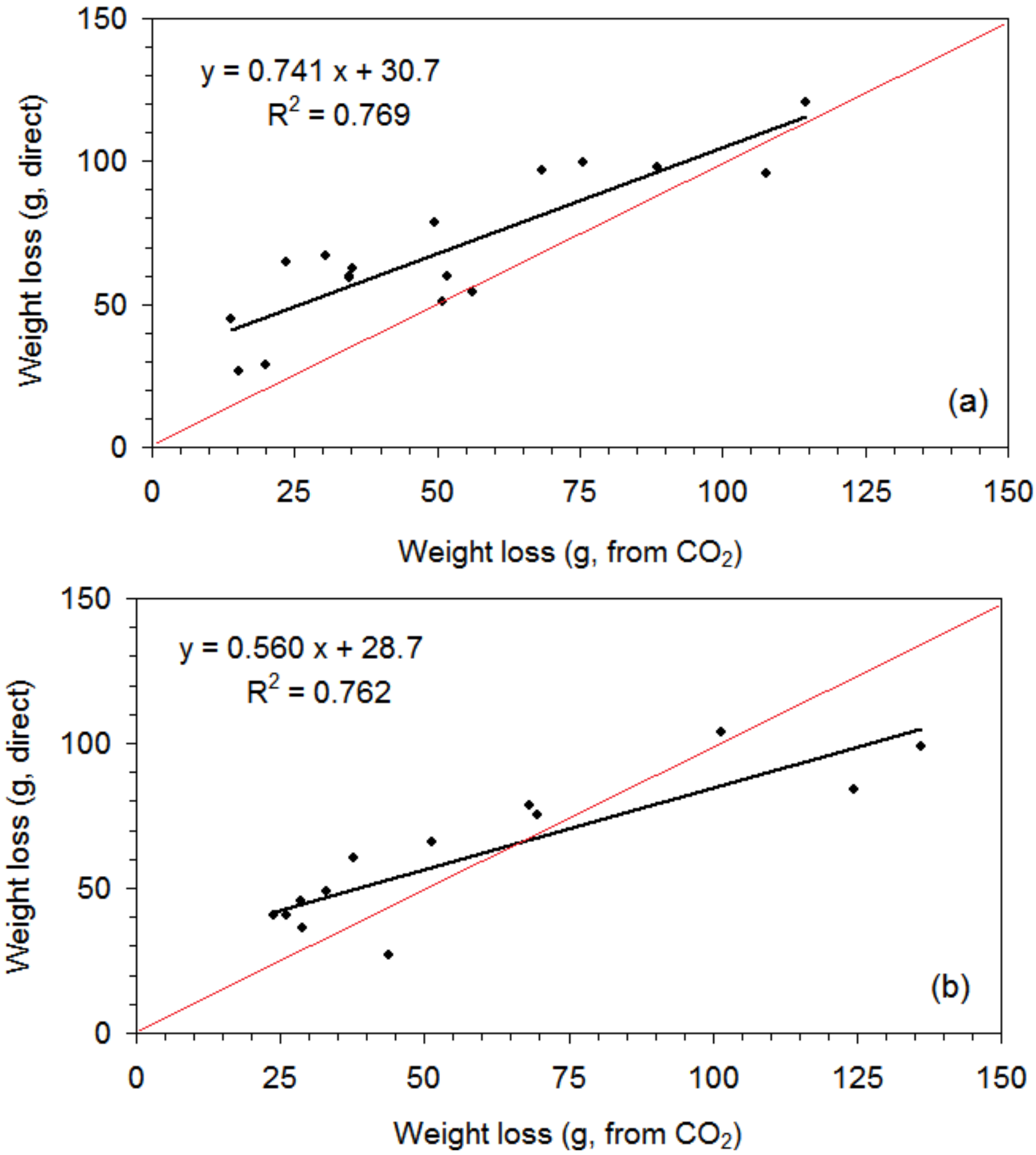
Mass loss from wood pieces (g over 3 years) as measured gravimetrically and as estimated from average CO_2_ emission rates in decay class 1 (DKC1, top), decay class 2 (middle), and decay class 3 (bottom). Black lines are linear regressions and red lines represent 1:1 correspondences.

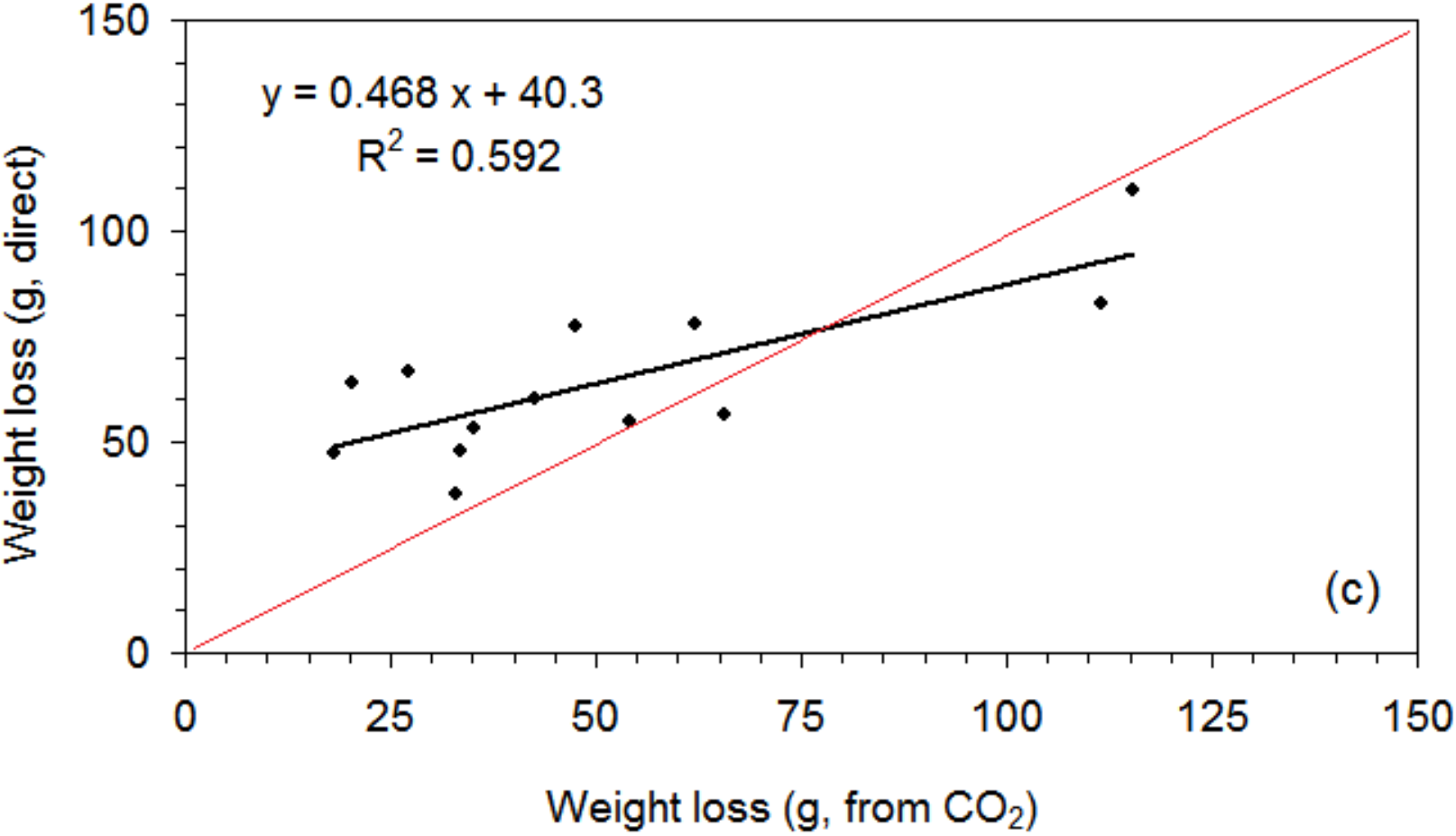

### SI Bioinformatic Command Scripts

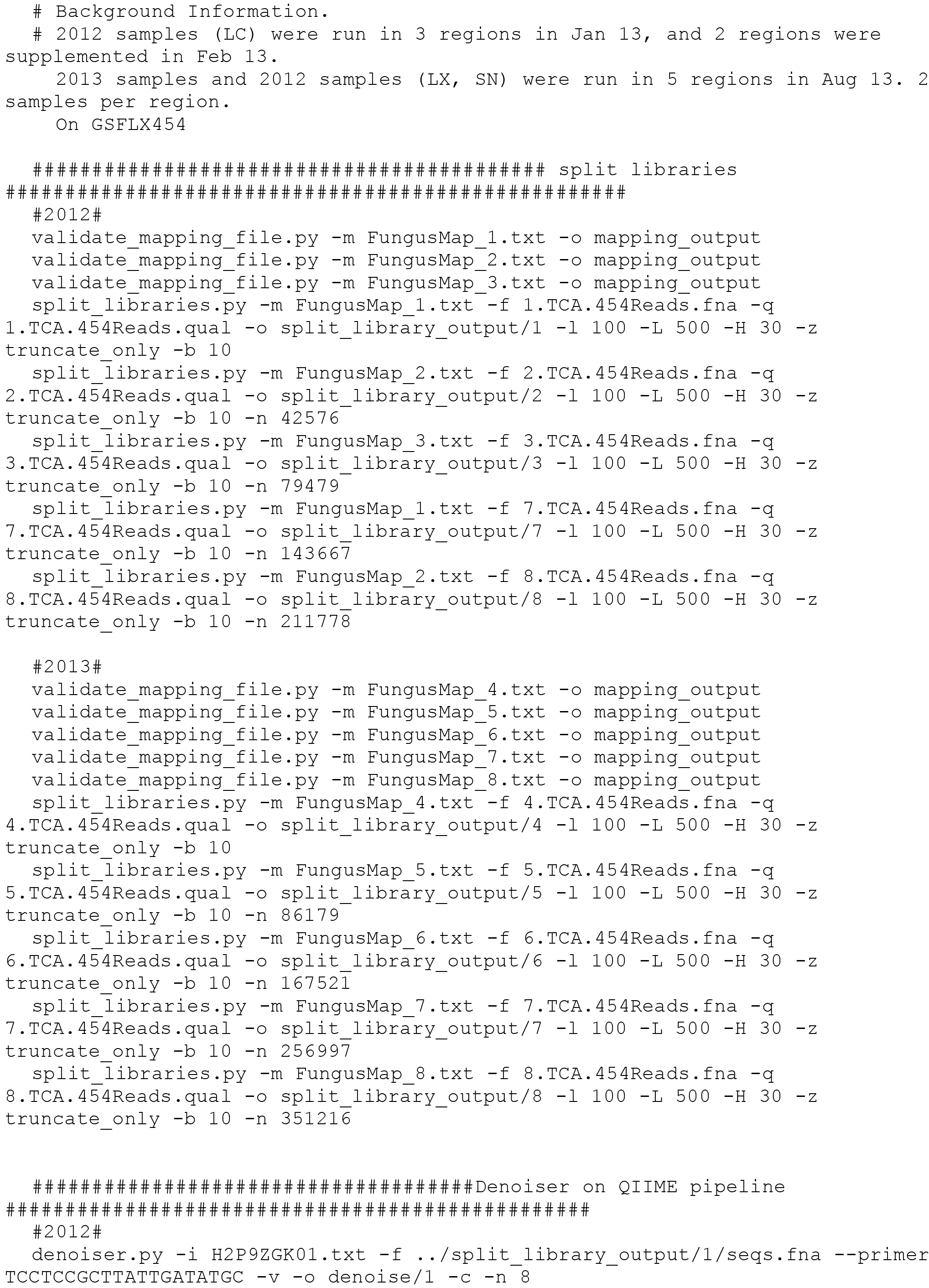

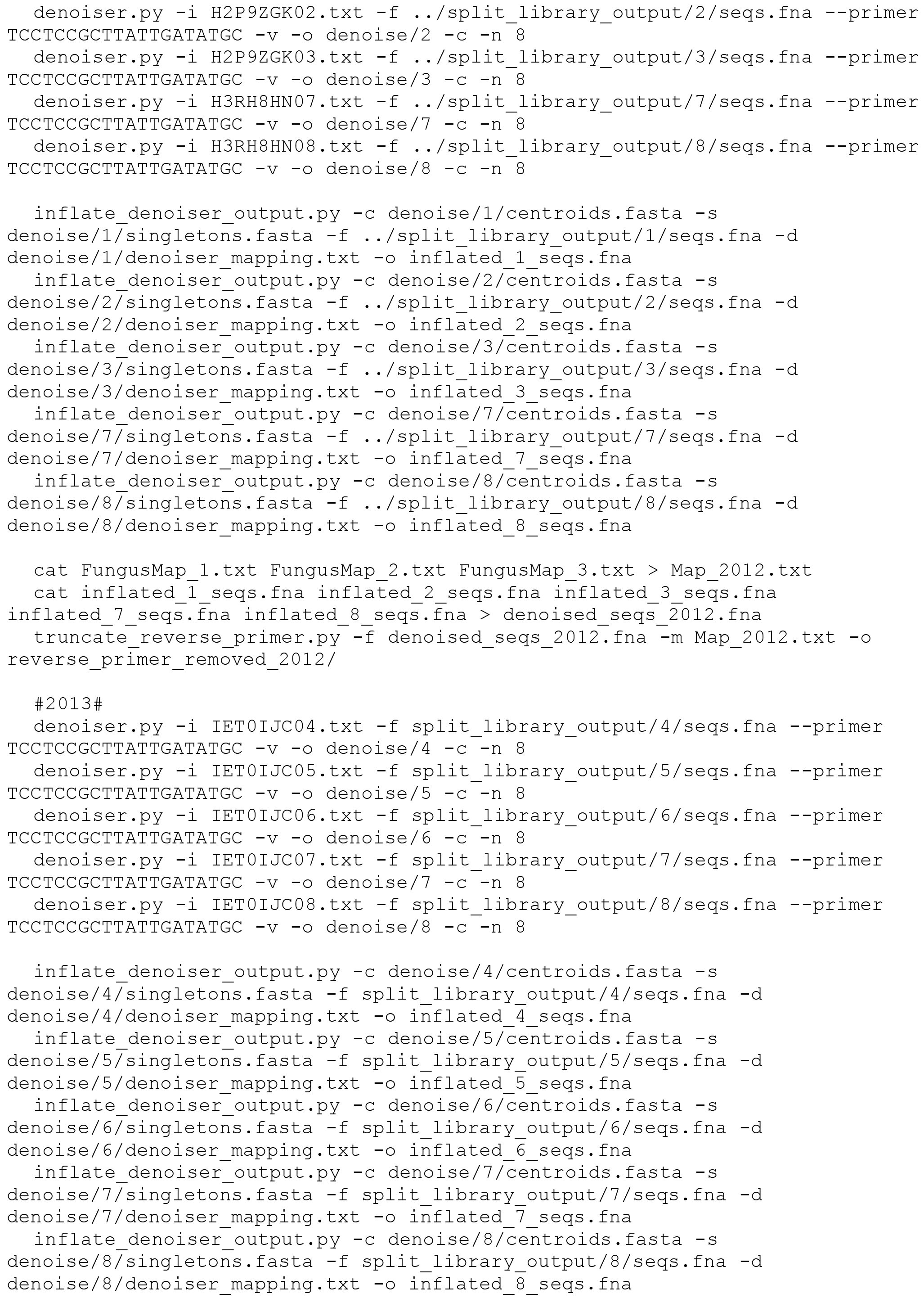

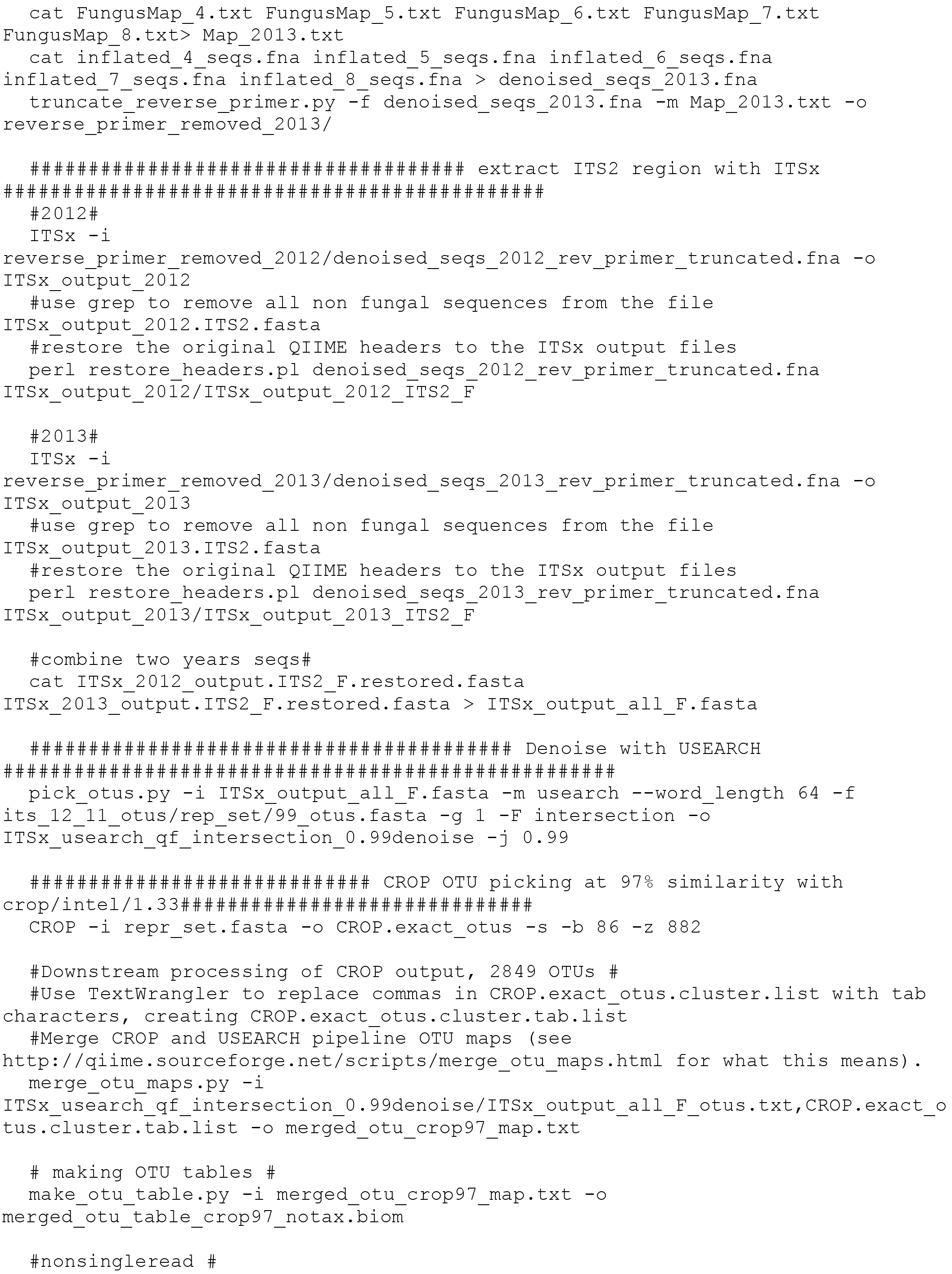

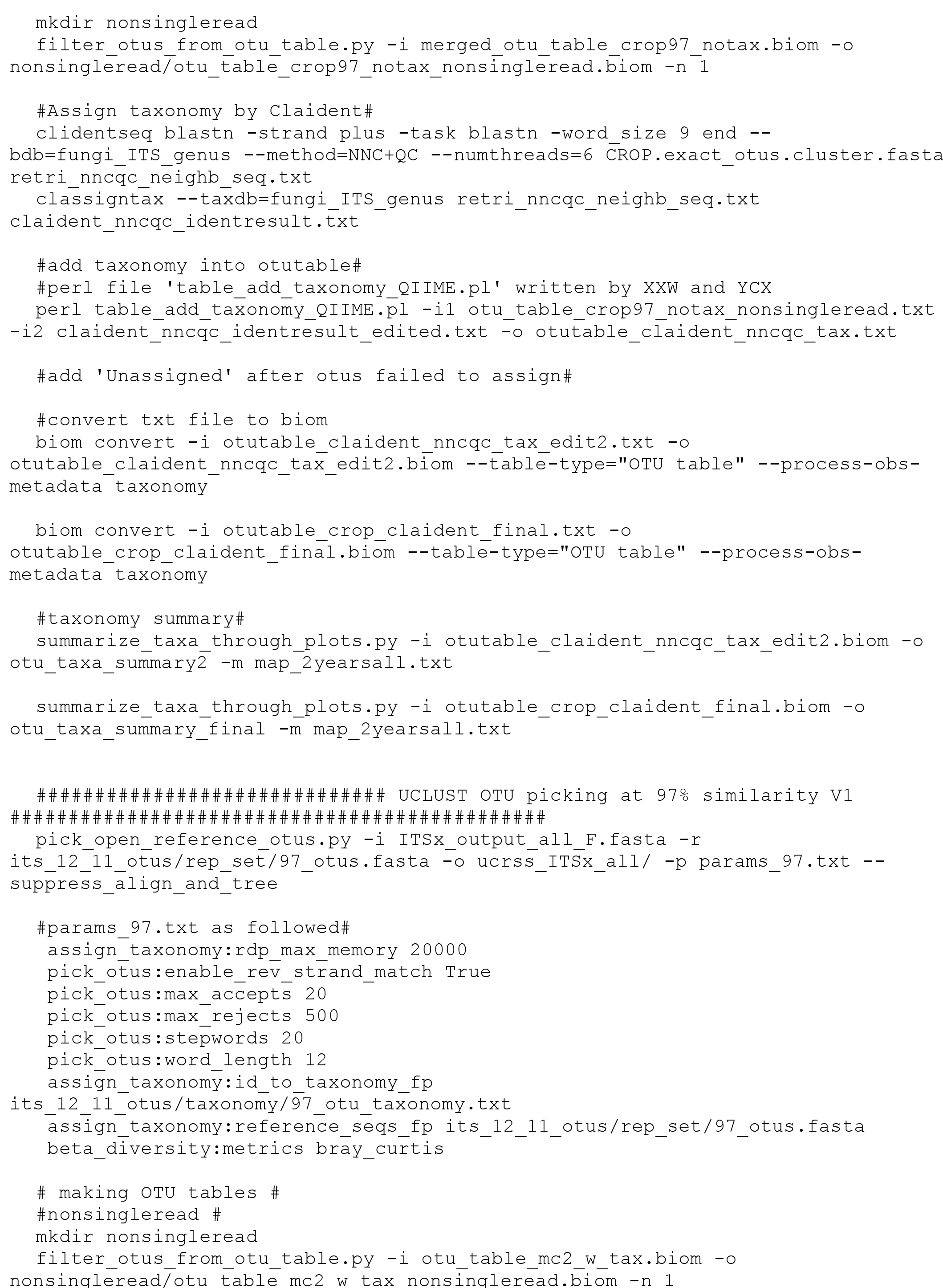

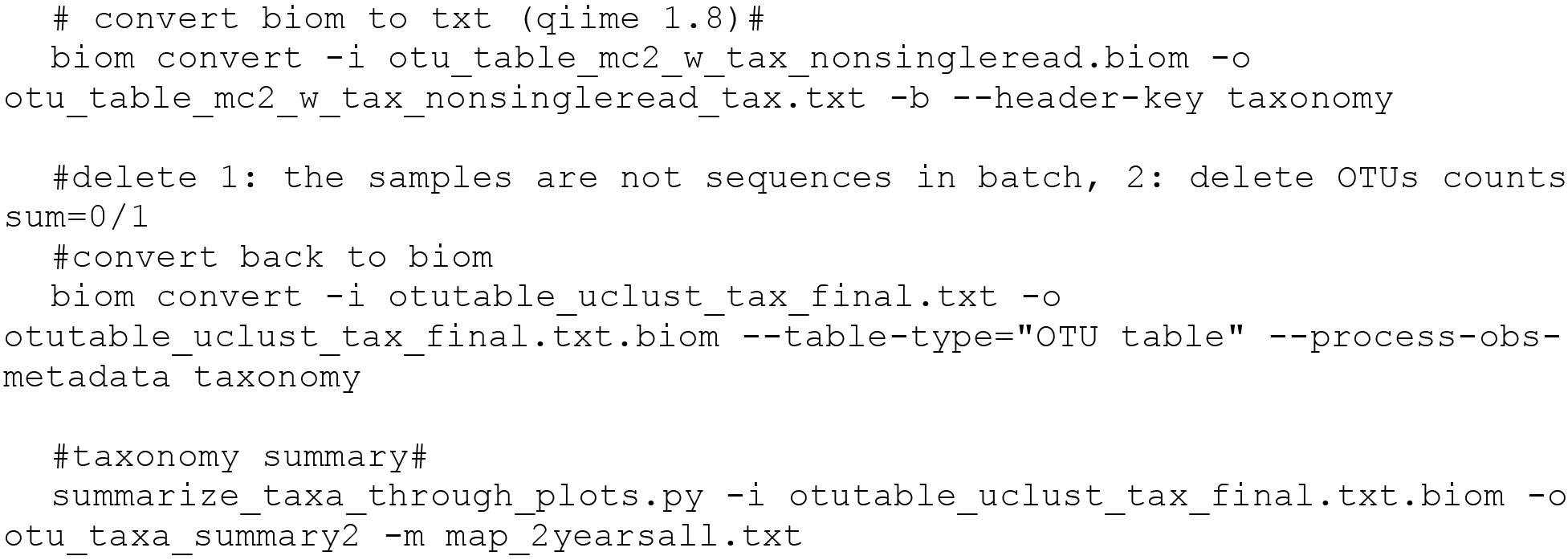

